# Blocking Minor Intron Splicing Disrupts DNA Repair and Overcomes Therapy Resistance in Prostate and Breast Cancer

**DOI:** 10.64898/2025.12.10.693392

**Authors:** Anke Augspach, Paola Francica, Kaitlin Girardini, Muriel Jaquet, Marika Lehner, Antonio Rodriguez-Calero, Charlotte Komarek, Martín González-Fernández, Hannah L. Williams, Xiaoyue Deng, Lea Lingg, Cristina Zivko, Vera Fuchs, Seynabou Diop, Sina Maletti, Simone de Brot, Alexander Ewe, Elena Perugini, Nigel B. Jamieson, Yoana Doncheva, Claire Kennedy Dietrich, Phillip Thienger, Dilara Akhoundova, Andrej Benjak, Achim Aigner, Rahul Kanadia, Sven Rottenberg, Mark A. Rubin

## Abstract

The minor spliceosome (MiS) is a specialized RNA-processing machinery upregulated in cancer, promoting oncogene expression. We uncovered an adaptive resistance mechanism driven by secretion of extracellular vesicles enriched in U6atac snRNA, which amplifies MiS activity and promotes therapy resistance. Here, we show that U6atac snRNA, a crucial MiS component, reverses this process when depleted, revealing it as a druggable vulnerability in therapy-resistant prostate and breast cancers. U6atac knockdown triggers R-loop–mediated DNA damage while impairing repair by downregulating key DNA repair factors, disabling both homologous recombination and non-homologous end joining. This dual effect sensitizes prostate and breast tumors to PARP inhibitors, cisplatin, and radiation, independent of BRCA status. Across multiple *in vitro* and *in vivo* models, MiS targeting demonstrates tumor-selective activity with minimal toxicity. These findings position U6atac as a central regulator of genome stability and establish MiS targeting as a promising approach to potentiate genotoxic therapy and overcome resistance.

**Statement of significance:** U6atac, a minor spliceosome component, is a crucial regulator of genome stability in cancer. Its knockdown triggers R-loop–driven DNA damage, downregulates DNA repair genes, and sensitizes tumors to DNA-damaging therapies while simultaneously blocking resistance mechanisms. Thus, minor spliceosome knockdown is a tumour-selective and broadly applicable therapeutic strategy.

## Main

Prostate cancer (PCa) and breast cancer (BCa) are the most common malignancies in men and women, respectively, and together account for more than one in eight cancer diagnoses worldwide as of 2024. Although outcomes are favorable in localized disease, survival drops dramatically once tumors metastasize, highlighting the pressing need for new therapeutic strategies.

Despite advances in precision and immune-based therapies, DNA-damaging agents, including PARP inhibitors, platinum chemotherapy, and radiation, remain central to treatment. Yet their impact is constrained. PARP inhibitors and platinum drugs show benefit mainly in the small fraction of patients (<5%) who harbor *BRCA1*/2 mutations, while adaptive resistance rapidly erodes efficacy in the rest. High-dose regimens also impose dose-limiting toxicity, restricting broader use. These limitations underscore the need for approaches that can sensitize *BRCA*-proficient tumors, expand the reach of DNA-targeting therapies, and overcome resistance while minimizing collateral damage, reinforcing the need for new therapeutic strategies.

Our previous work established the minor spliceosome (MiS) as a major vulnerability in PCa, where its disruption alters fundamental processes such as cell cycle progression and DNA repair. There are extensive parallels between PCa and BCa in their reliance on DNA repair pathways and their sensitivity to DNA-damaging therapies. Here, we further investigate the effects of blocking minor intron splicing on PCa and BCa, as it relates to DNA repair.

The MiS processes a rare class of introns in ~770 human genes. Many of these minor intron-containing genes (MIGs) encode regulators of cell cycle, MAPK signaling, and DNA repair, which are often hijacked during metastatic progression of cancer. In agreement, MiS activity is upregulated across metastatic PCa progression, and as expected, so are the MiS components, particularly the catalytic subunit U6atac snRNA^1^. We showed that MiS activity distinguishes aggressive tumors from benign tissue and that it aids in therapy resistance^1^. We identified the MiS as a dynamic regulatory node hijacked in cancer; these findings suggest that targeting its activity could selectively impair oncogenic pathways and restore sensitivity to DNA-damaging therapies.

Here, we discovered a direct mechanistic link between MiS function and DNA repair. We found that the MiS is a critical regulator of DNA repair, controlling key DNA damage response and survival factors, including DNA-PK, PARP1, BRCA1, TP53BP1, and CHK1/2. In this study, we investigated the impact of MiS knockdown (KD) in prostate and breast cancers. We found KD of U6atac impairs DNA repair and significantly sensitizes tumors to irradiation, cisplatin, and PARP inhibitors, even in BRCA-proficient cancers, demonstrating a shared therapeutic opportunity across tumor types. In response to DNA damage, U6atac levels and MiS activity increase, conferring therapy resistance. This process is supported by the secretion of U6atac-enriched extracellular vesicles, which propagate resistance upon uptake by recipient tumor cells. We uncovered a novel BRCA1-mediated regulation of MiS that involves modulation of U6atac levels, thereby positioning MiS as an ideal sensitizer to DNA-damaging agents. Indeed, siU6atac mediated MiS inhibition reduces the dose of irradiation, Cisplatin, and Olaparib required to kill cancer cells. This finding expands upon our previous report, where we nominated the MiS as a therapeutic target for lethal PCa, to now include first-line therapy for PCa and BCa. Our report, combined with that of Doggett et al., positions the MiS as a potential pan-cancer therapeutic target. A common concern with splicing inhibition is the associated toxicity that was observed with major spliceosome inhibitors. Thus, it is not surprising that the same concern extends to the MiS. However, it must be noted that the MiS targets <0.5% of introns, whereas the major spliceosome targets 99.5%. In fact, the therapeutic index of targeting the MiS was evident in our *in vivo* findings using xenograft models, where MiS KD enhanced therapeutic efficacy without observable systemic toxicity. Collectively, these results establish MiS KD as a translationally actionable strategy to improve efficacy, overcome drug resistance, and expand the application of DNA-targeting therapies to a broader patient population.

## Results

### U6atac upregulation correlates with an increase in the DNA repair program

Building on our prior discovery that U6atac is upregulated in therapy-resistant PCa^1^, we aimed to explore the association of U6atac and metastatically relevant biological programs such as DNA repair. Towards this goal, we performed spatial transcriptomics on advanced metastatic PCa. First, we identified areas with high and low U6atac expression through *in situ* hybridization with a customized probe against U6atac snRNA. U6atac high- or low-regions were then further stratified into tumor and tumor microenvironment (TME) regions based on pathology review and pan-cytokeratin expression (**Figure 1A and Supplemental Figure 1**). Then, the U6atac high- and low-coverage regions were subjected to spatial transcriptomics. We found that U6atac expression effectively segregates the tumor and TME into distinct clusters, as determined by principal component analysis (**Supplemental Figure 2**), and both U6atac-high tumors and TME showed a significant increase in MIG expression (**Supplemental Figure 3A, 3 B, and Supplemental Tables 1 and 2**). Gene expression analysis revealed that a subset of genes upregulated in U6atac high tumors enriched for immune response and oxidative phosphorylation, whereas none were observed in the U6atac low tumors (**Supplemental Figure 4**). Crucially, DNA repair genes, including PRKDC, ATR, XRCC5, XRCC6, NBN, or UBB, were upregulated in U6atac high tumors, thereby corroborating our previous findings that U6atac KD induces DNA damage (**Figure 1B and Supplemental Table 3**). In contrast, U6atac high TME showed only upregulation of two DNA repair genes, PTEN and POLE, suggesting that the MiS and MIGs are dynamically used in cancer progression (**Supplemental Figure 3C**).

**Figure 1.**
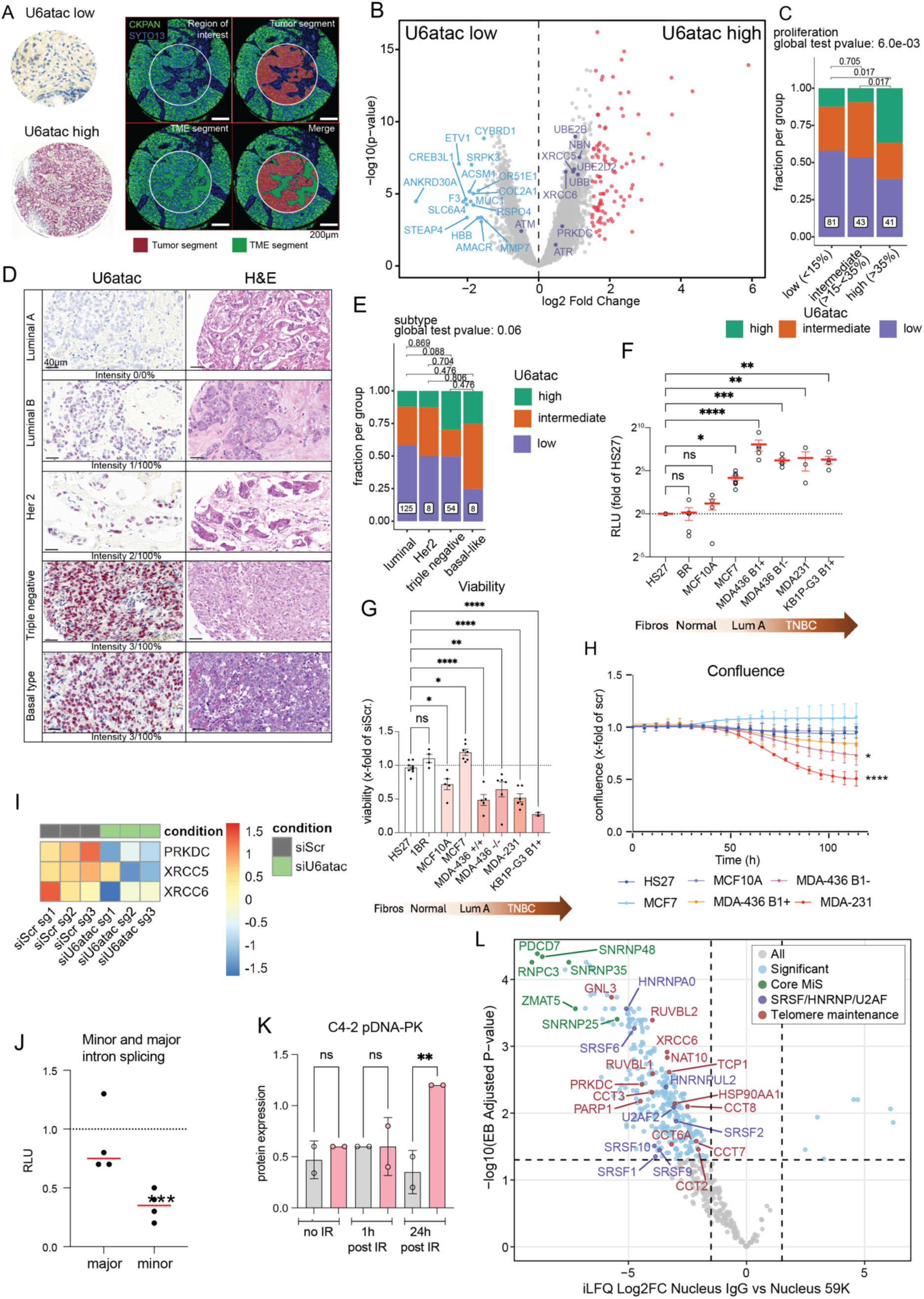
U6atac expression and MiS activity associate with tumor aggressiveness and DNA-PK signaling. (A) Spatial transcriptomics analysis of U6atac expression in high- and low-grade advanced prostate cancer (PCa). Left, region of interest manually selected for U6atac expression. Right, high-power view of tissue microarray core. The region of interest was segmented into CKPAN^+^ tumor and CKPAN^−^ tumor microenvironment segments (visualization antibodies: CKPAN and Syto13). (B) Genes highly expressed in U6atac-high tumors are enriched in both high- and low-grade PCa. Volcano plot showing differential gene expression between U6atac-high and U6atac-low tumors. Significantly upregulated genes are shown in red, significantly downregulated genes in blue, non-significant genes in grey, and DNA repair genes of interest in purple. (C) U6atac expression correlates with proliferation in breast cancer (BCa). Fraction of BCa cases showing low (purple), intermediate (orange), or high (green) U6atac expression in low-, intermediate-, and high-proliferation subgroups. (D) U6atac expression is increased in more aggressive BCa molecular subtypes. RNA in situ hybridization (RNAish) for U6atac (40× magnification, scale bar 40 μm) with corresponding H&E on the right. (E) Quantification of U6atac expression across BCa molecular subtypes. BCa cases showing low (purple), intermediate (orange), or high (green) U6atac expression are plotted by subtype (luminal A/B, HER2-positive, triple-negative, basal-like), with representative in situ images. (F) MiS activity correlates with BCa aggressiveness. Normalized luminescence values of minor spliceosome NanoLuc reporter in fibroblasts (Hs27, n = 8; BR, n = 5) and normal breast/BCa cell lines (MCF10A, n = 4; MCF7, n = 8; MDA-436 B1^+^ [BRCA-proficient], n = 5; MDA-436 B1^−^ [BRCA-deficient], n = 5; MDA-231, n = 3; KB1P-G3 B1^+^, n = 4). Each point is an independent experiment performed in triplicate; relative luminescence units (RLU; x-fold of Hs27) are plotted on a log_2_ y-axis. Mean ± SEM; Dunn’s post hoc tests versus Hs27: MCF7 (P = 0.0267), MDA-436 B1^+^ (P < 0.0001), MDA-436 B1^−^ (P = 0.0007), MDA-231 (P = 0.0096), KB1P-G3 B1^+^ (P = 0.0014); BR (P = 0.9544) and MCF10A (P = 0.6399) were not significantly different. Abbreviations: Fibros, fibroblasts; Lum A, luminal A; TNBC, triple-negative breast cancer. (G) Effect of MiS inhibition on cell viability correlates with BCa aggressiveness. Cell viability in fibroblasts (Hs27, n = 8; BR, n = 4), normal breast cells (MCF10A, n = 5), and BCa cell lines (MCF7, n = 7; MDA-436 B1^+^, n = 5; MDA-436 B1^−^, n = 6; MDA-231, n = 6; KB1P-G3 B1^+^, n = 2). Mean ± SEM; ordinary one-way ANOVA with Holm–Šídák post hoc test vs Hs27: MCF10A (P = 0.0351), MCF7 (P = 0.0351), MDA-436 B1^+^ (P < 0.0001), MDA-436 B1^−^ (P = 0.0036), MDA-231 (P < 0.0001), KB1P-G3 B1^+^ (P < 0.0001); BR (P = 0.1895) was not significant. Experiments were performed in triplicate. (H) Effect of MiS inhibition on tumor cell growth correlates with BCa aggressiveness. Growth curves of Hs27 (n = 4), MCF10A (n = 5), MCF7 (n = 6), MDA-436 B1^+^ (BRCA-proficient, n = 4), MDA-436 B1^−^ (BRCA-deficient, n = 3), and MDA-231 (n = 3) cells treated with siScrambled or siU6atac (2 nM). Cell growth is shown as a fold change relative to Day 0, normalized to siScrambled (dashed line). Mean ± SEM; ordinary two-way ANOVA (ns P > 0.05, *P < 0.05, **P < 0.01, ***P < 0.001). (I) DNA-PK and U6atac relationship. Heatmap of sgRNA enrichment scores from a C4-2 CRISPR–Cas9 screen. sgRNAs targeting PRKDC, XRCC5, and XRCC6 are selectively depleted in siU6atac-treated cells. Values depict normalized beta scores from MAGeCK MLE relative to Day 0. (J) Inhibition of DNA-PK preferentially decreases minor intron splicing. Normalized luminescence of dual minor/major spliceosome luciferase reporter in C4-2 cells (N = 5) treated with 10 μM DNA-PK inhibitor. Each point is an independent triplicate experiment, normalized to mCherry and DMSO control (set to 1, dashed line). One-way ANOVA with Šídák’s test: major intron splicing unchanged (P = 0.7452), minor intron splicing significantly reduced (P = 0.0004; ***). (K) MiS inhibition prolongs DNA-PK activation after irradiation. Quantification of pDNA-PK western blots in C4-2 cells (N = 4), transfected with siU6atac or siScr and exposed to 10 Gy irradiation. Cells were collected at baseline, 1 h, and 24 h post-IR. pDNA-PK was significantly increased at 24 h in siU6atac vs siScr (P = 0.0211). (L) DNA repair genes co-immunoprecipitate with the MiS component PDCD7. Volcano plots showing proteins enriched in C4-2 nuclear extracts immunoprecipitated with PDCD7 antibody (left) vs IgG control (right), based on three co-IP replicates. x-axis, log_2_ fold-change; y-axis, −log_10_ adjusted P value. Grey, non-differential; black/red, differentially expressed MIG-encoded proteins; triangles, proteins consistently differential across imputation; red, concordant iTop3 and iLFQ. MiS components are purple, DNA repair factors green, R-loop resolution proteins blue.

To expand our understanding of U6atac and DNA repair, we studied BCa with *BRCA1/2* mutations. Its similarities to PCa in the DNA damage response mechanisms highlight the potential for cross-therapy strategies and, thereby, new opportunities for targeted treatments^2^. We therefore investigated U6atac expression in tissue microarrays of BCa. Tumors with high U6atac showed a higher proliferation index, as determined by Ki67 immunohistochemistry, which was available for a subset of patients (N=165). The proliferation fraction was classified as low (<15%), intermediate (15%-<35%), or high (>35%). Tumors of patients presenting a high proliferation fraction showed a higher U6atac-expression when compared with low and intermediate proliferating tumors (p=0.02 and p=0.02, respectively) (**Figure 1C and Supplemental Figure 5**). In addition to the proliferative index, BCa is further stratified by IHC for specific markers, such as ER/PR and/or Her2. We discovered that U6atac expression levels increased progressively from less metastatic (luminal A and B) to more metastatic (triple-negative and basal BCa). A higher proportion of cases with elevated U6atac expression was observed in the triple-negative (30%) and basal-like (25%) subtypes compared to the luminal A/B (12%) and HER2-positive (13%) subtypes (**Figure 1D**). However, these differences did not reach statistical significance across groups (p > 0.05 for all comparisons) (**Figure 1E**). Given that U6atac is the rate-limiting factor for MiS activity, we investigated splicing activity by employing splice reporters where the coding sequence of nanoluciferase (Nluc) is interrupted by a minor intron, and that of firefly luciferase is interrupted by a major intron. Thus, the chemiluminescence of Nluc versus firefly provides a readout of basal splicing activity across various BCa cell lines, including non-cancerous and cancerous lines with different grades of invasiveness. We found that increased levels of U6atac were generally associated with enhanced MiS activity (**Figure 1F**). Specifically, normal fibroblasts (HS27 and BR) and breast epithelial cells (MCF10A) exhibited low MiS activity, whereas BCa cell lines starting with MCF7 showed elevated MiS activity, with the highest in those representing triple-negative BCa (**Figure 1F**). These data suggest that, as in PCa^1^, U6atac and minor intron splicing correlate with BCa aggressiveness, supporting the hypothesis that BCa is also susceptible to U6atac KD. Indeed, KD of U6atac, which decreases minor intron splicing but not major intron splicing (**Supplemental Figure 6**), showed no significant reduction in cell viability or growth in normal fibroblasts and primary breast cells, yet demonstrated a marked decrease in viability and growth in advanced BCa cells (**Figure 1G and H**). This effect was most pronounced in triple-negative BCa cell lines (MDA-231 and MDA-436).

To understand how MiS inhibition in aggressive cancer cells works, we performed an unbiased genome-wide CRISPR screen in PCa C4-2 cells (manuscript in preparation). As shown in **Figure 1I**, PRKDC (DNA-PK), XRCC5, and XRCC6, which recruit DNA-PK to DNA damage sites, were identified as synthetic lethal with siU6atac, as the targeting sgRNAs for these genes were significantly depleted in cells treated with siU6atac. This finding agreed with our spatial transcriptomics data, in which U6atac-high tumors correlated with genes involved in DNA repair, such as DNA-PK (*PRKDC*), a key kinase in the DNA damage response (**Figure 1B**). Therefore, we used a DNA-PK inhibitor in our C4-2 cells expressing the dual splice reporter, revealing that blocking DNA-PK reduced minor intron splicing, whereas major intron splicing remained unaffected (**Figure 1J**). Moreover, knockdown (KD) of U6atac reduced DNA-PK protein levels following irradiation in PCa C4-2 and BCa MDA-436 cells, supporting the relationship of DNA-PK to MiS **(Supplemental Figure 7**). However, phosphorylation of DNAPK was increased after irradiation plus MiS inhibition (**Figure 1K and Supplemental Figure 7**). This suggests that although U6atac KD reduces DNA-PK expression, DNA-PK phosphorylation persists, likely due to the extensive DNA damage in irradiated cells **(Supplemental Figure 7**). The surprising interaction of MiS and DNA repair genes was further confirmed when we performed immunoprecipitation of a crucial MiS protein, PDCD7, which is part of the MiS complex that recognizes the 5’ splice site of minor introns. Consistent with the role of PDCD7, we identified RNPC3, SNRP25, SNRNP35, SNRNP48, and ZMAT5, which are MiS-specific proteins^3^, were co-immunoprecipitated (**Figure 1L, Supplemental Figure 8, and Supplemental Tables 4 and 5**). Other crucial splicing factors such as U2AF2, SRSF2, and hnRNPs^4^ were also identified in our PDCD7 pulldown mass spec data. As expected, factors such as ZCRB1, ARMC7, ZRSR2, and others known to play a catalytic role in minor intron splicing are absent in the pulldown^5^. Thus, confirming the role of PDCD7 in the 5’ splice site recognition by the MiS. In this context, co-immunoprecipitation of DNA repair genes such as PRKDC, XRCC5, XRCC6, PARP1, CDC5L, and RUVBL1 was noteworthy, further reinforcing the upregulation of DNA repair genes in U6atac high tumors and, by extension, the MiS activity (**Figure 1B and H**).

### U6atac Knockdown Impairs Double-Strand Break Repair and Induces a BRCAness Phenotype

There are two main types of DNA repair pathways: one is non-homologous end joining (NHEJ), which is mediated by DNA-PK, and the other is homologous recombination (HR), which is regulated by BRCA1 and BRCA2^6^. The interaction of MiS-specific proteins with DNA repair genes upregulated in U6atac high tumors led us to investigate which DNA repair pathway is regulated by MiS^1^. For this purpose, we leveraged the DSB spectrum reporter assay in HEK239T cells, as described by van de Kooij et al., in which the relative fluorescence of BFP and GFP together reports the activation and/or inhibition of NHEJ and HR repair activity (**Figure 2A**)^7^. Using this assay, we demonstrate that *PRKDC (DNA-PK)* inhibition reduces NHEJ, as indicated by reduced BFP (**Figure 2A** and **Supplemental Figure 9**). The specificity of NHEJ inhibition was reflected in the lack of BFP reduction when we decreased BRCA1 and 2. Specifically, U6atac KD showed a reduction in BFP signal similar to that of PRKDC KD, suggesting that blocking MiS inhibits NHEJ. In contrast, we observed a decrease in GFP signal only in BRCA1 and BRCA2 KD conditions, but not in PRKDC KD, suggesting inhibition of HR. Notably, a similar reduction of GFP signal was observed upon siU6atac-mediated MiS inhibition. Taken together, U6atac-mediated MiS inhibition reveals the crucial role of minor intron splicing in both NHEJ and HR in cancer cells. Thus, underscoring the critical role of U6atac in promoting DNA repair, thereby bolstering our previous observation that U6atac-high cancers upregulate DNA repair genes (**Figure 2A and Supplemental Figure 9**).

**Figure 2.**
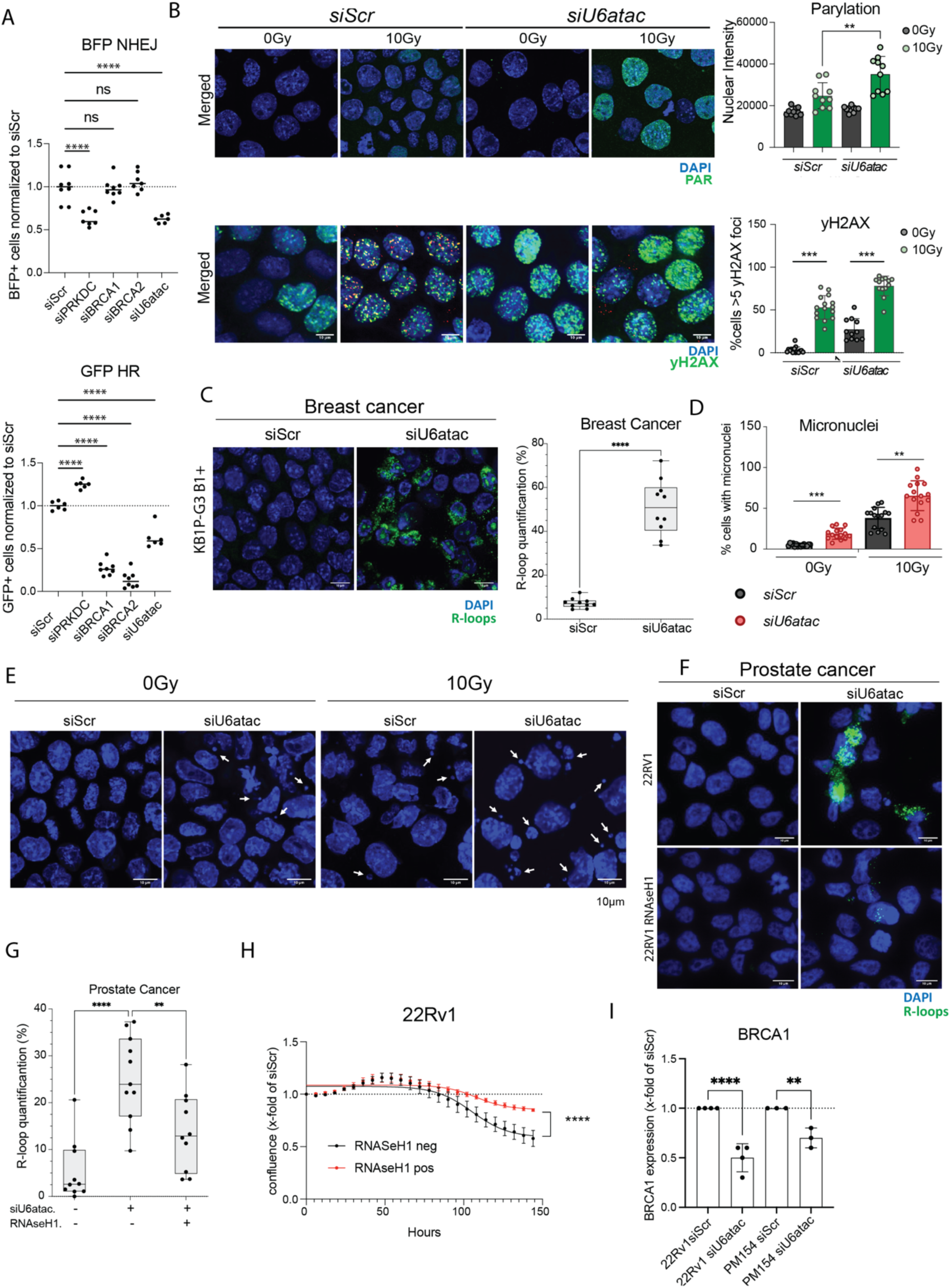
MiS inhibition induces DNA damage, R-loops, micronuclei, and involves BRCA1. (A) MiS inhibition affects non-homologous end joining (NHEJ) and homologous recombination (HR). Quantification of double-strand break spectrum assay in HEK293T cells treated with indicated siRNAs. Each dot represents an experiment run in duplicate; bars, mean ± SD. ANOVA with Tukey’s multiple comparisons. (B) MiS inhibition increases PARylation and γH2AX. Upper, representative PAR immunofluorescence images in KB1P-G3 B1^+^ cells 24 h post-IR, treated with indicated siRNAs. Mann–Whitney test, **P < 0.01. Lower, representative γH2AX foci images in the same conditions; Mann–Whitney test, ***P < 0.001. (C) MiS inhibition increases R-loop formation in BCa. Representative R-loop immunofluorescence images in KB1P-G3 B1^+^ cells 24 h post-IR with indicated siRNAs. Each dot is an acquisition field (~150 nuclei/field). Welch’s t-test: t = 10.86, df = 9.6, P < 0.0001 for siU6atac vs siScr. (D) MiS inhibition increases micronuclei formation in BCa. Quantification of micronuclei 24 h post-IR in KB1P-G3 B1^+^ cells treated with the indicated siRNAs. Scatterplots show the percentage of cells with micronuclei. Mann–Whitney test, **P < 0.01, ***P < 0.001. (E) Representative images of micronuclei in KB1P-G3 B1^+^ cells 24 h post-IR with indicated siRNAs. (F) MiS inhibition increases R-loop formation in PCa. Representative R-loop immunofluorescence in 22Rv1 wild-type or RNaseH1-expressing PCa cells treated with siScr or siU6atac, 24 h post-IR. (G) Quantification of R-loops in 22Rv1 wild-type or RNaseH1-expressing cells treated as in (F). Each dot is an acquisition field (~150 nuclei/field). One-way ANOVA, P < 0.0001; Šídák post hoc: WT siU6atac vs WT siScr, ****P < 0.0001; RNaseH1 expression significantly reduces R-loops (**P = 0.0065). (H) RNase H1 expression rescues growth defects caused by MiS inhibition. Growth curves of 22Rv1 cells overexpressing RNaseH1 (RNaseH pos) vs controls (RNaseH neg) after siU6atac (2 nM, n = 3). Confluence is shown as fold change vs Day 0, normalized to siScrambled (dotted line). Mean ± SD; two-way repeated measures ANOVA: significant time and time × RNaseH1 interaction (P < 0.0001). Šídák’s test shows rescue from ~100 h onward (*P < 0.05 to **P < 0.0001). (I) BRCA1 but not BRCA2 knockdown decreases U6atac expression in PCa. BRCA1 mRNA expression in 22Rv1 (n = 4) and PM154 (n = 3) cells treated with siU6atac (2 nM), shown as fold change vs siScrambled and normalized to GAPDH/ACTB. Bars, mean ± SD; dots, individual experiments. One-way ANOVA (P < 0.0001) with Šídák’s test: BRCA1 significantly decreased in both lines (22Rv1, ****P < 0.0001; PM154, P = 0.0042).

Another mechanism for DSB formation is single-strand breaks (SSBs), which are primarily repaired by Base Excision Repair (BER)^8^. PARP1, a known MIG, first detects most SSBs, which initiates repair through PARylation of its targets^8^. Indeed, an increase in PARylation was confirmed by IF in both BCa KB1P-G3 B1+ (**Figure 2B**) and PCa C4-2 and 22Rv1 cell lines (**Supplemental Figure 10A**). Upon MiS inhibition via siU6atac, we observed a decrease in canonical PARP1 expression in PCa 22Rv1 cells, accompanied by an increase in cleaved PARP1 and PARylation, which is a strong indicator of SSBs^9^. Thus, revealing a potential mechanism through which MiS inhibition induces PARPness phenotype in these cells (**Supplemental Figure 10A, B, and C**). Given the relationship between MiS inhibition, PARylation, and SSB, it was not surprising that this effect was markedly enhanced when cells were co-treated with a single dose of 10Gy radiation, which is known to cause DNA damage (**Figure 2B and Supplemental Figure 10A**). The conversion of SSBs to DSBs is a pivotal event in the cellular response to DNA damage, with profound implications for cancer therapy^10^. This process can drive genomic instability, influence tumor progression, and determine the effectiveness of treatments such as PARP inhibitors and DNA-damaging agents such as radiation and platinum-based chemotherapy. Therefore, we explored whether MiS inhibition leads to elevated DSB levels. Indeed, BCa and PCa cells treated with siU6atac showed increased levels of γH2AX, a marker for DSBs (**Figure 2B, lower panel and Supplemental Figure 11**). Again, the effect was more pronounced when cells were co-treated with 10Gy of radiation, suggesting that siU6atac enhances the effect of DNA-damaging agents. This finding indicates that DNA-damaging agents are aided by DNA damage caused by MiS inhibition and splicing defects. Indeed, the accumulation of unspliced RNA upon siU6atac treatment promotes the formation of RNA–DNA hybrids, known as R-loops, which contribute to genomic instability by impairing DNA replication and repair. Therefore, we performed immunofluorescence (IF) analysis using the S9.6 antibody, which specifically recognizes the A-form helical conformation characteristic of DNA-RNA duplex^11^. Our results demonstrate that indeed, U6atac KD induces a significant increase in R-loop accumulation in both BCa and PCa cells (**Figure 2C, 2F, 2G and supplemental Figure 12**). To confirm that siU6atac-mediated MiS inhibition led to elevated R-loop formation, we overexpressed RNase H1 in PCa cells, which significantly reduced R-loop accumulation (**Figure 2F and G, and Supplemental Figures 13 and 14**). Of note, the same experiment did not work in BCa cells, as RNAse H1 overexpression was highly suppressed in these cells. Regardless, the PCa experiment provides proof of concept that MiS inhibition results in elevated R-loop formation, which contributes to DNA damage, as reflected by both micronuclei formation and elevated yH2AX levels. Consistently, RNAse H1 overexpression, in addition to reducing R-loops, also resulted in a marked decrease in γH2AX foci (**Supplemental Figure 13**). R-loops are known to contribute to genomic instability by impairing DNA replication and repair. The persistence of R-loops or failure to resolve them can lead to substantial DNA damage, replication stress, and genomic instability^12^. Consistent with these findings, we observed that siU6atac treatment induced micronuclei, a global marker of chromosomal instability (**Figure 2D and E**). Notably, this effect was further enhanced in irradiated BCa cells (**Figure 2D and E**). Since reduced yH2AX indicates reduced DNA damage, it is not surprising that we also observed rescue of the siU6atac-induced growth delay, as evidenced by significantly longer maintenance of normal cell growth (**Figure 2H and Supplemental Figure 14**).

DNA-damaging agents induce both SSBs and DSBs. Unrepaired SSBs, particularly when PARP1-mediated repair signaling is impaired, can collapse replication forks and thereby convert SSBs into DSBs. In both PCa and BCa BRCA1 and BRCA2 play a vital role in HR-mediated repair of DSBs, and with BRCA1 being re-classified as a minor intron-like gene^13^we thought to study the effect of siU6atac on BRCA1/2 expression. We found that siU6atac mediated MiS inhibition did not change BRCA2 expression (**Supplemental Figure 15**). Yet it did significantly decrease BRCA1 expression in 22Rv1 PCa cells and PM154 organoids (**Figure 2I**). The ATM/ATR signaling axis senses DSBs and coordinates homologous recombination and checkpoint activation. Another mechanism for DSB sensing is the DNA-PK-dependent NHEJ pathway^14^. To understand how MiS inhibition impairs both HR and NHEJ pathways, we investigated how MiS inhibition engages with DNA damage checkpoint activation. Specifically, we assessed both total and phosphorylated levels of checkpoint kinases CHK1 and CHK2 at 1 hour and 24 hours following ionizing radiation in both BCa and PCa cells with and without siU6atac treatment. Notably, at 24 hours post-irradiation, all cancer cell lines pre-treated with siU6atac showed a marked reduction in levels of phosphorylated CHK1 and CHK2 protein (**Supplemental Figures 16 and 17**), an effect absent in scrambled control cells. To understand the decrease in CHK1 and CHK2 activation after irradiation, we investigated the phosphorylation status of the critical checkpoint kinase upstream kinase ATM after siU6atac treatment. We observed a reduction in ATM phosphorylation after irradiation in siU6atac-treated cancer cells (**Supplemental Figure 18**). Moreover, we observed a decrease in ATR protein levels, another CHK1 and CHK2 upstream kinase in cells treated with siU6atac and 10Gy radiation in BCa but not in PCa (**Supplemental Figure 19**). These findings connect MiS and DNA repair. Thus, siU6atac mediated minor intron splicing defect induces R-loop accumulation, which we believe is the source of both SSBs and DSBs. Simultaneously, MiS inhibition disrupts crucial MIGs and non-MIGs required for DNA damage response (**Supplemental Figure 20**).

### U6atac Knockdown Sensitizes Tumors to DNA-Damaging Therapies

The role of MiS in the DNA damage repair pathway led us to hypothesize that its inhibition would sensitize PCa and BCa cells to standard-of-care DNA-targeting agents, including radiation, cisplatin, and Olaparib. We first investigated previously conducted PARP inhibitor (PARPi, Olaparib) CRISPR sensitivity screens in BCa^15,16^ and PCa that were designed to identify genes synergistic with Olaparib treatment. Here, we found that knockout (KO) of unique MiS protein components appears to enhance sensitivity to Olaparib (**Supplemental Figure 21**). This data suggests that MiS inhibition is indeed synergistic with Olaparib, so we tested the effects of siU6atac combined with Olaparib by treating PCa/BCa cells with varying siU6atac concentrations (0, 2, 4, 8, 16, 32nM) and Olaparib (0, 0.5, 1, and 2 μM). We observed a trend indicating synergy, with the highest synergy scores of 10 in 22Rv1 and 5 in DU145 PCa cells and 26 in BCa (KB1P-G3 B1+) cells (**Figure 3A and Supplemental Figure 22A).** To also assess potential combinatorial effects with platinum-based cancer therapies, we next examined synergy between siU6atac (0, 2, 4, 8, 16, 32 nM) and cisplatin (0,1, 2, 3 μM). Co-treatment consistently yielded elevated synergy scores in BCa (18.4) and in PCa DU145 cells (33) but not in PCa 22Rv1 cells (1.8) (**Figure 3B and Supplemental Figure 22B**). These results indicate a robust therapeutic interaction, whose magnitude appears to be influenced by cell- and tissue-specific factors. These results were first corroborated in a functional clonogenic assay, which revealed that siU6atac significantly reduced the doses of Olaparib (5x), radiation (3x), and cisplatin (2x) in both BRCA-deficient and proficient BCa cells (**Figure 3C and Supplemental Figure 23A**). The same was observed in PCa (DU145, MSK16, and C4-2) cells with siU6atac plus radiation, where siU6atac reduced the dose of irradiation by (3x), whereas C4-2 cells were highly sensitive to both radiation and siU6atac by themselves, thus synergy was hard to assess by clonogenic assay (**Supplemental Figure 23B**). However, when we used CellTiter-Glo®, we observed that siU6atac reduced cell viability on its own, but significantly enhanced the effects of cisplatin, Olaparib, or radiation in C4-2 and 22Rv1 PCa cells (**Figure 3D, E and F and Supplemental Figure 24A and B**). Notably, this effect was observed even in BRCA-proficient models, typically resistant to Olaparib, supporting our hypothesis that siU6atac broadens the therapeutic window by sensitizing tumors to Olaparib and other DNA-damaging agents, regardless of BRCA status. Of note, siU6atac co-treatment with DNA-damaging agents showed a significantly better outcome in KB1P-G3 B1+ and B-BCa cells and LNCaP, LNCaP Rb1/TP53 KO, MSK16, and DU145 PCa cells (**Supplemental Figure 24A, B and C**). However, in C4-2 and 22Rv1 cells, siU6atac alone was so drastic that co-treatment with DNA-damaging agents improved outcome only marginally (**Figure 3D, E, F and Supplemental Figure 24 A and B**). This highlights cell specificity and that siU6atac sensitizes to DNA-damaging treatments.

**Figure 3.**
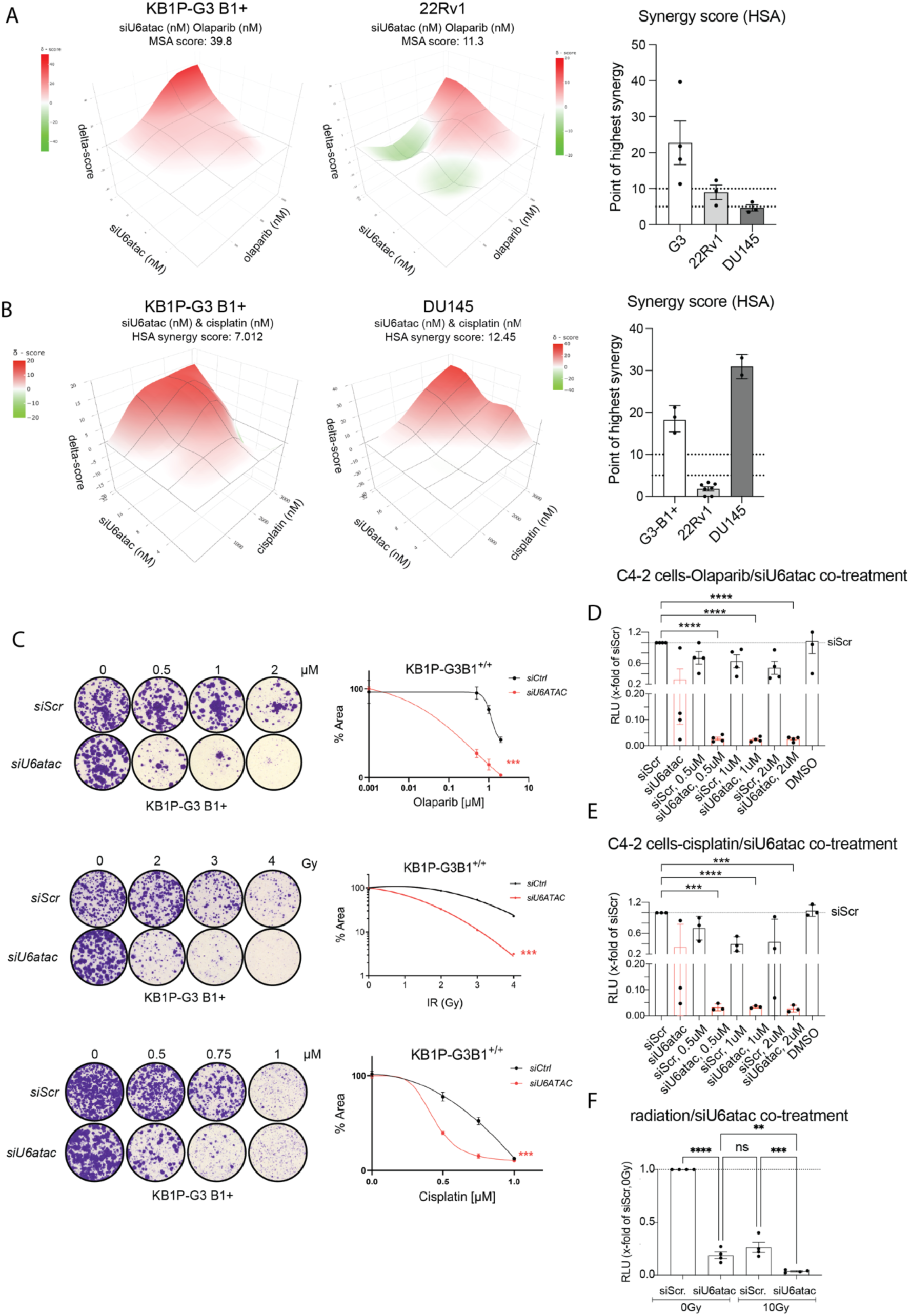
MiS inhibition sensitizes prostate and breast cancer cells to DNA-damaging therapies. (A) MiS inhibition sensitizes cells to olaparib in synergy assays. 3D interaction landscapes of siU6atac (0–32 nM) and olaparib (0–2 μM) in KB1P-G3 B1^+^ and 22Rv1 cells, analyzed by HSA model in SynergyFinder. Red, synergy (>0); green, antagonism (<0). Bar graph shows the highest synergy scores from three independent biological experiments (triplicates). (B) MiS inhibition sensitizes cells to cisplatin in synergy assays. 3D interaction landscapes for siU6atac (0–32 nM) and cisplatin (0–3 μM) in KB1P-G3 B1^+^ and 22Rv1, analyzed as in (A). (C) MiS inhibition enhances response to DNA-damaging agents in clonogenic assays. Left, representative growth assays in KB1P-BRCA1^+^ cells treated with indicated siRNAs and exposed to olaparib, IR, or cisplatin. Right, two-way ANOVA with Dunnett’s test; ***P < 0.001. (D) MiS inhibition sensitizes C4-2 cells to olaparib in viability assays. Cell viability after siU6atac (2 nM) and olaparib (0.5–2 μM), shown as x-fold of siScrambled (dotted line). N = 4 biological replicates (triplicates). Repeated measures ANOVA with Holm–Šídák: ****P < 0.0001. (E) MiS inhibition sensitizes C4-2 cells to cisplatin. Cell viability after siU6atac (2 nM) and cisplatin (0.5–2 μM), shown as x-fold of siScrambled. N = 3 biological replicates (triplicates). Repeated measures ANOVA (P = 0.0713, ns) with Holm–Šídák; siU6atac + cisplatin significantly reduced viability at all doses (***P = 0.0004, ****P < 0.0001). (F) MiS inhibition sensitizes C4-2 cells to irradiation. Cell viability after siU6atac (2 nM) ± IR (10 Gy), shown as x-fold of siScrambled. N = 4 biological replicates (triplicates). One-way ANOVA with Fisher’s LSD: siScr vs siU6atac (P < 0.0001), siU6atac vs siU6atac 10 Gy (P = 0.0028), siScr 10 Gy vs siU6atac 10 Gy (P = 0.0001), siU6atac vs siScr 10 Gy (P = 0.0958).

### U6atac expression increases with genotoxic stress in cancer

Recent studies have begun to link MiS components such as RNPC3 and SF3B1 to genome maintenance^17,18^. We have shown that prolonged androgen receptor signaling inhibitor (ARSi) treatment induces cellular stress and increases U6atac expression^1^. To assess whether genotoxic stress induced by DNA-damaging therapies exerts a similar effect, we investigated the impact of radiation, cisplatin, and Olaparib treatments on U6atac levels. Like ARSi enzalutamide treatment in PCa C4-2 cell line, all three treatments significantly upregulated U6atac expression compared to untreated controls (**Figure 4A**). Focusing specifically on radiation, we observed that this effect was selective to malignant prostate and breast cells: normal cell lines (HS27, MCF10A) showed no significant change in U6atac expression post-radiation, whereas prostate and breast cancer cell lines exhibited a marked increase (**Figure 4B**). Notably, this radiation-induced upregulation was more pronounced in BRCA wild-type cancer cells (LNCaP, C4-2, MDA-436 B1+, and MDA231) as compared to BRCA-deficient lines (DU145, 22Rv1, MCF7, and MDA-436 B1-) (**Figure 4B**). Underscoring the relationship of U6atac expression with genotoxic stress, we observed upregulation of U6atac in human patient-derived PCa organoids (PM154) exposed to 2-8 Gy radiation. Crucially, pre-treatment with siU6atac effectively suppressed this radiation-induced U6atac upregulation (**Figure 4C**).

**Figure 4.**
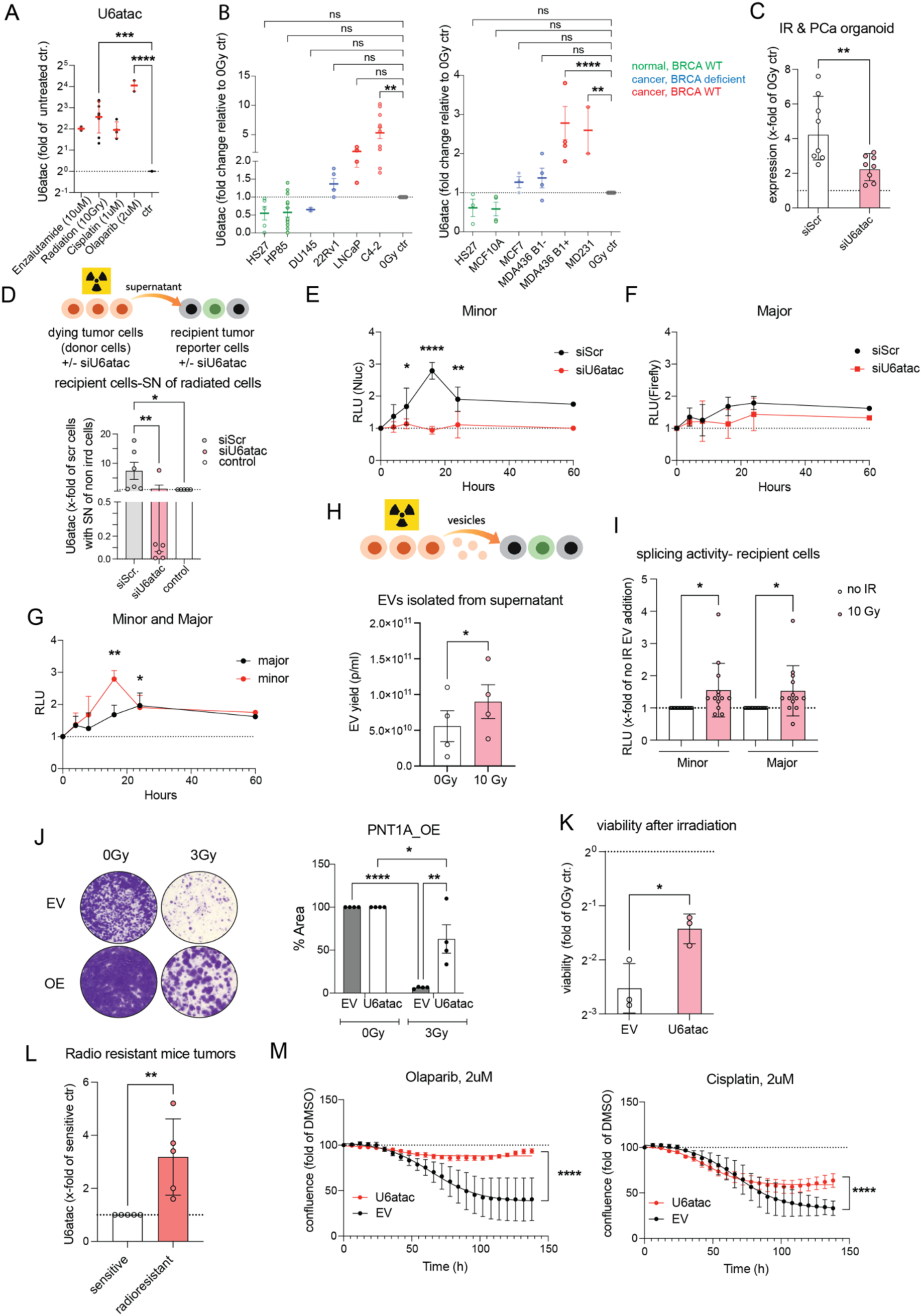
Genotoxic stress and extracellular vesicles modulate U6atac/MiS activity and therapy resistance. (A) Genotoxic stress increases U6atac expression. U6atac snRNA levels in C4-2 cells treated with enzalutamide (10 μM, n = 3), radiation (10 Gy, n = 8), cisplatin (2 μM, n = 4), or olaparib (2 μM, n = 2). Expression is shown as x-fold of untreated control, normalized to GAPDH/ACTB, on a log_2_ y-axis. One-way ANOVA (P < 0.0001) with Šídák’s test: radiation (*P = 0.0106) and olaparib (****P < 0.0001) significantly increased U6atac. (B) Radiation increases U6atac in BRCA1-proficient cancer cells. U6atac snRNA expression after 10 Gy IR in fibroblasts (Hs27) and prostate or breast cancer cell lines, shown as x-fold of non-irradiated controls (normalized to GAPDH/ACTB). Prostate: Hs27 (n = 5), HP85 (n = 13), DU145 (n = 2), 22Rv1 (n = 5), LNCaP (n = 4), C4-2 (n = 11); ANOVA (P < 0.0001) with Dunnett’s: C4-2 vs 22Rv1, adjusted P = 0.0011. Breast: Hs27 (n = 3), MCF10A (n = 4), MCF7 (n = 3), MDA-MB-436 B1^−^ (n = 4), MDA-436 B1^+^ (n = 5), MDA-231 (n = 2); ANOVA (P < 0.0001) with Dunnett’s: MDA-436 B1^+^ (P = 0.0005) and MDA-231 (P = 0.0089) vs MCF10A. Color code: green, normal BRCA wild-type; blue, BRCA-deficient; red, BRCA wild-type cancer. (C) Radiation increases U6atac in PCa organoids, blocked by siU6atac. U6atac expression in PM154 PCa organoids pre-treated with siU6atac or siScr (2 nM) for 48 h before IR (2–8 Gy), shown as x-fold of non-IR control. Mean ± SD, N = 8 paired samples. Paired t-test: t(7) = 4.382, P = 0.0032; mean difference –2.238 (95% CI –3.445 to –1.030), R² = 0.733. (D) Supernatant from irradiated cancer cells increases U6atac in recipient cells. Schematic of transfer of supernatant from irradiated C4-2 donor cells to recipient C4-2 reporter cells pre-treated with siScr or siU6atac. Bar plots show U6atac expression in recipients after 16–24 h, normalized to GAPDH/ACTB (N = 6). IR, irradiation. (E) Supernatant from irradiated cancer cells increases MiS activity in recipient cells. Normalized minor spliceosome (NanoLuc) activity in C4-2 reporter cells pre-treated with siScr or siU6atac (2 nM) and then given supernatant from irradiated C4-2 donors (10 Gy). Luminescence measured at indicated times (N = 3; mean ± SD). Mixed-effects model (REML) with Fisher’s LSD: siU6atac vs siScr differs at 16 h (P = 0.0382), 24 h (P < 0.0001), and 60 h (P = 0.0063), not at other times. (F) Major spliceosome activity is not significantly altered by supernatant transfer. Two-way repeated measures ANOVA for major spliceosome (Firefly): treatment (P = 0.6543), time (P = 0.1092), interaction (P = 0.646); no significant differences by Fisher’s LSD. (G) The supernatant from irradiated cells induces stronger MiS activity than MaS activity. Comparison of normalized minor (NanoLuc) vs major (Firefly) reporter activity. Mixed-effects (REML) analysis shows a significant effect of spliceosome type (P = 0.0245), with higher activity in MiS (difference –0.273, 95% CI –0.479 to –0.066). Fisher’s LSD: significant differences at 16 h (P = 0.0039) and 24 h (P = 0.0107), not at other times (P > 0.63). (H) Irradiation increases EV concentration in PCa. Schematic of EV transfer and EV yield (particles/ml) from supernatants of non-irradiated vs 10 Gy-irradiated C4-2 cells (N = 4 paired samples; mean ± SEM). Paired t-test, P = 0.049. (I) EVs from irradiated cells increase MiS and MaS activity in recipient cells. Minor and major spliceosome activity in C4-2 reporter cells after EV transfer from irradiated or non-irradiated donors, shown as x-fold of EVs from non-irradiated cells (N = 14; mean ± SD). One-way ANOVA (P = 0.0186); Holm–Šídák: EVs from irradiated donors significantly increase both minor (P = 0.0247) and major (P = 0.0495) reporter activity. (J) U6atac overexpression confers radioresistance in clonogenic assays. PNT1A cells overexpressing U6atac (OE) or empty vector (EV) were irradiated with 0 or 3 Gy. Representative colony images (left) and quantification of colony area (right) are shown (N = 4; mean ± SEM). Two-way ANOVA shows significant effects of irradiation and U6atac, with an interaction (P = 0.0050). Šídák’s tests: EV 0 Gy vs EV 3 Gy (****P < 0.0001), U6atac 0 Gy vs 3 Gy (*P = 0.0245), EV 3 Gy vs U6atac 3 Gy (**P = 0.0012). (K) U6atac overexpression confers radioresistance in viability assays. Viability of C4-2 cells overexpressing U6atac or EV 24 h after 10 Gy IR, shown as x-fold of non-IR control (log_2_ y-axis). N = 3; mean ± SD. Welch’s t-test, P = 0.0207. (L) Radioresistant mouse breast tumors display increased U6atac. U6atac expression in radioresistant vs matched radiosensitive mouse tumors, normalized to GAPDH and shown as x-fold of sensitive tumors (N = 5/group; mean ± SD). Mann–Whitney test, P = 0.0079. (M) U6atac overexpression confers resistance to olaparib and cisplatin. Confluence of C4-2 cells overexpressing U6atac (OE) or EV treated with olaparib (2 μM, left) or cisplatin (2 μM, right), shown as x-fold of DMSO control (N = 3; mean ± SD). For olaparib, Šídák’s test: U6atac OE vs EV significantly higher from 72 h (P = 0.0167) through 144 h (P < 0.0001). For cisplatin, significant from 84 h (P = 0.0269) through 144 h (P < 0.0001).

### U6atac-Rich Extracellular Vesicles Drive Adaptive Resistance to Genotoxic Therapy

Extracellular vesicles (EVs) are 30-150 nm microvesicles that bud directly from the cell membrane, mainly consisting of RNA and some proteins. Recently, it was shown that glioblastoma and ovarian cancer cells treated with radiation and cisplatin secrete EVs containing snRNAs, including U6atac^19,20^. Uptake of these EVs by recipient tumor cells was shown to enhance splicing and promote treatment resistance^19,20^. However, these studies did not interrogate minor intron splicing. Given that irradiation resulted in the most significant upregulation of U6atac snRNA, we reasoned that the increased resistance in recipient cells was owed to U6atac and, thus, minor intron splicing. To test this idea, we irradiated therapy-resistant PCa C4-2 cells (donor cells) and transferred their supernatant to recipient naïve C4-2 cells. The supernatant of irradiated donor cells triggered an increase in U6atac expression in recipient cells. However, when the naïve cells were pre-treated with siU6atac, we did not observe an increase in U6atac expression (**Figure 4D**). Similarly, the supernatant from cisplatin- or Olaparib-treated cancer cells induced an increase in U6atac expression in recipient cancer cells at both 16 and 24 hours. This increase was blocked when the recipient cells were pre-treated with siU6atac (**Supplemental Figure 25A**). U6atac levels in response to irradiation were dependent on both the donor and recipient cell type. For example, normal fibroblasts treated with supernatant from either irradiated fibroblasts or PCa C4-2 cells did not exhibit an increase in U6atac levels (**Supplemental Figure 26)**. Similarly, when supernatant derived from irradiated fibroblasts was used to treat PCa C4-2 cells, we did not observe a significant elevation in U6atac levels. The relationship between the irradiated cells that provide the supernatant, and the recipient cells is highly specific, as it works only when the source and recipient are PCa C4-2 cells (**Supplemental Figure 26**). Thus, U6atac upregulation might require both tissue specificity and a donor source from cancer cells. Next, we tested whether the upregulation of U6atac in recipient cells also led to elevated MiS activity. For this, we employed the dual splice reporter, which revealed that the recipient cells showed a peak in MiS activity 10-24 hours after exposure to supernatant from irradiated cells (**Figure 4E and G**). In contrast, major spliceosome activity remained unaffected (**Figure 4F and G**). The increase in MiS activity was attributable to elevated U6atac levels, as recipient cells were pre-treated (24h) with siU6atac and did not show an increase in MiS activity after exposure to supernatant derived from irradiated cells (**Figure 4E**). The same was observed with cisplatin and Olaparib (**Supplemental Figure 25B**).

The observation that U6atac snRNA, normally characterized by rapid turnover, appears both in the supernatant and in recipient cells led us to hypothesize that it is packaged within EVs. Indeed, we found that radiation significantly increased the number of EVs released into the supernatant of PCa cells (**Figure 4H**). Since our previous experiments used the entire supernatant, which led to upregulation of U6atac, we wanted to confirm that EVs were responsible for this upregulation. Treatment of PCa C4-2 cells with purified EVs from irradiated C4-2 cells alone resulted in elevated minor and major intron splicing as reported by the dual splice reporters (**Figure 4I**). Based on the siU6atac mediated reduction in MiS activity in recipient cells, we hypothesize that it is U6atac levels that primarily drive MiS activity and are integrated with the DNA damage response mechanism. If this is true, then we reasoned that increasing U6atac levels would protect against DNA damage. Indeed, overexpression of U6atac in normal prostate PNT1A cells led to increased resistance to irradiation as well as to cisplatin and Olaparib treatment (**Figure 4J, K, M, and Supplemental Figure 27**). Moreover, in radioresistant Brca1-deficient mouse mammary tumors^21^ we observed significant U6atac expression, further validating our findings in vivo (**Figure 4L**). In these models, *Barazas et al*. demonstrated that loss of 53BP1 creates an acquired vulnerability to radiotherapy in Brca1-deficient cells both *in vitro* and *in vivo*^21^. In line with this, we observed decreased 53BP1 protein expression following siU6atac treatment (**Supplemental Figures 28 and 29**).

Overall, these results suggest that DNA-damaging agents, such as radiation and cisplatin, and the indirect effect of Olaparib upregulate U6atac levels, which are then secreted in EVs and taken up by neighboring cancer cells.

### BRCA1-Mediated Upregulation of U6atac Enhances Minor Intron Splicing and Adaptive DNA Repair

DNA damage caused by radiation, Olaparib, or Cisplatin resulted in elevated U6atac levels (**Figure 4**), indicating that U6atac levels are controlled by molecular pathways triggered by DNA damage. This regulation of U6atac might represent an adaptive mechanism where U6atac levels are upregulated to leverage the MiS to enhance DNA repair capacity under genotoxic stress. Therefore, we sought to uncover molecular pathways linking DNA damage to changes in minor intron splicing. For this, we leveraged our dual-splice reporter in PCa C4-2 cells, which were plated in an arrayed RNAi screen using a cancer driver library of 164 cancer drivers (**Figure 5A**). The sensitivity and specificity of our dual splice reporter for distinguishing between minor and major intron splicing were validated using siU6atac and siU4.2, which selectively decreased minor and major intron splicing, respectively (**Figure 5B**). Additionally, siRNA against SF3B1, which is shared by both minor and major spliceosomes, led to a reduction in both splicing machinery after 96 hours (**Figure 5B**). KD of most genes did not reduce either minor or major splicing activity by less than 45%. But a select few oncogenes, such as STAT3, XPO1, KIT, SRSF3, and MYC, selectively reduced minor intron splicing by >65% (**Figure 5C**). KD of other context-dependent oncogenic drivers, such as SF3B1, FOXA1, and KLK2, also resulted in a selective reduction of minor intron splicing by >65%. Of note, we also captured KD of DNA repair factors, such as FANCE, which led to a selective reduction in minor intron splicing (**Figure 5C**). By contrast, BRCA1 KD decreased both minor and major intron splicing by >65%. We also identified genes-albeit a much smaller number-whose KD selectively inhibited the major spliceosome (e.g., BRAF, CDKN21A). Given our discovery that DNA damage upregulates U6atac, a crucial regulator of MiS activity, we explored how DNA damage and the KD of these 164 genes would affect minor and major spliceosome activity. For this, we repeated the RNAi screen in C4-2 reporter cells irradiated with 10Gy. We discovered that KD of BRCA1, CDK4, and TRRAP, which originally reduced both minor and major intron splicing in untreated cells upon irradiation, rescued minor intron splicing but not major (**Figure 5D and Supplemental Figure 30 and 31**). In contrast, KD of MSH2, which reduced minor intron splicing in naïve cells by < 65%, further enhanced this reduction after irradiation (**Figure 5D, blue**). The fact that MSH2 is another mismatch repair gene suggests a potential radiosensitizing effect in this context (**Figure 5D, blue**). Significantly, KD of all DNA repair-associated genes exacerbated the defect in major intron splicing upon irradiation.

**Figure 5.**
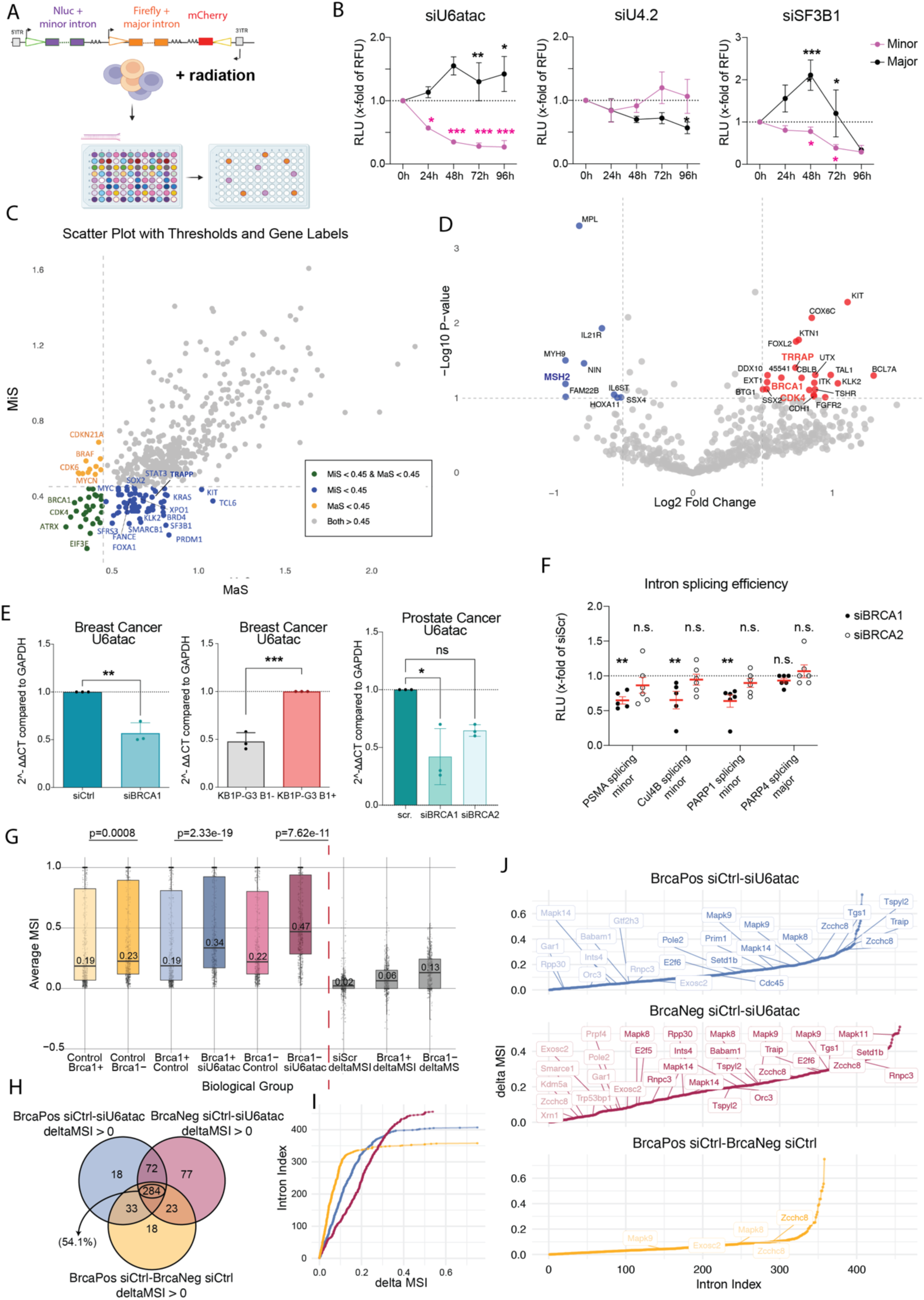
Dual reporter screen identifies BRCA1-dependent regulation of U6atac and MiS activity. (A) Schematic of dual minor/major spliceosome activity reporter. The piggyBac-based reporter contains Nanoluc (minor intron, purple) and Firefly (major intron, orange) luciferase genes, each under a separate promoter with a PEST sequence. An mCherry cassette in opposite orientation serves as an internal control. C4-2 reporter cells were treated with radiation (10 Gy), olaparib, or cisplatin, followed by reverse transfection with an siRNA cancer driver library; luciferase readouts identified drivers altering splicing activity. (B) Specificity of dual minor/major spliceosome reporter. Normalized reporter activity in C4-2 cells following siU6atac, siU4.2, or siSF3B1 (2 nM; 0–96 h). Minor (Nanoluc) and major (Firefly) activities are plotted as fold changes relative to siScrambled. Mean ± SEM, N = 3. Two-way ANOVA with Fisher’s LSD shows significant changes for siU6atac vs control (minor: 24 h *P = 0.0155; 48 h ***P = 0.0005; 72 h ***P = 0.0002; 96 h ***P = 0.0001; major: 48 h **P = 0.0026; 96 h *P = 0.0171), siU4.2 vs control (major 96 h *P = 0.0367), and siSF3B1 vs control (major 48 h ***P = 0.0004; minor 72 h *P = 0.0361; minor 96 h *P = 0.0173; major 96 h *P = 0.0252). (C) RNAi screen results in untreated cells. Scatter plot of untreated screen: each point represents a gene knocked down with three siRNAs in three technical replicates. X-axis, relative major spliceosome (MaS) activity; y-axis, relative minor spliceosome (MiS) activity. Genes with >55% luminescence decrease (MiS or MaS < 0.45) are highlighted: green (both MiS and MaS), blue (MiS only), orange (MaS only), grey (no change). Selected cancer genes are labeled. (D) RNAi screen in irradiated vs non-irradiated cells. Volcano plot comparing MiS activity effects of knockdowns in 10 Gy-irradiated vs non-irradiated cells. x-axis, log_2_ fold-change; y-axis, –log_10_ P value. Red genes: stronger effect in untreated cells (IR partially rescues MiS defect). Blue genes: stronger effect under IR (IR amplifies MiS defect). Selected genes highlighted. (E) BRCA1 alters U6atac expression. qPCR of U6atac in BCa (N = 3) and PCa (N = 3) cells. Left, BCa cells with siCtrl vs siBRCA1; middle, U6atac in KB1P-G3 B1^−^ vs KB1P-G3 B1^+^; right, 22Rv1 PCa cells with siScrambled, siBRCA1, or siBRCA2. Mean ± SEM. U6atac is significantly reduced by siBRCA1 in BCa (P = 0.0023), higher in KB1P-G3 B1^+^ vs B1^−^ (P = 0.0006), and in PCa, Friedman test P = 0.0556; Dunn’s test: modest reduction with siBRCA1 (P = 0.0495), not siBRCA2 (P = 0.3061). (F) BRCA1 alters MiS activity. Intron splicing efficiency in C4-2 reporter cells depleted of BRCA1 or BRCA2. Nanoluc reporters contained minor introns from PSMA, CUL4B, or PARP1; Firefly contained the major intron from PARP4. RLU were normalized to mCherry and expressed as fold change relative to siScrambled. Mean ± SEM, N = 5. Dunnett’s test shows BRCA1 knockdown significantly impairs minor intron splicing (PSMA, CUL4B, PARP1), with no effect on the major intron (PARP4); BRCA2 knockdown is not significant (P < 0.01, ns = not significant). (G) BRCA1 deficiency decreases MiS activity. Boxplots of average mis-splicing index (MSI) across four groups (Brca^+^/siScr, Brca^+^/siU6atac, Brca^−^/siScr, Brca^−^/siU6atac; n = 4/group). Kruskal–Wallis tests with Dunn’s post hoc and Bonferroni correction assess the effects of Brca loss and U6atac loss in Brca^+^ or Brca^−^ backgrounds. Median MSI values are indicated; P values correspond to indicated comparisons. (H) Overlap of introns with increased retention. Venn diagram showing overlap of introns with significantly increased retention (ΔMSI > 0) across BRCA-positive, BRCA-negative, and control siRNA conditions; numbers indicate unique or shared introns, with percentages of total introns in parentheses. (I) Cumulative distribution of introns with increased retention. Cumulative plots of introns with ΔMSI > 0 in expressed minor intron–containing genes (MIGs) and genes of interest (GOIs), for BRCA-positive (blue), BRCA-negative (red), and control siRNA (yellow). Intron index reflects rank order by ΔMSI. (J) Genes with intron retention show differential expression by BRCA status. Genes with increased minor intron retention across the indicated comparisons are shown. Cumulative plots (as in I) display introns with ΔMSI > 0 in MIGs and GOIs for BRCA-positive (blue), BRCA-negative (red), and control (yellow) groups.

BRCA1 is a key mediator of DNA repair, U6atac levels increase upon DNA damage, and minor spliceosome activity depends on U6atac abundance. Therefore, the selective rescue of minor intron splicing after BRCA1 knockdown in irradiated cells prompted us to investigate how BRCA1 might regulate U6atac levels. Thus, we performed siRNA-mediated BRCA1 KD in PCa and BCa cells, which resulted in a significant reduction in U6atac levels (**Figure 5E**). Moreover, BRCA1-deficient BCa cells showed lower U6atac levels than BRCA1-proficient cells (**Figure 5E, right panel, and 5 G),** confirming previous findings that irradiation increases U6atac, especially in BRCA1-proficient PCa cells (**Figure 4B**). Consistently, BRCA1 KD significantly impaired minor intron splicing (**Figure 5F**), while major intron splicing remained largely unaffected across three independent splicing reporter constructs (**Figure 5F**). Notably, the KD of BRCA2 did not decrease U6atac or minor intron splicing (**Figure 5E and F**,). Together, these findings underscore that the BRCA1-U6atac-DNA repair axis in part regulates MiS activity, and that future studies will be needed to delineate the exact players and biological pathways.

To further elaborate on the relationship between BRCA1/U6atac, we leveraged two BCa models, one with and one without BRCA1. Specifically, endogenous BRCA1 is ablated along with TP53 (KB1P-G3 B1-), and the human BRCA1 coding sequence is reintroduced to generate BRCA1-proficient cells (KB1P-G3 B1+). siScr and siU6atac-treated BRCA1-proficient and BRCA1-deficient BCa cell lines were subjected to RNAseq. Consistent with BRCA1 regulating U6atac levels, we observed elevated minor intron retention in BRCA1-deficient cells compared with BRCA1-proficient cells. In contrast, BRCA1 proficient cells treated with siU6atac showed statistically significant elevation of minor intron retention compared to siScr treated BRCA proficient cells (**Figure 5G**). In keeping with BRCA1 loss and altered minor intron splicing, we observed further enhancement of minor intron retention in BRCA1-deficient cells treated with siU6atac (**Figure 5G-J**). This enhancement of siU6atac-mediated MiS inhibition in BRCA1-deficient cells was clearly observed when we compared the delta mis-splice index (MSI) of siScr, the difference in the level of minor intron retention between siU6atac and siScr in their respective cell lines. Here, the delta MSI comparison showed that in control conditions, BRCA deficiency results in moderate MiS inhibition as reflected by elevated minor intron retention (**Figure 5G-J**). Predictably, siU6atac results in elevated MSI compared to siScr and BRCA1-proficient cells, and this effect is further exaggerated in BRCA1-deficient cells (**Figure 5G**). To further resolve intron-level effects, we restricted our analysis to introns that showed elevated retention. The measure of intron retention was determined by subtracting the MSI value of the siU6atac condition from the MSI value of the same intron in the control. For each pairwise comparison between control and siU6atac-treated samples, we ranked intron retention by delta-MSI value, from highest to lowest. This revealed progressive shifts in the number of responsive introns and the distribution of their delta-MSI values, consistent with increased intron retention upon the combined loss of Brca1 and U6atac (**Figure 5I**). Intersection of the intron IDs represented across all positive Δ-MSI series revealed a core set of 284 introns that were commonly affected (**Supplemental Table 6**). Within this core, we identified several genes implicated in DNA repair, DNA damage response, and R-loop resolution (**Figure 5J**). Notably, the largest group of unique introns (77) was detected exclusively in the siU6atac-in Brca1-deficient cells (**Figure 5J**). This set included multiple genes involved in DNA repair and R-loop regulation (**Figure 5I and J and Supplemental Table 7**). TP53BP1 was particularly striking, as its increase in minor intron retention here is consistent with our earlier findings that siU6atac reduces TP53BP1 expression (**Supplemental Figures 28 and 29**). In contrast, the set of unique introns identified in the BRCA-proficient context also contained DNA repair genes, though to a much lesser extent (**Supplemental Table 7**). These findings indicate that siU6atac treatment impairs DNA repair and triggers DNA damage in both BRCA-proficient and BRCA-deficient cells, but with a more pronounced effect in the BRCA1-deficient background. This heightened sensitivity is consistent with the newly discovered role of BRCA1 in supporting minor intron splicing, such that BRCA1 loss lowers the threshold for MiS dysfunction and unmasks a BRCA-dependent splicing liability. Collectively, these data support a model in which BRCA1 is activated upon DNA damage, leading to upregulation of U6atac and thereby enhancing minor intron splicing and contributing to an adaptive DNA repair response.

### U6atac Knockdown Potentiates PARP Inhibition and Reduces Tumor Growth Without Toxicity In Vivo

Encouraged by our *in vitro* findings, we sought to evaluate whether targeting the MiS, either as monotherapy or in combination with Olaparib, an inhibitor of PARP1, a MIG, would reduce the growth of xenografted 22Rv1 tumors in vivo. To block the MiS, we encapsulated siU6atac in polymeric nanoparticles for intraperitoneal (i.p.) injection^22^ and siScrambled served as a control (**Figure 6**). Once tumors reached a volume of 70–80 mm³, mice were treated with siU6atac or siScrambled nanoparticles, with or without Olaparib, alongside vehicle controls. Tumor volumes and body weights were measured three times a week using digital calipers until tumors exceeded 1000 mm³ or for a maximum of 6 weeks. QRT-qPCR assessed the KD efficiency of U6atac in tumors and various tissues, including prostate, breast, heart, liver, lung, muscle, spleen, kidney, testis, bone marrow, and brain. Three independent *in vivo* experiments were conducted, each varying in siU6atac dose or treatment duration. Only tumors demonstrating ≥50% U6atac KD relative to siScrambled controls were included in the final analyses of tumor growth and weight (**Supplemental Figure 32A**). Treatment with siU6atac alone significantly reduced tumor growth compared to the scrambled control (**Figure 6A and Supplemental Figure 32B**). The anti-tumor effect was further enhanced by co-treatment with Olaparib (**Figure 6B, C, and G**), with the combination group exhibiting significantly reduced tumor growth compared to the siScrambled/Olaparib control group. Additionally, siU6atac co-treated with Olaparib reduced tumor growth, such that the 200 mm3 tumor size observed in siU6atac treatment alone at 10 days was observed in co-treatment at 20 days. Thus, reflecting the enhanced reduction of proliferation of tumor cells in co-treated animals (**Figure 6C**). Throughout the treatment period, mice exhibited normal weight gain in all groups, including those receiving siU6atac and Olaparib, suggesting no overt toxicity (**Figure 6D and E**). No changes in animal behavior or histopathological abnormalities in major organs were observed, despite a significant decrease in U6atac in multiple organs in some animals, including the prostate, liver, kidney, spleen, and brain (**Figures 6H and I and Supplemental pathology reports)**. In a subset (N=8) of animals with xenografts injected with siU6atac plus Olaparib or siScrambled plus Olaparib, we performed an endpoint analysis. Specifically, within the approved ethics committee for animal experimentation, we could allow animals with xenografts to go for 6 weeks or 100mm3 tumor size, whichever one comes first. Within this endpoint, we observed that 5 of 10 siScrambled plus olaparib animals reached their endpoint. In contrast, only one reached this endpoint in the 7 animals treated with siU6atac plus Olaparib and U6atac KD >50% (**Figure 6F and G**). Collectively, we nominate U6atac KD as a viable therapeutic strategy to treat cancer in vivo that will require further toxicity studies in the future. Notably, we show that MiS inhibition, which impairs PARP1 splicing and expression, is synergistic with Olaparib, a clinically relevant PARPi prescribed to patients with BRCA-mutated PCa/BCa. Together, these data position MiS inhibition as a versatile therapeutic strategy for prostate and breast cancers, with promise for broader application across a spectrum of tumors.

**Figure 6.**
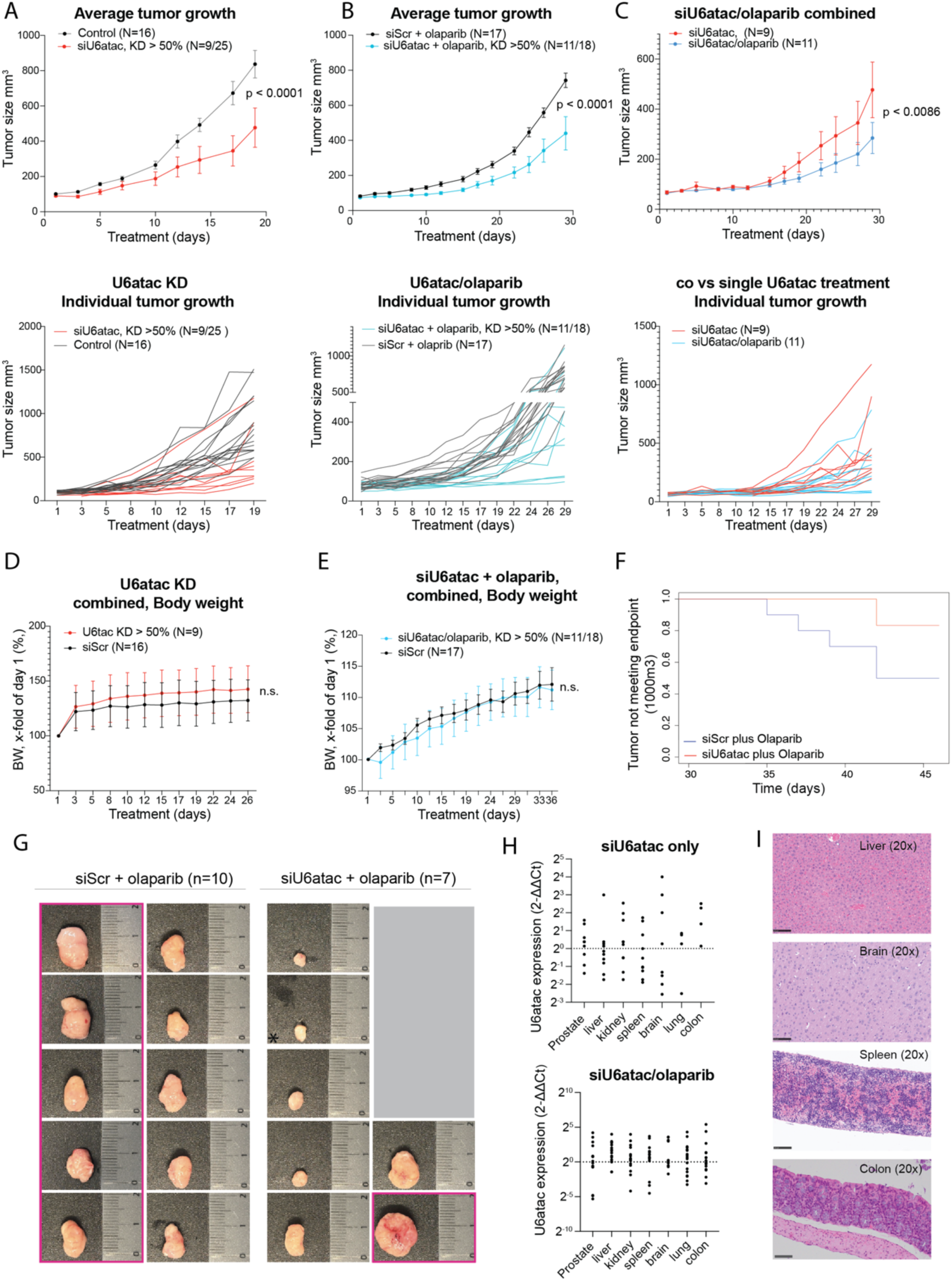
U6atac targeting suppresses tumor growth and enhances the efficacy of olaparib in vivo without overt toxicity. (A) siU6atac monotherapy slows tumor growth in vivo. Average tumor growth curves of xenografts treated with siU6atac (red; only tumors with >50% knockdown included, N = 9/25) or control siRNA (black, N = 16). Pooled from two experiments; mean ± SEM. Tumors with U6atac knockdown grew significantly more slowly at later timepoints (day 32, **P = 0.0001; final measurement, ***P < 0.0001; Šídák’s test): bottom, individual tumor growth curves. (B) siU6atac plus olaparib slows tumor growth more than olaparib alone. Average tumor growth curves of xenografts treated with siU6atac + olaparib (cyan; tumors with >50% knockdown, N = 11/18) or siScrambled + olaparib (black, N = 17). Mean ± SEM, two experiments. Co-treated tumors grew significantly more slowly at later time points (day 22, P = 0.0175; days 24, 26, 29, ***P < 0.0001; Šídák’s test; bottom: individual tumor curves). (C) siU6atac + olaparib outperforms siU6atac monotherapy. Average tumor growth curves for siU6atac alone (red, N = 9) vs siU6atac + olaparib (blue, N = 11). Mean ± SEM. Co-treatment significantly reduced growth vs siU6atac alone at later stages (day 27, P = 0.0265; day 29, **P = 0.0007; Fisher’s LSD): bottom, individual tumor curves. (D) siU6atac monotherapy does not affect body weight. Average body weight relative to day 1 in mice treated with siU6atac (red; >50% knockdown, N = 9) or siScrambled (black, N = 16). Mean ± SEM; no significant differences (ns). (E) siU6atac + olaparib does not affect body weight. Average body weight relative to day 1 in mice treated with siU6atac + olaparib (cyan; >50% knockdown, N = 11/18) or siScr + olaparib (black, N = 17). Mean ± SEM; no significant differences (ns). (F) siU6atac + olaparib delays time to tumor endpoint. Kaplan–Meier curves for time to ethical endpoint (≥1000 mm³) in siScr + olaparib vs siU6atac + olaparib–treated tumors. Curves are descriptive; statistical analysis used endpoint incidence due to low events. By study end, 5/10 siScr controls and 1/7 siU6atac + olaparib tumors reached ≥1000 mm³ (Fisher’s Exact Test, P = 0.304). (G) Representative images of excised tumors at endpoint. Tumors from siScr + olaparib (left) and siU6atac + olaparib (right)–treated mice. Pink boxes indicate tumors exceeding 1000 mm³ that triggered early sacrifice. Scale bars indicate millimeters. Data from one biological experiment followed to the endpoint. (H) siU6atac mono- and co-treatment reduce U6atac in mouse organs. Mouse U6atac expression normalized to Gapdh and shown as fold-change vs pooled control animals (log_2_ scale). Each dot, one animal; red bars, group means. Tissues: prostate, liver, kidney, spleen, brain, lung, colon. Top, siU6atac alone; bottom, siU6atac + olaparib. Data pooled from all single- and co-treatment experiments. (I) Histopathological analysis of mouse tissues after U6atac knockdown. Representative H&E-stained sections of liver, brain, spleen, and colon (20× magnification) from treated animals. No morphological abnormalities or differences were observed. Scale bars = 100 μm.

## Discussion

MiS represents a unique vulnerability of cancers as they leverage the functions executed by its targets, MIGs^1^. Indeed, MIG expression can be leveraged to distinguish normal from PCa and advanced PCa, which aligns with elevated U6atac expression during metastatic progression. Here, we show that U6atac levels can also be used to resolve TME, as reflected by U6atac high regions. Notably, through spatial transcriptomics, we discovered that a subset of cancer cells indeed leverages MiS to enhance their proliferative capacity by modifying DNA repair pathways. This idea is reinforced explicitly by DNA repair factors, including PARP1, DNA-PK, XRCC5/Ku80, and XRCC6/Ku70, which are core components of the non-homologous end joining (NHEJ) pathway (**Figure 1**). Here, we extend our study of the roles of MiS and MIGs, and DNA repair in PCa and BCa to show that, as in our previous finding, BCa progression also tracks with increasing U6atac levels and MiS activity. The therapeutic index of inhibiting the MiS is evident in our observation that siU6atac does not have a drastic effect on the proliferation of less aggressive cells, such as HS27 fibroblasts, healthy breast cells, and non-metastatic BCa cells. In contrast, metastatically aggressive TNBC and basal-like BCa cells are highly responsive.

The use of BCa cells, which are often known to engage DNA repair pathways, along with our previous report that MiS inhibition disrupted DNA repair in PCa cells, prompted us to further investigate the role of MiS in DNA repair. A CRISPR screen showed that knocking out DNA-PK, XRCC5/Ku80, and XRCC6/Ku70 markedly increased cell death upon siU6atac treatment, suggesting a synthetic lethal interaction between U6atac depletion and NHEJ disruption. Surprisingly, pulldown of the MiS-specific protein PDCD7 co-immunoprecipitated DNA repair proteins, including DNA-PK (PRKDC), PARP1, XRCC5, and XRCC6, buttressing the relationship between the MiS and DNA repair. Of course, under baseline conditions, KD of DNA repair–related genes such as *FANCE, BRCA1, CDK4*, and *TRAPP* significantly reduced minor intron splicing, positioning these genes as upstream activators of U6atac and the MiS (**Figure 5**). Interestingly, this indicates a feedback loop between BRCA1 and U6atac: while U6atac contributes to BRCA1 expression, BRCA1 also influences U6atac levels. After irradiation, KD of BRCA1, CDK4, or TRAPP no longer markedly reduced minor intron splicing, suggesting that the defect was rescued. This supports a model in which radiation activates these genes, which in turn enhance U6atac expression and MiS activity. This observation is consistent with our previous reports, which showed that the radiation-induced increase in U6atac was particularly pronounced in BRCA1-proficient models compared to BRCA-deficient ones. Loss-of-function studies further demonstrated that U6atac levels and minor intron splicing decrease upon BRCA1 (but not BRCA2) KD. In line with this, RNA sequencing revealed that minor intron retention events are markedly elevated in BRCA-deficient BCa cells relative to BRCA-proficient cells, consistent with BRCA1’s role in suppressing R-loops and promoting co-transcriptional splicing fidelity^23^. In fact, AZD7648-mediated inhibition of DNA-PK further reduced MiS activity without disrupting major splicing, suggesting a unique relationship between DNA repair machinery and the MiS. In agreement, DNA damage, which elevates U6atac levels, resulted in decreased total DNA-PK abundance while enhancing its phosphorylation, indicating activation of the DNA-PK axis. Thus, here we show for the first time how impaired capacity to repair irradiation-induced lesions following U6atac depletion might provide a new line of combination therapy to treat lethal PCa and BCa.

Crucially, we show that MiS inhibition disrupts both NHEJ and HR pathways, underscoring how MiS inhibition can be combined with DNA-damaging agents such as radiation, chemotherapy, and PARPi for enhancing efficacy and reducing dosage. Indeed, DNA damage was observed in yH2AX-positive cells treated with siU6atac at 0Gy, and this was further enhanced when siU6atac was combined with 10Gy irradiation. Elevated PARylation was observed after siU6atac treatment, indicating increased levels of DNA damage upon MiS inhibition. This finding was further bolstered by an increase in micronuclei in siU6atac treated samples with and without irradiation. Together, these findings underscore the fact that MiS inhibition causes DNA damage. The mechanism by which MiS causes DNA damage was unclear, but we now show that it can occur via R-loop formation. During transcription, newly synthesized RNA can often base-pair with DNA, forming RNA-DNA hybrid loops that are resolved upon intron splicing and the decoration of the spliced mRNA with proteins. Dysfunctional minor intron splicing leads to stalled transcriptional elongation, thereby triggering the accumulation of unspliced pre-mRNA, which increases the likelihood of RNA-DNA hybrid (R-loop) formation. Indeed, siU6atac results in increased R-loop formation, which we show is specifically reversible by RNase H1 treatment, confirming that MiS inhibition results in RNA-DNA hybrid structures. Given that we identified MIGs responsible for DNA repair and that these genes interact with the MiS, it is not inconceivable that MiS inhibition through R-loop formation causes DNA damage and that, through inhibition of MIGs and non-MIGs, it impairs the activation of the DNA damage response. Additionally, the observed upregulation of γH2AX following MiS knockdown (**Figure 2**) suggests that accumulated R-loops lead to DNA double-strand breaks, further linking minor intron splicing defects to the DNA damage response. If the damage is irreparable, this would trigger cell death to maintain genomic integrity. Indeed, overexpression of RNase H1, an enzyme that resolves R-loops, effectively rescued cells from siU6atac-induced cell death. (**Figure 2**). However, this effect could not be demonstrated in BCa cells, as RNase H1 expression was lost in both KB1P-G3 B1^+^ and MDA-436-B1^+^ cells. Given that R-loop resolution is essential for maintaining genomic integrity, these findings suggest that MiS KD disrupts splicing and acts at the transcription-repair interface, thereby enhancing tumor vulnerability to DNA-targeting therapies. In summary, U6atac KD exerts a multifaceted impact on DNA repair, not only impairing repair capacity but also generating extensive DNA damage, likely by promoting R-loop formation. This dual effect ultimately compromises genome stability and leads to cell death.

Given that we identified MIGs responsible for DNA repair and that these genes interact with the MiS, it is not inconceivable that MiS inhibition through R-loop formation causes DNA damage and that, through inhibition of MIGs and non-MIGs, it impairs the activation of the DNA damage response. Additionally, siU6atac reduced BRCA1 levels without affecting BRCA2. Although not initially considered a minor intron gene, *BRCA1* has recently been reclassified as a minor intron-like gene, which, in a context-dependent manner, can respond to the MiS. BRCA1 is crucial for HR-mediated repair of DSBs, and mutations in *BRCA1* increase sensitivity to DNA-targeting therapies, which is essential for coordinating DNA repair and maintaining genomic stability. This point is further illustrated by the experiment we conducted to compound the DNA damage caused by siU6atac with irradiation. We discovered that siU6atac impairs activation of key DNA damage response factors such as ATM, ATR, CHK1, and CHK2, which are essential for coordinating DNA repair and maintaining genomic stability, further underscoring why siU6atac, combined with irradiation, has a drastic DNA damage accumulation and disrupted DNA repair mechanisms, would result in elevated DNA damage. As a proof-of-concept, we overexpressed U6atac in both normal and cancerous prostate cells, significantly enhancing their resistance to radiation, cisplatin, and Olaparib.

Regardless of the source of DNA damage, a key feature that cancer cells evolve is the ability to repair it. Thus, it is not surprising that many therapeutic strategies exist to impair cancer cells’ ability to repair DNA. Towards this goal, Olaparib, a PARPi, is currently prescribed to treat cancers with mutations in BRCA1, CHK2, and Rad51. Here, we report that MiS inhibition synergizes with Olaparib to block cancer cell proliferation, independent of BRCA1 mutation status. The synergy here, besides MiS inhibition blocking DNA repair checkpoints, uniquely blocks PARP1, which harbors a minor intron and is essential for the repair of both single- and double-strand DNA breaks, thereby enhancing the effects of Olaparib. The versatility of MiS inhibition in synergizing with radiation and Olaparib was confirmed, as it also improved the efficacy of cisplatin in blocking cancer cell proliferation. In all, we provide robust evidence that MiS inhibition alters the DNA repair machinery and offer the first proof-of-concept for using MiS inhibition as a combination therapeutic strategy to reduce the use of widely used first-line therapies, including radiation, cisplatin, and Olaparib. This finding is highly promising, as currently, DNA-damaging agents are indispensable in cancer therapy. Still, the associated side effects are often devastating for patients, and no combined treatment for dose reduction exists presently in the clinic. Thus, we nominate MiS inhibition with its multifaceted impact on DNA repair as an ideal combination therapy candidate to enable dose reduction and minimize off-target effects. Altogether, these findings indicate that siU6atac-mediated KD of minor intron splicing induces DNA damage, likely by increasing minor intron retention of PARP1 and thereby decreasing expression and accumulation of R-loops. The buildup of R-loops arises from both impaired resolution due to reduced PARP1 and increased unspliced RNA species resulting from defective minor intron splicing. This leads to the generation of both SSBs and DSBs, while repair through the ATM and ATR-dependent pathway is compromised owing to deregulation of key DNA repair and checkpoint proteins (ATM, ATR, CHK1, CHK2, BRCA1, and 53BP1). Although DNA-PK is activated and compensates for DSB sensing, downstream repair remains defective, resulting in persistent DNA damage and enhanced radiosensitivity (**Figure 2**).

The relationship of DNA-damaging agents and MiS was recently reported by Pavlyukov et al., who showed that radiation triggers glioblastoma cells to secrete EVs enriched in spliceosomal snRNAs, which, upon uptake, confer radiation resistance to recipient glioblastoma cells^19^. Consistent with this report, we show that cancer cells treated with DNA-damaging agents upregulate U6atac levels, which are constantly under high turnover and thus constitute a crucial bottleneck for MiS activity. The fact that supernatant from cells treated with radiation resulted in enhancement of MiS activity and not major spliceosome activity suggests a unique relationship between DNA damage caused by irradiation and the enhancement of the Mis components and thereby the activity in the recipient cells. The fact that both minor and major spliceosome activity in recipient cells was enhanced when we took supernatant from cancer cells treated with Olaparib and cisplatin indicates that spliceosome activity is a crucial mediator of the evolution of therapy resistance. Of course, inhibitors for major spliceosome have been explored as therapeutics for cancer, but their toxicity is significantly higher. It is here that the small footprint of the MiS lends itself to a viable therapeutic index. We have shown that MiS is indeed critical for early embryonic development as it is required for rapidly dividing stem cells. However, in postnatal life, the use of MiS is reduced and is re-triggered in cancer. This relationship between Mis, DNA damage, and therapy resistance is further bolstered by the observation that overexpression of U6atac in normal prostate PNT1A cells conferred resistance to irradiation. Thus, it isn’t surprising that radiation-resistant mouse tumors showed high levels of U6atac compared to the matching sensitive tumors. Finally, the supernatant exchange experiment was thought to most likely deliver EVs packed with U6atac. The secretion of a large number of EVs upon irradiation is consistent with the observation that, in our spatial transcriptomics of U6atac high regions, U6atac-high tumors are enriched for genes involved in EV biogenesis (**Supplemental Figure 33).** However, when we isolated EVs from irradiated cells, recipient cells treated with these EVs showed enhanced activity of both minor and major spliceosomes, contradicting our observation with the entire supernatant. One possibility is that the supernatant, in addition to carrying EVs, may also contain factors that selectively enhance MiS over the major spliceosome.

We hypothesized that this multifaceted impact on DNA repair could be exploited to sensitize cancer cells to DNA-damaging therapies, potentially enabling dose reduction and minimizing off-target effects. This approach is particularly relevant for breast and prostate cancers, given the chronic nature of their treatment and the need for long-term survival without compromising quality of life. Our hypothesis was further supported by PARPi sensitivity screens in BCa and PCa, which showed that KO of specific MiS protein components enhances Olaparib sensitivity. *In vitro e*xperiments demonstrated a synergistic effect between siU6atac and DNA-targeting agents, as confirmed by long-term growth studies and viability assays (**Figure 3**). In fact, radioresistant mouse tumors exhibit elevated U6atac expression, suggesting that MiS plays a role in radioresistance. The thread connecting MiS inhibition to downstream effects primarily runs through elevated minor intron splicing defects, reflected in elevated minor intron retention. Here, we report that BRCA1 is a key modulator of MiS activity through an as-yet-undetermined mechanism of U6atac regulation. We show that U6atac levels are reduced when we downregulate BRCA1, leading to decreased MiS activity. Indeed, RNAseq analysis showed elevated minor intron retention in BRCA1-negative BCa cells compared to BRCA1-positive versus BRCA1-negative BCa cells, without any exogenous perturbation of the MiS. This finding suggests that BRCA1 regulates U6atac levels; as a result, when we treat BRCA-positive cells with siU6atac, the effect is less pronounced than in BRCA-negative cells treated with siU6atac. While there are many ways in which DNA damage could activate U6atac, such as p38MAPK activation^24,25^, here we suggest that BRCA1 is a crucial regulator of MiS activity by controlling U6atac levels. The exact mechanism by which BRCA1 regulates U6atac and MiS activity is currently under investigation and will be presented in future studies. In the context of radioresistance, TP53BP1 levels play a critical role, such that reductions in either its mRNA or protein levels can aid in radiosensitizing resistant tumors. Serendipitously, TP53BP1 is a MIG, and, as expected, siU6atac in BRCA-deficient cells shows elevated intron retention in TP53BP1, whereas this defect is less dramatic in BRCA1-proficient cells.

Regardless, we observed a reduction in TP53BP1 protein levels in BRCA proficient cells. Thus, we suggest that MiS inhibition can be used in tumors independent of BRCA status, thereby underscoring the significance of MiS inhibition as a novel therapeutic target, both as monotherapy and in combination with DNA-damaging agents.

These assays showed that the combined treatment of siU6atac with cisplatin, Olaparib, and/or irradiation significantly reduced cell growth and viability, yielding superior efficacy compared to each treatment alone. Notably, the combined treatment also significantly affected BRCA-proficient or AR-inactive BCa and PCa cells that do not respond to either Olaparib or siU6atac alone. This finding is highly promising as no combined treatment currently exists in the clinic that enables lower radiation doses or expands the patient groups eligible for PARP inhibitors.

Finally, we validated the MiS as a cancer vulnerability in *in vivo* mouse xenograft models, thereby demonstrating this finding for the first time (**Figure 6**). Notably, this is also the first time that siRNA targeting a MiS snRNA has been delivered via nanoparticles in an *in vivo* setting and tested in combination with Olaparib. Importantly, despite the systemic delivery method, which led to U6atac depletion in several organs, including the liver and kidneys, no signs of toxicity were observed.

Our approach demonstrated that U6atac KD significantly slowed tumor growth *in vivo* compared to scrambled controls. This effect was further enhanced when combined with Olaparib, with co-treatment resulting in significantly decreased tumor growth after several injections.

Our findings broaden the clinical relevance of DNA repair–targeted strategies by showing that KD of minor intron splicing via U6atac KD sensitizes both BRCA-proficient and -deficient prostate and breast cancers to DNA-damaging therapies. This vulnerability complements clinical advances such as the routine use of PARP inhibitors in BRCA-deficient malignancies, while extending benefit to BRCA wild-type tumors. Importantly, *in vivo* studies demonstrated tumor-selective activity with minimal toxicity, underscoring strong translational potential.

## Acknowledgments

Funding from the Prostate Cancer Foundation Challenge Award to AA, RK, and MAR.

Funding from the SAKK/Astellas GU-Oncology Award 2024 to AA.

We thank Dr Laura Brandt for Rb1/TP53 KO cells

We thank the Next Generation Sequencing Platform of the University of Bern for performing the high-throughput sequencing experiments

We thank the Core Facility Proteomics & Mass Spectrometry of the University of Bern for the mass spectrometry experiments and analysis

## Author contributions

- Conceptualisation: AA, PF
- Data Curation: AA, PF, MJ, ML, HW, NJ, SM, LL, CZ, VF, SD, SM, EP, PT
- Investigation AA, PF, MJ, ML, NJ, SM, LL, CZ, VF, SD, SM, EP, PT
- Formal analysis: AA, PF
- Computational/pathological analysis: AA, JR, CK, SdB, MG, HW, XD, KG
- Funding acquisition: AA, MR, RK
- Methodology: AA, PF, ML, AE, AAi
- Project administration: AA
- Supervision: AA, PF
- Visualization: AA, PF, KG, PT, HW, CK, MG
- Manuscript original draft: AA
- Writing-review & editing: PF, MR
- Resources: AA, PF, AE, AAi, MR, SR, RK
- Senior author: RK, SR, MR
- Correspondent: SR, MR

## STAR Methods

### Resource availability

#### Lead contact

Further information and requests for resources and reagents should be directed to and will be fulfilled by the Lead Contact, Mark Rubin

### Material availability

This study did not generate new unique reagents.

### Data and Code Availability

- The spatial transcriptomics and bulk RNAseq generated during this study will be submitted to the European Genome-phenome Archive and all datasets are publicly available as of the date of publication.
- This paper does not report original code.
- Any additional information required to reanalyze the data reported in this paper is available from the lead contact upon request.

### Experimental model and study participant details Cell and Organoid lines

HS-27 (male, ATCC, RRID:CRL-1634), PNT1A (male, Sigma Aldrich, CB_95012614), LNCaP (male, ATCC, RRID: CVCL_1379), C4-2 (male, ATCC, RRID: CRL-3314), 22Rv1 (male, ATCC, RRID: CRL-2505), PC3 (male, ATCC, RRID: CRL-3470) were maintained in RPMI medium (Gibco, A1049101), supplemented with 10% FBS (Gibco, 10270106), and 1% penicillin-streptomycin (Gibco, 11548876) on poly-L-lysine coated plates. HEK293T cells (female, ATCC, RRID: CVCL_0063), MDA-MB-231 (female, ATCC, RRID: HTB-26. Referred in the text as MDA-231), and DU145 cells (male, ATCC, RRID: CVCL_0105) were maintained in DMEM (Gibco, 31966021), supplemented with 10% FBS, and 1% penicillin-streptomycin. All cell lines were grown at 37 °C with 5% CO_2_.

KB1P-G3 (*Trp53^-/-^;Brca1^-/-^)* cells were derived from KB1P mammary tumors as previously described^26^. The KB1P-G3 BRCA1+ cell line (*Trp53^-/-^;Brca1^+/+^)* (referred in the text as KB1P-G3 B1+) was derived from the KB1P-G3 cell line, which was reconstituted with human BRCA1^21^. All mouse-derived cells were grown at 37°C in 3% O_2_ and cultured in Dulbecco’s modified Eagle medium Nutrient mixture F-12 (DMEM/F12, Gibco) supplemented with 10% Fetal Bovine Serum (FBS), penicillin/streptomycin (50 U/ml, Gibco), 5 ng/ml cholera toxin (Sigma Aldrich), insulin (5 µg/ml, Sigma Aldrich) and 5 ng/ml murine Epidermal Growth Factor (mEGF, Sigma Aldrich). The human MDA-MB-436 BRCA1-cell line (referred in the text as MDA-436 B1-was derived from a metastatic breast cancer patient and harbor a BRCA1 5396+1G > A mutation that results in loss of BRCA1 protein expression^27^. The human MDA-MB-436 BRCA1+ cell line (referred in the text as MDA-436 B1+) was derived from the MDA-MB-436 BRCA1-cell line, which was reconstituted with human BRCA1^28^. MDA-MB-436 BRCA1- and MDA-MB-436 BRCA1+ cells were grown at 21% O_2_ in Roswell Park Memorial Institute 1640 Medium (RPMI 1640; Gibco) supplemented with 10% fetal bovine serum (FBS), 2 mM L-glutamine and penicillin/streptomycin (50 U/ml, Gibco).

Isogenic cell lines from LNCaP cells were generated using CRISPR-Cas9 to knockout RB1 and/or TP53. Single cell sorting was performed using FACS and multiple clones were isolated and monitored for sufficient gene knockout by western blotting.

All cell lines were authenticated by STR analysis and regularly (every 3 month) tested for mycoplasma.

PM154 3D organoids were maintained in three-dimension according to the previously described protocol^29,30^. Briefly Advanced DMEM (Thermo Fisher Scientific, 31966047) with GlutaMAX 1x (Thermo Fisher Scientific, 35050061), HEPES 1mM (Thermo Fisher Scientific, 15630056), AA 1x (Life Technologies, 15240-062), 1% penicillin-streptomycin, B27 (Thermo Fisher Scientific,17504001), *N*-Acetylcysteine 1.25 mM (Sigma-Aldrich, A9165), Recombinant Murine EGF 50 ng/ml (PeproTech, 315-09), Human Recombinant FGF-10 20 ng/ml (Peprotech, 100-26), Recombinant Human FGF-basic 1 ng/ml (Peprotech, 100-18B), A-83-01 500 nM (Tocris, 29-391-0), SB202190 10 μM (Sigma-Aldrich, S7076), Nicotinaminde 10 mM (Sigma-Aldrich, N0636), (DiHydro) Testosterone 1 nM (Fluka, 10300), PGE2 1 μM (Tocris, 2296), Noggin conditioned media (5%) (PeproTech, 120-10C) and R-spondin conditioned media (5%) (PeproTech, 315-32). The final resuspended pellet was mixed with growth factor-reduced Matrigel (VWR, BDAA356239) in a 1:2 volume ratio. Droplets of 40 μl cell suspension/Matrigel mixture were pipetted onto each well of a six-well cell suspension culture plate (Huberlab, 7.657185) To solidify the droplets the plate was placed into a cell culture incubator at 37 °C and 5% CO_2_ for 30 min. Subsequently 3 ml of human organoid culture media was added to each well. 50 % of the media was exchanged every 3−4 day during organoid growth. Organoids were passaged as soon as they reached a size from 200 to 500 um. To this end, organoid droplets were mixed with TrypLE Express (Gibco) and placed in a water bath at 37 °C for a maximum of 5 min. The resulting cell clusters and single cells were washed and re-cultured, according to the protocol listed above.

### *In situ* validation collection

For BCa we analyzed the U6atac expression in three independent breast cancer cohorts. Tissue micro-arrays were kindly provided by Rupert Langer, Tapia Coya and Simone Münst.

### Method details

#### Spatial Transcriptomics

Detailed slide preparation has been previously described^1^. Primary visualisation antibody PanCK - epithelium (Cy3) and Syto13 - nuclei (FITC) were used to aid area of illumination (AOI) selection. Tumour compartment was selected based upon PanCK+ and Syto13+ expression, tumour microenvironment was selected based up PanCK-expression. Each TMA was cut onto a separate Leica Bond superfrost plus slide. Antigen retrieval and antibody staining was carried out on a Leica Bond automated stainer following standard protocols. The slides where then hybridized with probes from the Whole Transcription Atlas (WTA) targeting 18,000 protein coding genes. Following hybridization slides were scanned on the GeoMx DSP instrument and regions of interest (ROIs) and AOIs comprising either tumor epithelium or tumor microenvironment areas, were selected. High and low U6ATAC classified cores were selected based upon previous RNAScope labelling for U6ATAC. Following AOI selection, photocleavable oligonucleotides probes were exposed to UV-light, cleaved and the subsequent aspirate was collected into a 96-well DSP collection plate. Following rehydration with DEPC-treated water, the aspirated probes were added to the corresponding well of a new 96-well PCR plate which contained GeoMx Seq Code primers and PCR master mix. Full details of library preparation and sequencing have been previously described^31^. The sequencing runs for the GeoMX DSP were performed by Azenta.

Conversion of FASTQ files to DCC files was performed. Quality control was performed according to manufacturer’s protocols (MAN-10154-01 GeoMx-NGS Data Analysis User Manual) using the GeoMx NGS pipeline (v2.3.3.10). Non-normalized raw counts were extracted from the DSP Analysis Suite software (v.3.1.0194) for 49 AOIs (30 tumor, 19 tumor microenvironment), and the remaining data processing was performed in R (v.4.2.0). Samples with ≥60% aligned reads, and ≥1.5% of all genes exceeding the limit of quantitation (LOQ) (equating to 250 genes from the 18,000 panel) were retained (47/49 AOIs). Gene filtering was performed separately for tumor and tumor microenvironment compartments. Genes exceeding LOQ in more than 10% of the samples were retained for downstream analysis (Tumor: 14,857/18,677 genes retained, tumor microenvironment: 13,535/18,677 retained). Filtered counts were subsequently processed using DESeq2^32^, normalization using DESeq2 internal normalization method (median of ratios) was performed prior to differential gene expression analysis. Gene set enrichment analysis (GSEA) was performed and visualizations made using ClusterProfiler^33^ ggplot2 and MSigDB genesets^34^. Curated genelists comprising DNA repair genes positively associated with androgen receptor activity were based upon Polkinghorn et al. (2013)^35^, Benjamini-Hochberg was performed to account for multiple testing and an adjusted p-value of 0.05 was applied representing a 5% false discovery rate.

For spatial differential expression Genes with adjusted *p* < 0.05 and log_2_ fold change > 1.5 were classified as upregulated (104 genes), whereas genes with adjusted *p* < 0.05 and log_2_ fold change < –1.5 were classified as downregulated (16 genes). A simplified results table was generated after removing incomplete Ensembl identifiers. A volcano plot of log_2_ fold change versus –log_10_(*p*-value) was generated in R using ggplot2 and ggrepel. Significantly upregulated genes were shown in red, significantly downregulated genes in blue, non-significant genes in grey, and DNA repair genes of interest in purple. Upregulated and downregulated gene lists were used for Gene Ontology (GO) enrichment analysis with enrichGO (ClusterProfiler) using *org.Hs.eg.db*, Benjamini–Hochberg correction, and a q-value cutoff of 0.05. GO terms were simplified using a similarity cutoff of 0.6, and the top 20 terms ranked by adjusted *p* were visualized using ggplot2.

Code availability for analysis:

https://github.com/Xiaoyue-Deng/U6atac_Spatial_Transcriptomics

#### IP and Mass spectrometry analysis

For the co-immunoprecipitation (co-IP) using an anti-PDCD7 antibody, nuclear fractions of C4-2 cells were isolated using the using the Universal CoIP Kit (Actif Motif). Chromatin of the nuclear fraction was mechanically sheared using a Dounce homogenizer. Nuclear membrane and debris were pelleted by centrifugation and protein concentration of the cleared lysate was determined with the Pierce BCA Protein Assay Kit (Thermo Fisher Scientific). 2 μg of the anti-PDCD7 antibody (ab131258, Abcam) and 2 μg of rabbit IgG Isotype Control antibody (026102, Thermo Fisher Scientific) were incubated with 2 mg protein supernatant overnight at 4 °C with gentle rotation. The following morning, 30 μl of Protein G Magnetic Beads (Active Motif) were washed twice with 500 μl CoIP buffer and incubated with Antibody-containing lysate for 1h at 4 °C with gentle rotation. Bead-bound MiS complexes were washed 3 times with CoIP buffer and twice with a buffer containing 150 mM NaCl, 50 mM Tris-HCL (pH 8) and Protease and Phosphatase inhibitors. Air-dried and frozen (−20 °C) beads were subjected to mass spectrometry (MS) analysis. Proteins on the affinity pulldown beads were re-suspended in 8 M Urea/50 mM Tris-HCl pH 8, reduced 30 min at 37 °C with DTT 0.1 M/100 mM Tris-HCl pH 8, alkylated 30 min at 37 °C in the dark with IAA 0.5 M/100 mM Tris-HCl pH 8, diluted with 4 volumes of 20 mM Tris-HCl pH 8/2 mM CaCl_2_ prior to overnight digestion at room temperature with 100 ng sequencing grade trypsin (Promega). Samples were centrifuged and the magnetic beads trapped by a magnet holder in order to extract the peptides in the supernatant.

The digests were analyzed by liquid chromatography (LC)-MS/MS essentially as described elsewhere with the sole exception that the active acetonitrile gradient for peptide separation was 60min instead of 90min [Pilotto F, Douthwaite C, Diab R, Ye XQ, Al Qassab Z, Tietje C, Mounassir M, Odriozola A, Thapa A, Buijsen RAM, Lagache S, Uldry AC, Heller M, Müller S, van Roon-Mom WMC, Zuber B, Liebscher S, Saxena S. (2023) Early molecular layer interneuron hyperactivity triggers Purkinje neuron degeneration in SCA1. Neuron 111:1–21. https://doi.org/10.1016/j.neuron.2023.05.016.].

MS data was interpreted with MaxQuant (version 1.6.14.0) against a SwissProt human database (release 2020_02) with common contaminants (keratins, trypsin etc.) included using the default MaxQuant settings, allowed mass deviation for precursor ions of 10 ppm for the first search, maximum peptide mass of 5500 Da, match between runs activated with a matching time window of 0.7 min and the use of non-consecutive fractions for the different pulldowns to prevent over-fitting. Settings that differed from the default setting included: strict trypsin cleavage rule allowing for 3 missed cleavages, fixed carbamidomethylation of cysteines, variable oxidation of methionines and acetylation of protein N-termini.

Protein intensities are reported as MaxQuant’s Label Free Quantification (LFQ) values, as well as Top3 values (sum of the intensities of the three most intense peptides); for the latter, variance stabilization was used for the peptide normalization. Imputation and significance test follow essentially ref [Uldry AC, Maciel-Dominguez A, Jornod M, Buchs N, Braga-Lagache S, Brodard J, Jankovic J, Bonadies N, Heller M. Effect of Sample Transportation on the Proteome of Human Circulating Blood Extracellular Vesicles. Int J Mol Sci. 2022 Apr 19;23(9):4515. doi: 10.3390/ijms23094515. PMID: 35562906; PMCID: PMC9099550]. Missing peptide intensities for Top3 and missing protein intensities for LFQ were imputed in the following manner: if there was at most 1 evidence in one group of replicates, the missing values was drawn from a Gaussian distribution of width 0.3 centered at the sample distribution mean minus 2.5× the sample standard deviation; all other missing values were replaced by the maximum likelihood estimation method. The Benjamini and Hochberg method was further applied to correct for multiple testing. The criterion for statistically significant differential expression is that the maximum adjusted *p*-value for large fold changes is 0.05, and that this maximum decreases asymptotically to 0 as the log2 fold change of 1 is approached (with a curve parameter of one time the overall standard deviation).

For downstream analysis a contingency table was first generated to compare proteins selected by the Top3 method versus those selected by LFQ intensity. Proteins selected by both approaches were retained for further analysis. To assess nuclear enrichment, a second contingency table was created comparing protein detection across at least two replicates in *C42_Nucleus_59K* versus *C42_Nucleus_IgG*. Proteins detected in at least one nuclear dataset were carried forward to visualization. Enrichment in the 59K pulldown was determined by filtering for proteins with an adjusted p-value < 0.05 and a negative log_2_ fold change for the contrast C42_Nucleus_IgG – C42_Nucleus_59K. Gene Ontology enrichment analysis for these proteins was performed with the enrichGO function from the clusterProfiler package in R, using org.Hs.eg.db genome, the Benjamini–Hochberg method for multiple-testing correction, and a q-value cutoff of 0.05. Simplified GO terms (cutoff 0.6) were prioritized by adjusted p-value. A volcano plot of iLFQ log_2_ fold change versus –log_10_(*p*-value) was generated using **ggplot2** and **ggrepel** to visualize differential nuclear protein enrichment between *C42_Nucleus_IgG* and *C42_Nucleus_59K*.

#### Generation of U6atac and RNAseH1 overexpressing cell lines

LV290591 – RNU6ATAC Lentiviral Vector (Human) (CMV) (pLenti-GIII-CMV-GFP-2A-Puro) as well as the corresponding empty vector control were purchased from ABM. pLV[Exp]-CMV>mRnaseh1[NM_011275.3]:IRES:EGFP (VB241112-1626vyn) Lentiviral vector and corresponding empty vector were purchased from vector builder. DNA was amplified via chemical transformation of One Shot Mach1 T1 Phage-Resistant Chemically Competent E. coli cells (Invitrogen, C862003). Lentivirus was produced in HEK293T cells by transfection with the constructs, and subsequent virus containing media was used to transduce 22Rv1and MDA-123 cells. Three days post transduction the cells were subjected to puromycin selection (1 µg/mL). After the selected cells reached a confluence of 80%, they were FACS sorted for GFP positivity in the Flow Cytometry and Cell Sorting Facility (FCCS) Core facility of the University of Bern. This procedure was repeated every two months to ensure sustained plasmid expression and to prevent the emergence of plasmid-negative cell populations.

#### Drug treatments

Spliceostatin A (MedChem Express, HY-16466) and pladienolide B (MedChem Express, HY-16399) were added to the culture medium at a final concentration of 0.5 µM. Olaparib (Lubio Science, HY-10162-1G) was applied at the indicated concentrations for 96–120 h, with re-administration every 48 h. Cisplatin (Selleck Chemicals, S1166) was added to the culture medium for 96 h. Enzalutamide (Selleck Chemicals, S1250) was applied at a final concentration of 20 µM for 96 h.

DNA-PK inhibitor AZD7648 (Selleck Chemicals, S8843) was applied at final concentration of 10uM for 96h.

#### Cell irradiation

Irradiation was performed with an X-RAD SmART irradiator (Precision X-Ray) at the doses indicated in the corresponding figures.

#### Cell transfection and siRNA-mediated knock-down

##### Cells

siRNA SMARTpool siRNAs against *BRCA1* (Ambion Silencer Select Predesigned siBRCA1, ThermoFisher, 4392420) and *BRCA2* (Ambion Silencer Select Predesigned siBRCA1, ThermoFisher, 4390824), siRNA were purchased from ThermoFisher. siRNA against DNA-PK (on-target plus, smart pool L-005030-00-0005) was purchased from Dharmacon. siRNAs against RNU6atac (4390771, nn264646) and the Silencer Select Negative Control (4390843) were purchased from Thermo Fisher Scientific and siRNA against mouse U6atac was purchased from Ambion (4390827). Transfection was performed for the respective timepoints on attached cells using the Lipofectamine™ RNAiMAX Transfection Reagent (Thermo Fisher Scientific, 13778150) to the proportions of 16pmol of 20 μM siRNA per well.

##### Organoids

Before transfection organoids were cultured for 2-3 weeks in human organoid growth medium. Media was removed and organoids were first mechanically dissociated. To obtain single cells organoids were trypsinized in 1ml TriplE (Thermo Fisher Scientific, 12605036) for 15-18 minutes at 37C. The reaction was stopped with 1ml growth media and cells were spun for 5 minutes at 300g. Subsequently the cells were strained and counted. Per condition one million cells were plated in a 6 well. Lipofectamine™ RNAiMAX complexes were prepared according to the standard Lipofectamine™ RNAiMAX protocol. In short, 5ul of RNAiMAX reagent and 40 nM of siRNA plus 10% FBS were each diluted in 125 ul Opti-MEMH medium. Both mixes were pooled and incubated for 10 minutes before the siRNA-reagent complex was added to the cells. Cell/siRNA mix was centrifuged at 600 g at 32C for 60 min and then incubated over night at 37C. The next day cells were resuspended and collected by centrifugation (300g, 5min, RT). The pellet was resuspended in 280 ul Matrigel and the mix was separated into 7 drops that were added into a 6 well. Organoids were grown in human organoid media for 96h (CTG assay) or seven days (cell counting assay).

#### RNA extraction from cells and qRT-PCR

RNA from cultured cells was isolated using the ReliaPrep™ miRNA Cell and Tissue Miniprep System (Promega, Z6212). For extracellular vesicles (EVs), RNA was extracted with the ReliaPrep™ FFPE Total RNA Miniprep System (Promega, Z1002). For FFPE samples, tumor blocks were sectioned into 10 µm slices using a Leica Histocore Multicut microtome, and RNA was isolated from eight sections with the ReliaPrep™ FFPE Total RNA Miniprep System (Promega, Z1002). For fresh-frozen mouse tumors, tissue pieces were disrupted in lysis buffer with a stainless steel bead (Qiagen, REF 69989) using a Qiagen TissueLyser system, and RNA was subsequently isolated with the ReliaPrep™ miRNA Cell and Tissue Miniprep System (Promega, Z6212) according to the manufacturer’s instructions.

Synthesis of complementary DNAs (cDNAs) using FIREScript RT cDNA Synthesis Kit (Solis BioDyne, 06-15-00200) and real-time reverse transcription PCR (RT-PCR) assays using HOT FIREPol EvaGreen qPCR Mix Plus (Solis BioDyne, 08-24-00020) were performed using and applying the manufacturer protocols. Quantitative real-time PCR was performed on the ViiA 7 system (Applied Biosystems). All quantitative real-time PCR assays were carried out using at least three technical replicates. Relative quantification of quantitative real-time PCR data used GAPDH, ACTB as housekeeping genes. Primer sequences are listed in **Supplemental Table 8**.

#### Growth Assays

To assess the growth and survival upon treatment with olaparib, cisplatin or irradiation, KB1P-G3 B1+, KB1P-G3, PNT1A (overexpressing U6atac or empty vector control), C4-2 and DU145 cells were seeded in 6-well plates in the following densities: 2,000 cells/well (KB1P-G3 B1+) or 4,000 cells/well (KB1P-G3) or 8,000 cells/well (DU145, PNT1A and C4-2). All the cells were treated with the mentioned drug or irradiation at the indicated dosages after 24 h. IR-treated cells were subsequently exposed to repeated irradiation on day 2 and 3. The treatment of cells with DMSO or indicated concentrations of olaparib lasted for the whole duration of the experiment. The medium with DMSO or olaparib was refreshed twice a week. The control, DMSO-treated plates were fixed 7 days after seeding, the olaparib-treated plates were fixed after 10 days. For the cisplatin treatment, cells were plated 24h prior addition of cisplatin-containing media. After 24 h, medium was refreshed and all the plates were fixed after 7 days. The fixation was done with 4% formalin and the surviving colonies stained with 0.1% crystal violet. The cell survival and growth were analyzed in an automated manner using the ImageJ ColonyArea plugin growth assays in isogenic cell lines *cells were transfected with siRNA against U6atac of negative control as previously described. After 24h,* a total of 5,000 cells per well were seeded in clear 96-well plates. After overnight incubation, 1µM Olaparib was added. Plates were read in an IncuCyte S3. Every 6h, phase object confluence (percentage area) for cell growth was measured.

#### Bulk RNA sequencing

Total RNA was extracted from MDA-436 and KB1P-G3 B1+ and B-cells treated for 96h with siU6atac or siScrambled. The recommended DNase treatment was included. The quantity and quality of the extracted RNA was assessed using a Thermo Fisher Scientific Qubit 4.0 fluorometer with the Qubit RNA BR Assay Kit (Thermo Fisher Scientific, Q10211) and an Advanced Analytical Fragment Analyzer System using a Fragment Analyzer RNA Kit (Agilent, DNF-471), respectively. Thereafter cDNA libraries were generated using an Illumina Stranded Total RNA Prep, Ligation with Ribo-Zero Plus (Illumina, 20040529) in combination with IDT for Illumina RNA UD Indexes Sets A and B (Illumina, 20040553 and 20040554, respectively). The Illumina protocol was followed exactly with the recommended input of 100 ng total RNA. The quantity and quality of the generated NGS libraries were evaluated using a Thermo Fisher Scientific Qubit 4.0 fluorometer with the Qubit dsDNA HS Assay Kit (Thermo Fisher Scientific, Q32854) and an Advanced Analytical Fragment Analyzer System using a Fragment Analyzer NGS Fragment Kit (Agilent, DNF-473), respectively. As a further quality control step, prior to NovaSeq 6000 sequencing, the pooled cDNA library pool underwent paired end sequencing using iSeq 100 i1 reagent v2, 300 cycles (illumina, 20040760) on an iSeq 100 sequencer. The library pool was re-pooled to ensure an equal number of reads/library and then paired end sequenced using a NovaSeq 6000 S4 reagent kits v1.5, 300 cycles (Illumina, 20028312) on an Illumina NovaSeq 6000 instrument. The quality of the sequencing runs was assessed using Illumina Sequencing Analysis Viewer (Illumina version 2.4.7) and all base call files were demultiplexed and converted into FASTQ files using Illumina bcl2fastq conversion software v2.20. The average number of reads/ libraries was 82 million. The RNA quality-control assessments, generation of libraries and sequencing runs were performed at the Next Generation Sequencing Platform, University of Bern, Switzerland.

##### Breast Cancer RNAseq, Retention Strategy

Our analysis of intron retention included four biological groups defined by the presence/absence of Brca and/or U6atac and consisting of n = 4 replicates per group (Brca+/siScr, Brca+/siU6atac, Brca-/siScr, Brca-/siU6atac). Three pairwise comparisons were performed to assess the effect of (1) Brca loss, (2) U6atac loss in the presence of Brca, and (3) U6atac loss in the absence of Brca. Paired-end, total RNA sequencing reads from each sample were mapped to GRCm38.99 using Hisat2 and uniquely mapped reads were used downstream for the assessment of minor, minor-like, and hybrid intron retention. Intron classification, coordinates, and transcript- and gene-level associations for these introns were sourced from REF Olthof 2024. Intron retention events in expressed minor intron-containing genes (MIGs) or “genes of interests” (GOIs, i.e., minor-like or hybrid intron-containing genes) were identified using a custom bioinformatics approach described in (REFs Olthof 2019; 2024). Briefly, we assessed intron retention events that were supported in at least one replicate of either biological group by: ≥4 reads spanning exon–intron junctions, with at least one read supporting the 5′ splice site and the 3′ splice site and read coverage extending across >95% of the intron body. The mis-splicing index (MSI) of introns that passed this filtering criteria and resided in genes that were expressed at or above a 1.0-TPM threshold in at least one biological group, as determined by IsoEM2, were considered in subsequent analyses. For each pairwise comparison, the global differences between groups were assessed using a Kruskal-Wallis rank-sum test with post-hoc Dunn’s test with Bonferroni correction. To visualize the distribution of introns with increased retention, introns with a delta-MSI > 0 from each pairwise comparison were collected and ranked in ascending order of delta-MSI (from 1 to n)

#### *In situ* validation collection

We analyzed the U6ATAC expression in three independent breast cancer cohorts comprising 195 patients. Of them, 65% (N=125), 4% (N=8), 4% (N=8) and 27% (N=54) were classified as luminal A or B, Her2-positive, basal-like and triple negative, respectively. Tissue micro-arrays were kindly provided by Rupert Langer, Tapia Coya and Simone Münster.

#### U6atac *in situ* hybridization

mRNA ISH was performed by automated staining using Bond RX (Leica Biosystems) and Basescope® technology (Advanced Cell Diagnostics, Hayward, CA, USA). All slides were dewaxed in Bond dewax solution (product code AR9222, Leica Biosystems) and heat-induced epitope retrieval at pH 9 in Tris buffer based (code AR9640, Leica Biosystems) for 15 min at 95° and Protease treatment for 5 min. The following probes from RNAscope 2.5 LS (Advanced Cell Diagnostics) were used: BaseScope™ LS Probe - BA-Hs-RNU6ATAC-1zz-st-C1ref 1039918, PPIB-1zz ref 710178 and DapB-1zz ref 701028, were used as positive and negative control respectively. Probe efficiency was tested using U6atac overexpressing C4-2 cells (5 million) of which 50% were treated with siU6atac RNA.

All probes were incubated at 37° for 120 min. Basescope™ 2.5 LS Assay (Ref 323600, Advanced Cell Diagnostics) was used as pre-amplification system. Subsequent the reaction was visualized using Fast red as red chromogen (Bond polymer Refine Red detection, Leica Biosystems, Ref DS9390) for 20 min. Finally, the samples were counterstained with Haematoxylin, air dried and mounted with Aquatex (Merck). Slides were scanned and photographed using Pannoramic 250 (3DHistech). U6atac intensity was scored manually by a pathologist (Charlotte Komarek and Antonio Rodriguez-Calero) blinded to the clinical data, using the digital online TMA scoring tool Scorenado^36^ (University of Bern, Switzerland) especially developed for TMA scoring on de-arrayed spots.

U6ATAC score was calculated by multiplying the percentage of positive cells (in 5% steps from 0% up to 100%) by the intensity (low, 1; intermediate, 2 or high, 3). When more than one spot was assessable for the same tumor type and same patient, an average of the obtained scores for the different spots was calculated. The final score ranging from 0-300 was categorized in three discrete groups: <= 100, low; <= 200, intermediate and > 200, high. Further statistical analysis was performed using “R” software. When testing a difference in ratios of categories between groups two tests are generally used: Fisher’s exact test and Chi-squared (χ²) test. Paired comparisons are based on McNemars test.

#### Immunofluorescence

Cells were seeded on coverslips in 24-well plates 3 days prior the experiment. To analyze parylation, Rloops and ɣH2AX foci formation, DNA damage was induced by γ-irradiation (10 Gy) 24h or at the indicated time point prior to fixation. Before fixation, cells were pre-extracted for 5 min on ice in 10 mM PIPES pH6.8, 300 mM sucrose, 50 mM NaCl, 3 mM EDTA, 0.5% Triton X-100. Fixed cells were washed with PBS and permeabilized for 20’ in 0.2% (v/v) Triton X-100/PBS. Next, slides were washed three times with 0.2% Tween-20/PBS and blocked with staining buffer (PBS, BSA (2% w/v), glycine (0.15% w/v), Triton X-100 (0.1% v/v)) for 1h at RT. Incubation with the primary rabbit polyclonal anti-pan-ADP-ribose binding reagent (#MABE1016, Millipore), rabbit phospho-histone H2AX (ser139) (also mentioned as anti-ɣH2AX) (#2577, Cell Signaling), mouse anti-DNA-RNA hybrid (S9.6) (#ENH001, Kerafast) diluted 1:1000 in staining buffer was carried out for 1h at RT. Slides were then washed four times for 5’ with 0.2% (v/v) PBS-Tween-20 and then incubated with Goat anti-mouse IgG (H+L) Cross-Absorbed Secondary Antibody, Texas Red-X (# T-862, Thermo Scientific), with Goat anti-Rabbit IgG (H+L) Cross-Adsorbed Secondary Antibody Alexa Fluor 647 (Thermo Scientific, Cat#A21244) or with Alexa Fluor 488 goat anti-rabbit IgG (H+L) (Thermo Scientific, Cat#A11034) diluted 1:2500 in staining buffer for 1h at RT. Slides were washed three times for 5’ with 0.2% PBS-Tween-20, once with PBS and then mounted with Duolink *In Situ* mounting medium with DAPI (#DUO82040, Sigma Aldrich). Z-stack fluorescent images were acquired using the DeltaVision Elite widefield microscope (GE Healthcare Life Sciences). Multiple fields of view were imaged per sample with Olympus 100X/1.40, UPLS Apo, UIS2, 1-U2B836 objective and sCMOS camera at the resolution 2048 x 2048 pixels. Deconvolution of the acquired images was performed by the softWoRx DeltaVision software. Image analysis was performed using Fiji image processing package of ImageJ. Briefly, all nuclei were detected by the “analyze particles” command and all the foci per nucleus were counted with the “find maxima” command. Data were plotted with Prism software.

#### R-loop quantification

##### PCa

Quantitative Rloop and RNase H1 cell expression analysis was performed on fluorescent images (TIF) with Visiopharm software (2025.02 x64. Visiopharm, Hørsholm, Denmark). The following workflow was applied: 1) Cell segmentation: Deep learning (DL) (U-Net) classification for individual DAPI (red) stained nucleus detection followed by nuclear dilation by 50 pixels to define the cell body. 2) Cell classification: Based on the presence of Rloop (blue; cell defined as positive if a mean blue pixel intensity of 300 or more was present in at least 5% of the cell) and RNase H1 (green; cell defined as positive if a mean green pixel intensity of 150 or more was present in at least 30% of the cell) staining, the cells were categorized into 4 groups: double negative; Rloop positive/RNase H1 negative; RNase H1 positive/Rloop negative; double positive. 3) The output was defined as follows: absolute and relative (percentage) cell counts (for each cell category); mean Rloop cell expression (measured as blue pixel intensity) of Rloop positive/RNase H1 negative and double positive cells.

##### BCa

As for PCa, Visiopharm software was used for the quantitative assessment of fluorescent images (TIF). Specifically, nuclear Rloop immunolabeling was quantified. The analytical workflow and output parameters established for the PCa samples were subsequently optimized for the BCa cell line to account for distinct R-loop expression patterns. In contrast to the PCa cell line, which exhibited a rather diffuse, finely granular nuclear and cytoplasmic signal, the BCa cell line showed one to three well-defined nuclear foci per cell, each approximately 1.5–3 µm in diameter. The following workflow was applied: 1) Cell segmentation: Deep learning (DL) (U-Net) classification for individual DAPI (blue) stained nucleus detection. 2) Detection and labeling of individual nuclear Rloop foci (labelled green) (DL U-Net classification). 3) The output was defined as follows: total cell count; total count nuclear Rloop foci; count of Rloop foci per individual nucleus (range 0 to 3); relative (percentage) counts of Rloop foci compared to total nuclear count.

#### Analysis of micronuclei formation

Cells were seeded onto coverslips in 24-well plates and transfected with the indicated siRNAs 24 h later. After 48 h of treatment, the cells were irradiated with 10 Gy. At 24 h post-irradiation, cells were washed with PBS and fixed with 4% (v/v) PFA/PBS for 20 mins in RT. Cells were then washed 3 times in 0.2% (v/v) PBS-Tween-20 and permeabilized for 20 mins in 0.2% (v/v) Triton X-100/PBS. Then, slides were washed 3 times with PBS and blocked for 30 mins at RT in staining buffer (PBS, BSA (2%), glycine (0.15%), Triton X-100 (0.1%)). Slides were counterstained with DAPI (1:50,000 dilution, Cat# D1306, Life Technologies) and washed 5 times more with PBS before mounting in Fluorescence mounting medium (Cat# S3023, Dako). Z-stack images were acquired using the DeltaVision Elite widefield microscope (GE Healthcare Life Sciences). Multiple fields of view were imaged per sample with Olympus 100X/1.40, UPLS Apo, UIS2, 1-U2B836 objective and sCMOS camera. Frequency of micronuclei positive cells was analyzed manually in FIJI image processing package of ImageJ (1.8.0). Scatterplot graphs of % cells with micronuclei were drawn using GraphPad Prism 9. Error bars represent the median and 95% (confidence interval) CI. *P*-values were calculated via 2-way ANOVA with Tukey’s post-test.

#### Immunoblotting

Cells were lysed in GST-Fish buffer (10 % (v/v) Glycerol, 50 mM Tris-HCl pH7,4, 100 mM NaCl, 1 % (v/v) Nonidet P-40, 2 mM MgCl_2_, 1 mM PMSF) with freshly added protease and phosphatase inhibitors. Total protein concentration was measured using the Pierce BCA Protein Assay Kit (Thermo Fisher Scientific). 50 ug protein samples were resolved 4-15% and /or 8% Mini-Protean TGX gels (BioRad, 456-1084 and Sure-page Bis-Tris gels (Witec) with Tris-MOPS-SDS Running Buffer) in SDS-PAGE and transferred to nitrocellulose membranes using the iBlot2 system (Thermo Fisher, IB23001). Proteins larger than 100KDa were transferred using the wet-blot technique using a Towbin buffer containing 20% methanol. Blots were blocked for 1 hour at room temperature 5% BSA/TBST and incubated overnight at 4 °C with primary antibodies (**Supplemental Table 9**) which were dissolved in 5% BSA/TBST buffer. After 3 washes, the membrane was incubated with secondary antibody conjugated to horseradish peroxidase for 1 h at room temperature. After 3 washes, signal was visualized by chemiluminescence using the Luminata Forte substrate (Thermo Fisher Scientific, WBLUF0100) for strong antibodies and WesternBright Sirius-HRP Substrate (Witec AG, K-12043-D10) for weak antibodies. Images were acquired with the FUSION FX7 EDGE Imaging System (Witec AG).

#### Synergy assays

KB1P-G3 B1+, 22Rv1, and Du145 cells were seeded in 96-well plates at 5,000 cells per well. After 24 h, cells were transfected with 0, 4, 8, 16, or 32 nM siU6atac or a scrambled siRNA control. Twenty-four hours post-transfection, each condition was treated with 0, 0.5, 1, or 2 µM Olaparib, with Olaparib re-applied 72 h and 120h after transfection. Cell viability was measured 6 days after siRNA transfection using the CellTiter-Glo® Luminescent Cell Viability Assay (Promega, G9243) on a Tecan Infinite M200 PRO reader, following the manufacturer’s protocol. Each experiment included three technical replicates per condition and was repeated in three to four independent biological replicates. Expected drug combination responses were calculated using the Highest Single Agent (HSA) reference model in SynergyFinder^37^. Deviations between observed and expected values were reported as synergy (positive) or antagonism (negative). We applied an HSA synergy threshold of ±5 instead of the SynergyFinder default of ±10. Our experiments used triplicate measurements per cell line, low variability (CV <15%), and a focused compound set, reducing measurement noise and increasing sensitivity to subtle, biologically relevant deviations from additivity. Prior studies have shown that HSA scores follow a near-normal distribution around zero and that synergy scoring remains robust in low-variability setups, supporting the use of a ±5 cutoff in this context^38–40^

#### EV isolation

Extracellular vesicles (EVs) were isolated from serum-free, conditioned cell culture media (CCM). The CCM was first cleared of floating cells and large cell debris by differential centrifugation (300 x g for 10 min, and 2’000 x g for 20 min, at 4 °C). The EVs in the CCM were concentrated by ultracentrifugation (100’000 x g, 90 min, 4 °C) and subsequently purified by size exclusion chromatography (SEC) with an IZON® automated fraction collector equipped with original qEV 2 columns. The eluted fractions containing EVs were further concentrated down to 0.5-1 mL using Amicon® ultrafiltration systems (regenerated cellulose, 10 kDa molecular weight cut-off). The yield, zeta potential and size distribution profiles of the purified EVs were quantified by nanoparticle tracking analysis (NTA), using a ZetaView system (Particle Metrix, software version: 8.05.05 SP; 488 nm laser). The instrument’s camera sensitivity and shutter values were set to 80 and 100 ms-1, respectively. The videos were acquired at 11 separate positions, and ≥200 particles had to be traced for at least 15 consecutive frames.

##### EV follow up

To assess the impact of EVs on splicing, 20,000 cells were seeded per well in white 96-well plates. After 24 h, the culture medium was replaced with a mixture of 50% exosome-free FBS medium and 50% EVs resuspended in PBS. As a control, cells were treated with 50% medium and 50% PBS alone. Splicing activity was measured at 2, 4, 8, 16, and 24 h after treatment.

#### Reporter assays

##### C4-2 prostate cancer (PCa)

were generated by stable genomic integration of a PiggyBac (PB) plasmid to simultaneously monitor minor and major spliceosome activity (**Figure 5A**). The minor spliceosome reporter consisted of a Nanoluc luciferase (Nluc) coding sequence interrupted by minor introns from the CUL4B gene, selected to specifically report minor intron splicing. The major spliceosome reporter consisted of a Firefly luciferase coding sequence interrupted by a major intron from the PARP4 gene. Intron selection was based on prior siU6atac data and met the following criteria: (1) high retention without aberrant alternative splicing upon minor spliceosome knock down; (2) intron length of ~200–600 bp; (3) biological relevance of the host gene; (4) no cross-reactivity to knock down of the other spliceosome; and (5) Position Weight Matrix scores for the 5′ splice site and branch point >90%. Both luciferase reporters contained a PEST sequence to promote rapid protein degradation, ensuring that luminescence reflected newly synthesized protein from successfully spliced transcripts. Reporter transcription was driven by EF1A (Nluc) or CAG (Firefly) promoters of comparable strength to minimize transcriptional variation. An mCherry cassette at the 3′ end of the PB plasmid served as a marker for transcriptional activity and integration copy number. Plasmids were synthesized by VectorBuilder.

For stable integration, C4-2 cells were co-transfected with 0.5 µg circular transposon plasmid and 0.5 µg helper PBase transposase using Lipofectamine 2000 (Invitrogen) according to the manufacturer’s protocol. Cells were seeded at 1.5 × 10⁵ per well in a 24-well plate one day prior to transfection. One-week post-transfection, mCherry-positive cells were isolated by FACS to establish stable reporter lines.

##### Splicing activity measurements

To quantify minor and major intron splicing activity, cells were seeded in white 96-well plates (Huberlab, 7.655 098) at 8,000 cells per well and treated according to assay conditions. Luciferase activity was measured using the Nano-Glo® Dual-Luciferase® Reporter Assay System (Promega, N1620) following the manufacturer’s instructions. Briefly, cells were equilibrated to room temperature, and mCherry fluorescence was recorded. Firefly luciferase substrate (100 µL) was then added to each well, plates were shaken for 3 min, and luminescence was measured. Subsequently, NanoLuc substrate (100 µL) was added, plates were shaken for 3 min, incubated at room temperature for 10 min, and NanoLuc luminescence was recorded using a Tecan Infinite M200 PRO plate reader. Firefly and NanoLuc signals were normalized to mCherry fluorescence, and results were expressed as fold change relative to the corresponding reference control.

##### DSB-Spectrum reporter

DSB-Spectrum_V1 reporter assay was performed as previously described with few adaptations (Van De Kooij et al., 2022). HEK293FT cell containing DSB-Spectrum_V1 were a kind gift from Michael B. Yaffe and Haico van Attikum. Briefly, 10,000 cells were seeded per well in a 96-well plate on day 0, cells were seeded in quadruplicates for each siRNA. On day 1, transfection with the indicated siRNA was performed with Lipofectamine RNAiMAX (Thermo Scientific, #13778150) according to manufacturer’s protocol. On day 2, 24 h after transfection, media was refreshed. 6h later, duplicates of target cells were transfected using Lipofectamine 2000 (Thermo Scientific, #11668030) with pX459-Cas9-sgRNA-iRFP constructs containing either an AAVS1 sgRNA (control) or a sgRNA targeting DSB-Spectrum BFP sequence. On day 6, cells were trypsinized, resuspended in 1%BSA in PBS and subjected to flow cytometry with the CytoFLEX S (Beckman Coulter) for RFP, GFP and BFP intensity. Data was analyzed with FlowJo software. Fractions of HR positive (GFP+) or NHEJ positive (BFP+) population were calculated from cells transfected with the sgRNA construct (RFP+).

#### RNAi screen

The RNAi screen was performed using the Silencer™ Select Human Cancer Genome siRNA Library (Thermo Fisher Scientific, A30142). Lyophilized siRNAs (0.25 nmol per well) were reconstituted with 25 µL nuclease-free water to 10 µM, diluted 1:10 to 1 µM, and aliquoted into 50 µL working stocks for storage at −20 °C. Based on pilot experiments with siGAPDH (Thermo Fisher Scientific, AM4605), a final concentration of 25 nM siRNA (5 pmol per well from a 1 µM stock) was selected, achieving ~70% GAPDH knockdown.

For screening, C4-2 CUL4B reporter cells were reverse transfected in 96-well plates with or without irradiation (10 Gy, 15 min post-irradiation). For each well, 5 µL siRNA (1 µM) was diluted in 20 µL Opti-MEM (Gibco), mixed gently, combined with 0.3 µL Lipofectamine™ RNAiMAX (Thermo Fisher Scientific), and incubated for 20 min at room temperature. A suspension of 10,000 cells in 150 µL antibiotic-free growth medium was then added to each well. Plates were incubated for 72 h at 37 °C, and luciferase activity was measured using the NanoLuc assay protocol described above. Each plate included siScrambled controls for normalization.

##### RNAi screen data analysis

RNAi screen data were analyzed as described for the splicing reporter assay. Technical replicates (n = 3) were averaged and normalized to scrambled siRNA controls to calculate fold change. Genes were classified for scatter plot visualization as: (blue) minor splicing reduced by >65% only, (orange) major splicing reduced by >65% only, (green) both reduced by >65%, or (grey) neither reduced by >65%. To assess irradiation effects on upstream minor spliceosome regulation, parallel screens were performed in untreated and irradiated reporter cells (10 Gy, 15 min post-irradiation). For each gene, log_2_ fold change (irradiated/untreated) and –log_10_ p-values were computed in RStudio (R 4.4.2), and paired t-tests were applied across biological replicates. Unless otherwise stated, thresholds were set to |log_2_FC| ≥ 0.5 and p < 0.1. Genes were categorized as *Upregulated*, *Downregulated*, or *Not Significant*, and volcano plots were generated using ggplot2 and ggrepel in R.

#### Cell growth experiments

##### Viability

Cells were treated according to assay conditions. Cell viability was determined with a Tecan Infinite M200PRO reader using the CellTiter-Glo® Luminescent Cell Viability Assay according to manufacturer’s directions (Promega, G9243). Viability values were calculated as *x*-fold of control cells.

##### Cell confluence

Cells were treated according to assay conditions. And confluence was determined using the Incucyte S3 instrument and the IncuCyte S3 2018B software (Essen Bioscience, Germany). Values were calculated as x-fold of timepoint 0 and then as fold-change in confluency as compared to siScrambled or DMSO treated control cells.

#### Xenograft experiments

##### Mouse models

NOD *scid* gamma (NSG) mice (5-6 weeks old; purchased from Charles River Laboratories, Sulzfeld, Germany) were maintained at a controlled ambient temperature of 20°C ± 2°C in a constant humidified atmosphere of 50% ± 10% on a 12 h automatic light/dark cycle.

Animals were housed in ventilated cages and given standard presterilized rodent chow and water ad libitum. Mice were allowed to acclimate for 2 weeks before being used in experiments. Only male mice were used in this study. All animal experiments were performed in accordance with national regulations and approved by the Cantonal Veterinary Ethical Committee, Switzerland (licenses no. BE46/21 and no. BE35/24).

<u>Preparation of P10Y/siRNA complexes</u> the 10 kDa linear polyethylenimine derivative (LP10Y) modified with approx. 33% tyrosine was synthesized as described previously^22^. The LP10Y/siRNA polyplexes were assembled based on a polymer/siRNA mass ratio of 2.5. Per injection, either 10 µg or 20 µg siRNA (200 µM stock concentration) were diluted in TH buffer (10% (w/v) trehalose, 20 mM HEPES pH 7.4) to a final volume of 75 µL. In a separate tube, 25 µg or 50 µg LP10Y (10 mg/mL stock concentration) were diluted in 75 µL TH buffer. The siRNA dilution was added into the polymer dilution and vigorously mixed by pipetting and incubated for 30 minutes at room temperature. Finally, 150 µl of the polyplexes were intraperitoneally injected into the mice.

<u>Tumor xenografts and treatment at the age of 8 weeks,</u> male NSG mice were subcutaneously injected in the flank with 22Rv1 prostate cancer cells (2 × 10⁶ cells/mouse). 22Rv1cells were previously mixed with Matrigel (1:1) before injection. A total amount of 100ul was injected. The procedure was performed under controlled anesthesia of Isoflurane.

Tumor growth was monitored, and mice were randomized into treatment or control groups (n = 8–10 tumors/group) when tumors reached ~60–80 mm³. P10Y/siRNA polyplexes (150 µL containing 10 or 20 µg siRNA) were administered intraperitoneally (i.p.) three times per week. Olaparib was freshly prepared on the day of injection by dissolving in 10% DMSO, diluting with PBS containing 10% hydroxypropyl-β-cyclodextrin (HPβCD) to 5 mg/mL, and administered i.p. 5 times per week at 50 mg/kg (vehicle: 10% HPβCD/PBS with 10% DMSO). Injection sites for siRNA and olaparib were alternated between administrations.

<u>Monitoring and endpoint criteria</u> Tumor dimensions were measured 3 times per week with digital caliper and tumor volume was calculated using the formula 4/3pi*((sqrt(L*W))/2)3, where L is the minor tumor axis and W is the major tumor axis. Body weight and animal behavior were recorded throughout the treatment period to assess biocompatibility. At endpoint, mice were euthanized, and tumors plus organs were harvested for RNA extraction and histopathological analysis by immunohistochemistry. Endpoint criteria included tumor volume ≥ 1000 mm³ and /or completion of the planned treatment duration.

Experimental design: Three independent xenograft experiments were performed:

- Experiment 1: siRNA monotherapy (10 µg and 20 µg) for 3 weeks.
- Experiment 2: siRNA monotherapy (20 µg) and combination therapy with Olaparib for 4 weeks.
- Experiment 3: siRNA (10 µg) plus Olaparib for up to 6 weeks.

Only tumors with ≥ 50% U6atac knockdown were included in the final analyses.

#### Genome-wide CRISPR/Cas9 screens analysis

Genome-wide CRISPR/Cas9 screens were previously published^15,16^. For the analysis in this study, quality control was performed using R software^3^. Sequence alignment and enrichment analysis (day 0 vs PARPi-treated population) were carried out using the MAGeCK Maximum Likelihood Estimation (MLE)^41^ module and R package MAGeCKFlute Dataset^42^ of MAGeCK MLE analysis results of the CRISPR/Cas9 screen on RPE1-h*TERT* cells was extracted from the Supplemental Table 1 of Nordermeer *et al.,* (2018)^15^ Datasets were robust z-normalized and filtered against an untreated control population. Genes were considered depletion hits only when scoring at least one standard deviation under the median of each screen, under treated condition and not in an untreated control population. Functional Enrichment analysis was performed using the R package *clusterProfiler^43^*on the following databases: KEGG Pathways, Reactome, Gene Ontology: Biological Process, Complex, and Hallmark.

