## Supplementary material for "Blocking Minor Intron Splicing Disrupts DNA Repair and Overcomes Therapy Resistance in Prostate and Breast Cancer": Sup Figures

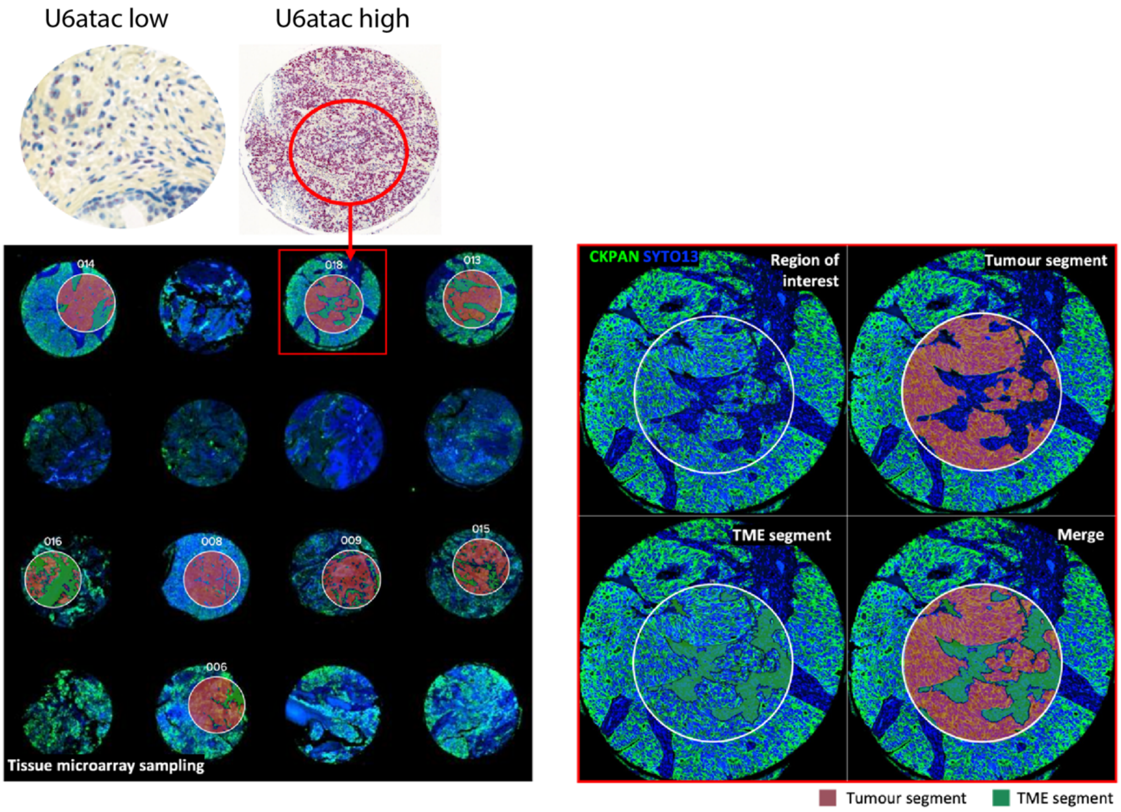


**Supplemental Figure 1. GeoMx DSP analysis of U6ATAC-high and U6ATAC-low tumour regions in a tissue microarray.** Tissue microarray (TMA) cores previously classified as U6ATAC-high or U6ATAC-low based on RNAscope labeling were sectioned onto individual slides for GeoMx Digital Spatial Profiling (DSP). Slides were stained with PanCK (Cy3) to identify epithelial/tumour regions and Syto13 (FITC) to label nuclei, enabling segmentation of tumour and tumour-microenvironment (TME) compartments. Representative TMA cores are shown (left), with tumour-enriched PanCK+ regions marked in red and PanCK– TME regions in green. For each core, regions of interest (ROIs) were selected and further segmented into tumour (PanCK+/Syto13+) or TME (PanCK–) areas of illumination (AOIs) (right).

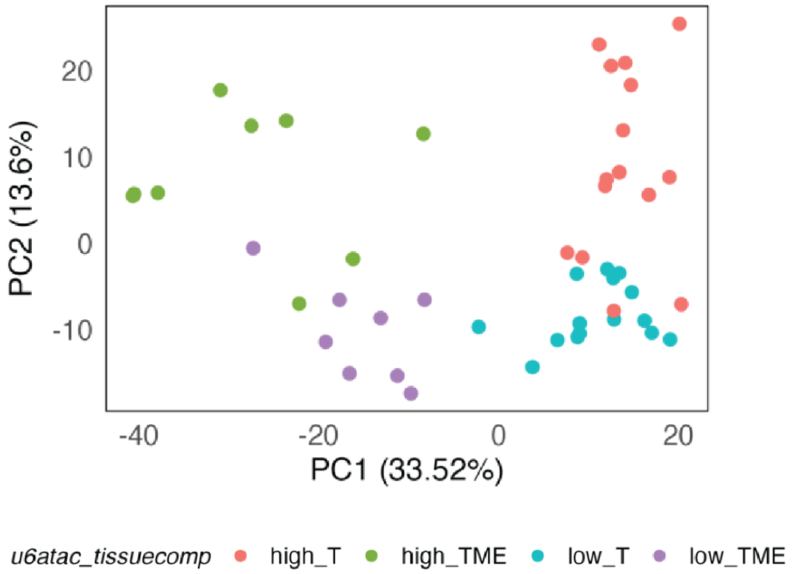


**Supplemental Figure 2. PCA of GeoMx DSP transcriptomes stratified by U6ATAC status and tissue compartment.** Principal component analysis of quality-filtered Whole Transcriptome Atlas data from tumour and tumour microenvironment (TME) area of illuminations (AOIs) (47/49 passing QC) shows clear segregation by U6ATAC level and compartment (U6ATAC-high vs. U6ATAC-low; tumour vs. TME). Gene filtering and normalization were performed as described, and DESeq2-normalized counts were used for PCA.

A

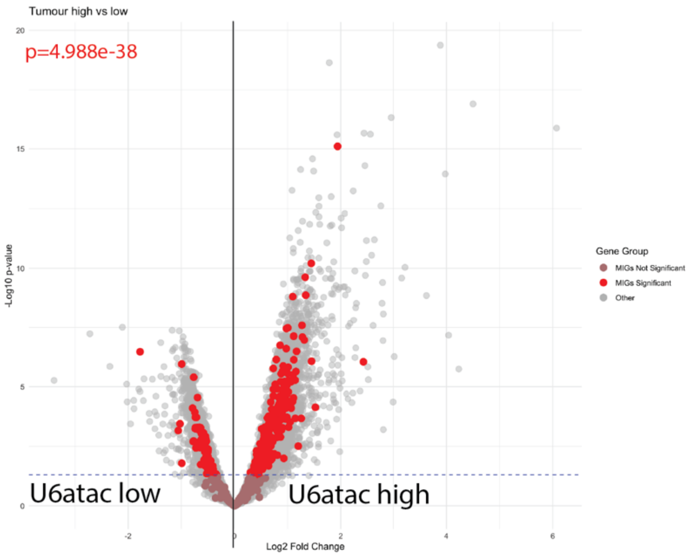
Tumour high vs low
p=4.988e-38
-Log10 p-value
Log2 Fold Change
U6atac low
U6atac high
Gene Group
MIGs Not Significant
MIGs Significant
Other

B

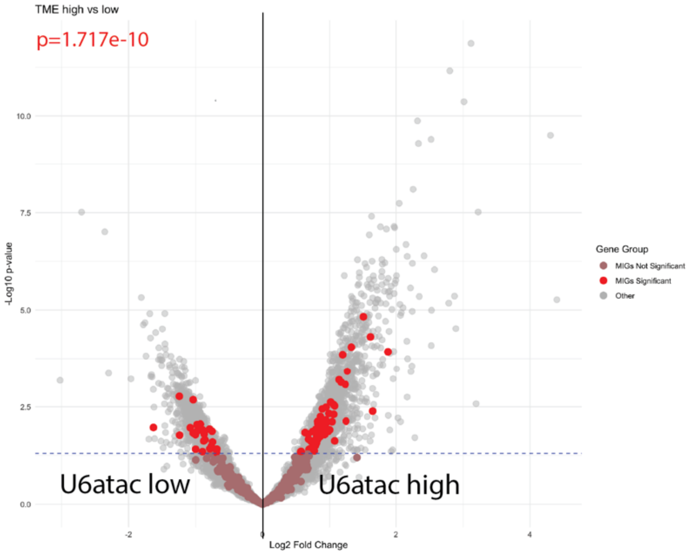
TME high vs low
p=1.717e-10
-Log10 p-value
Log2 Fold Change
U6atac low
U6atac high
Gene Group
MIGs Not Significant
MIGs Significant
Other

C

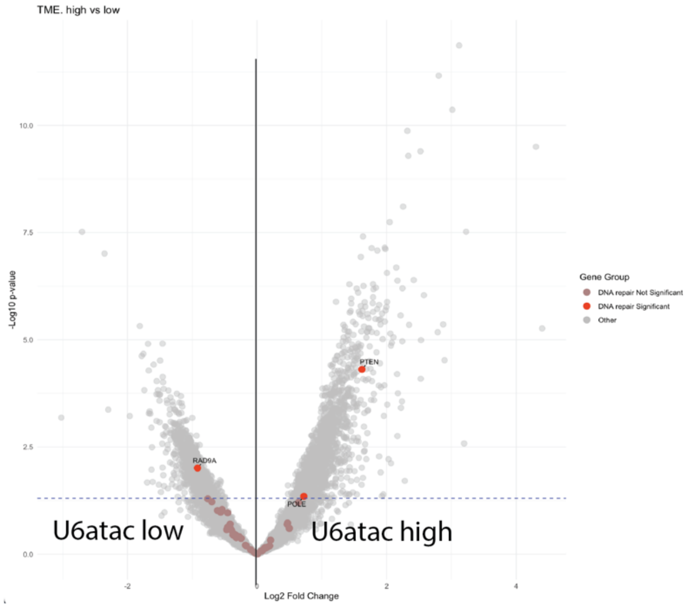
TME. high vs low
-Log10 p-value
Log2 Fold Change
PTEN
RAD9A
POLE
U6atac low
U6atac high
Gene Group
DNA repair Not Significant
DNA repair Significant
Other

**Supplemental Figure 3. Differential expression analysis of U6ATAC-high versus U6ATAC-low regions in tumour and TME compartments.** A) Volcano plot showing differential gene expression between U6ATAC-high and U6ATAC-low tumour segments. B) Volcano plot for the tumour microenvironment (TME) segments. Differential expression was performed using DESeq2 (Wald test) with Benjamini–Hochberg correction; genes with adjusted $p < 0.05$ were considered significant. Minor intron genes (MIGs) from a curated list are highlighted, with significant MIGs in red and non-significant MIGs in pale red/brown. Processed expression values were additionally compared between U6ATAC groups using the Wilcoxon rank-sum test. Volcano plots display $\log_2$ fold change versus $-\log_{10}(p$ value). C) Volcano plot depicting differential gene expression between U6atac-high and U6atac-low TMEs. DNA repair–associated genes significantly enriched in U6atac-high tumors are highlighted in red, with selected key regulators of the DNA damage response labeled

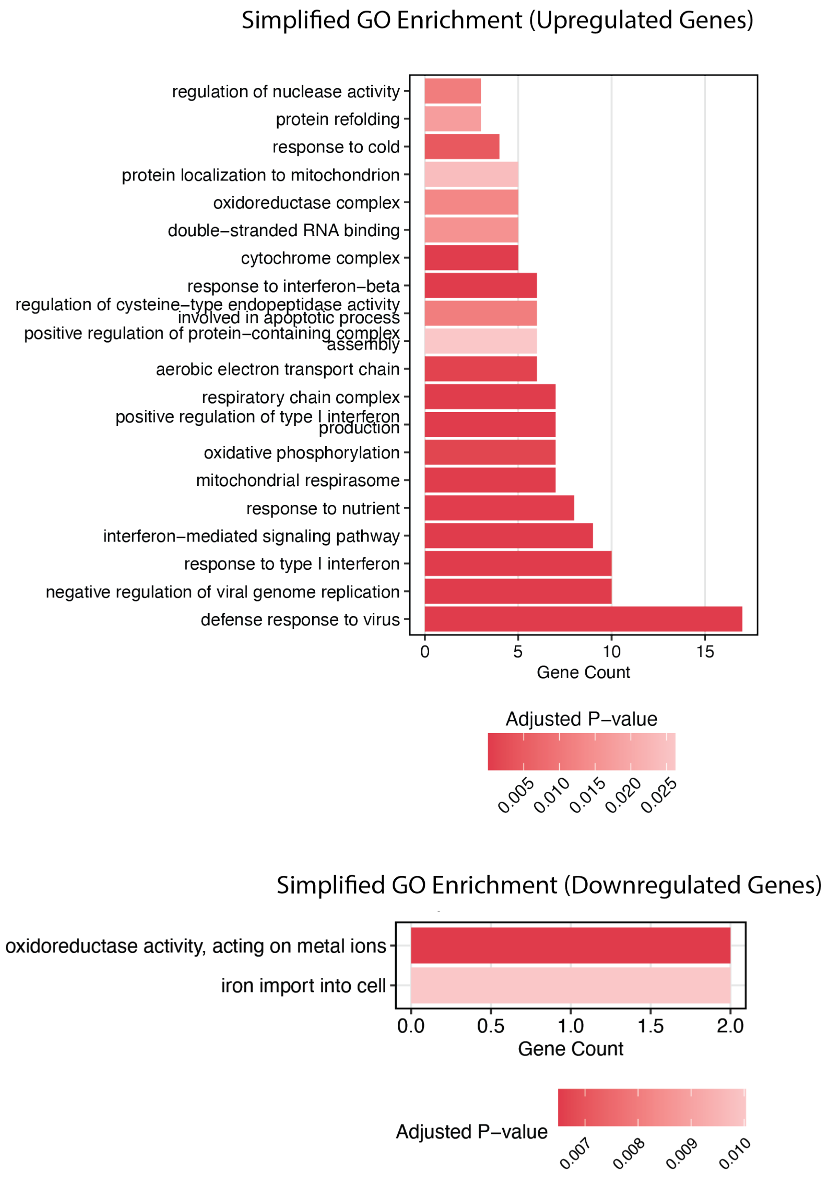


**Supplemental Figure 4 GO enrichment for genes dysregulated in U6atac-high tumors.** Spatial transcriptomics analysis showing simplified GO terms enriched among upregulated (top) and downregulated (bottom) genes in U6atac-high tumor regions. Bars indicate gene counts; color scale reflects adjusted P-values.

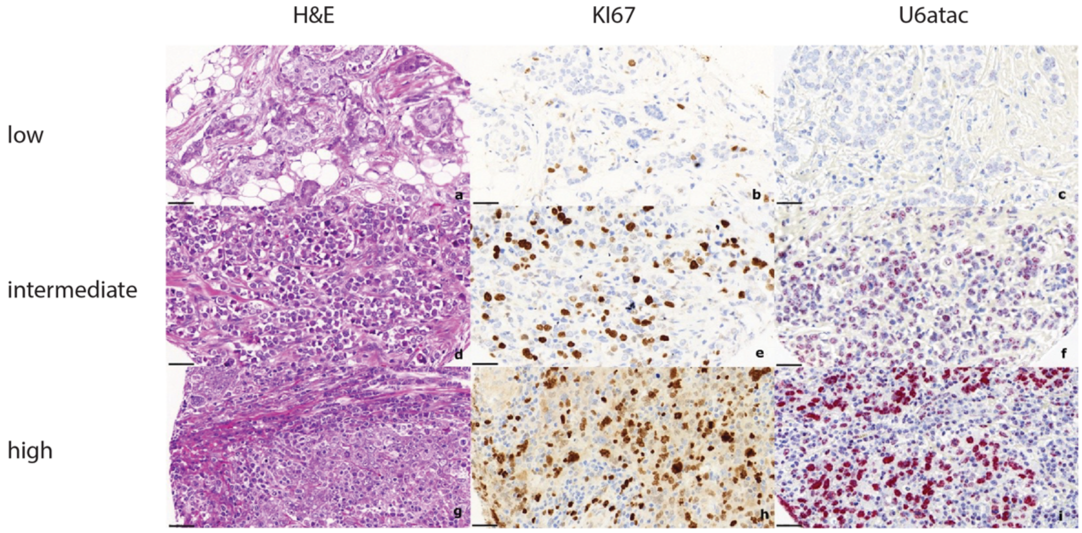


**Supplemental Figure 5.** Comparison of breast cancer (NST) with low (a-c), intermediate (d-f), and high proliferation index (g-i). Left: Corresponding HE. Middle: Ki67 immunohistochemical staining. Right: RNAish for U6atac. All in 40x magnification with a scale bar of 40uM

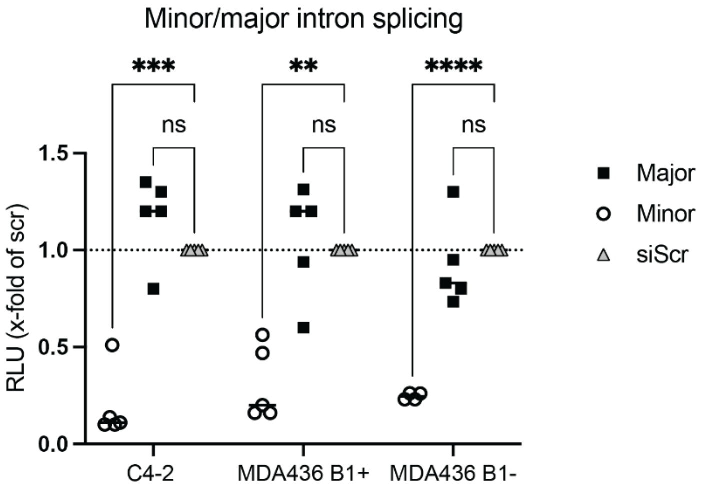


**Supplemental Figure 6. U6atac depletion selectively impairs minor intron splicing.** Dual-luciferase minor/major intron splicing reporter assays performed in C4-2 PCa cells and BRCA1-positive (B+) and BRCA1-negative (B-) MDA-MB-436 BCa cells transfected with siU6atac or siScr control. Relative luminescence units (RLU) are shown as fold-change over siScr. U6atac knockdown markedly reduced minor intron splicing in all three models (C4-2: adjusted $p = 0.0009$; MDA436 B1+: adjusted $p = 0.0022$; MDA436 B1–: adjusted $p < 0.0001$), while major intron splicing remained unchanged (ns). Data were analyzed using a mixed-effects model (REML) with Dunnett’s post hoc test.

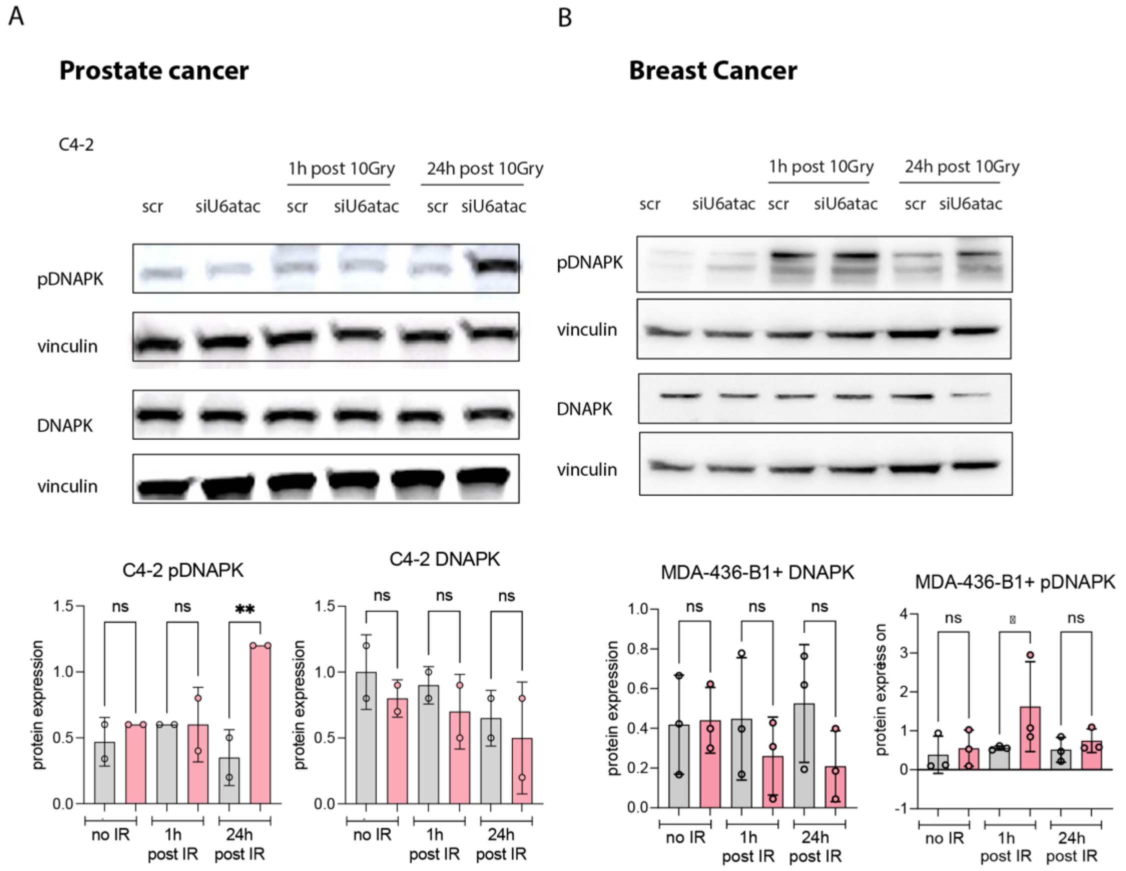


**Supplemental Figure 7.** (A) Western blot analysis of C4-2 prostate cancer cells (N=4) and (B) BRCA1-positive MDA-MB-436 cells (N=3) transfected with siU6ATAC or siScr control and exposed to 10 Gy irradiation. Cells were collected at baseline (no IR), 1 h, or 24 h post-irradiation. Blots were probed for phosphorylated DNA-PK (pDNA-PK), total DNA-PK (450 kDa), and vinculin as a loading control. Quantification of protein expression is shown below each blot. Post-hoc testing showed a significant increase in pDNA-PK at 24 h after siU6atac versus siScr ($p$=0.0211). Total DNA-PK levels were unchanged. In MDA-MB-436 B1$^{+}$ cells, siU6ATAC induced a modest, transient increase in pDNA-PK at 1 h post-irradiation ($p$<0.05).

| | LFQ: Not Selected | LFQ: Selected |
|---|---|---|
| Top3: Not Selected | 1762 | 0 |
| Top3: Selected | 5 | 540 |

Filter for proteins that were captured in at least 2 reps of the nuclear fraction for either bait

| | IgG: Not Seen | IgG: Seen |
|---|---|---|
| 59K: Not Seen | 30 | 6 |
| 59K: Seen | 380 | 124 |

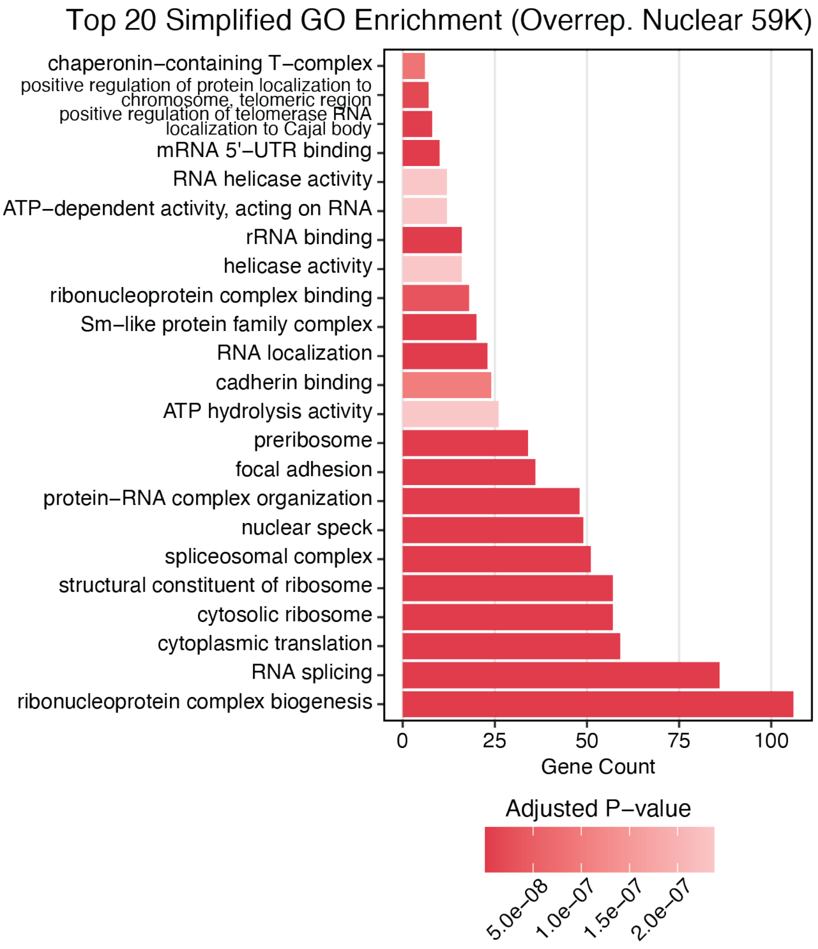


**Supplemental Figure 8 PDCD7 pulldown interactome and GO enrichment analysis.** PDCD7 nuclear pulldown mass spectrometry was analyzed using two independent quantification strategies (LFQ and iTop3). Proteins enriched by both methods and detected in ≥2 nuclear replicates were considered high-confidence interactors. Gene Ontology enrichment of the intersecting significant proteins reveals strong overrepresentation of RNA-binding, spliceosomal, and ribonucleoprotein complex biogenesis pathways. Bars indicate gene counts, with color representing adjusted P-values.

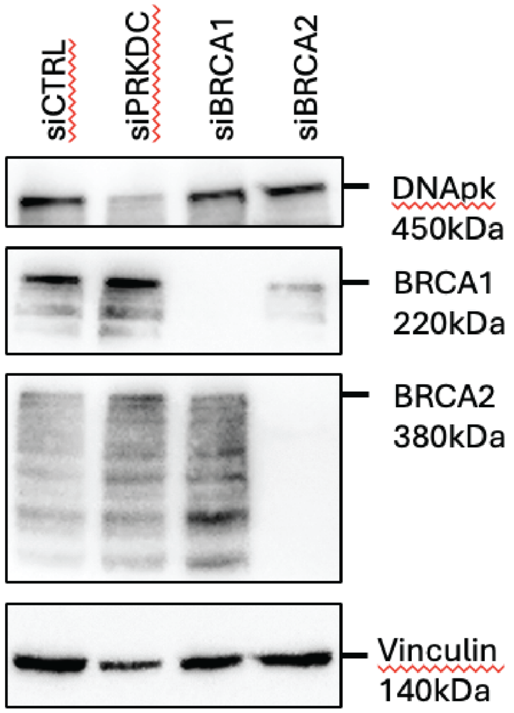


**Supplemental Figure 9. siRNA-mediated knockdown of DNA-PK, BRCA1, and BRCA2 in HEK293 cells.** HEK293 cells were transfected with control siRNA (siCTRL) or siRNAs targeting PRKDC (siPRKDC; DNA-PKcs)**,** BRCA1 (siBRCA1)**,** or BRCA2 (siBRCA2)**.** Whole-cell lysates were collected 72 h post-transfection and analysed by Western blot. Immunoblotting shows efficient depletion of DNA-PKcs (450 kDa), BRCA1 (220 kDa), and BRCA2 (380 kDa) in their respective siRNA conditions. Vinculin (140 kDa) was used as a loading control

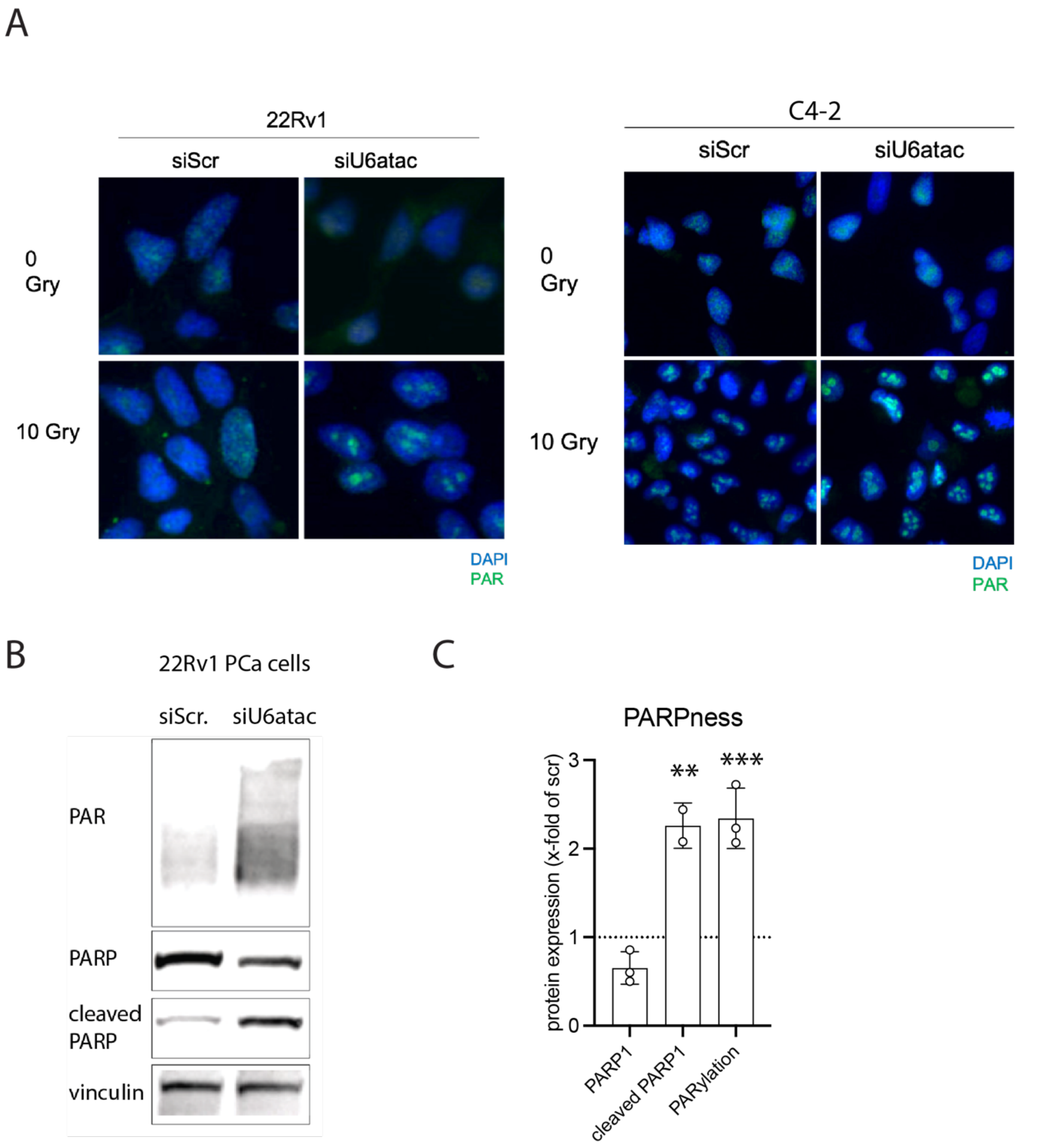


**Supplemental Figure 10. A) U6atac knockdown increases PARP1 expression and PARylation in 22Rv1 prostate cancer cells.** (A Representative immunoblot of 22Rv1 cells transfected with control siRNA (siScr) or siU6atac for 96 h. Protein levels of PAR, total PARP1, cleaved PARP1, and vinculin (loading control) are shown. **B)** Quantification of PARP1, cleaved PARP1, and global PARylation relative to siScr-treated cells (set to 1). Bars represent mean ± SD from N = 3 independent experiments. Two-way ANOVA assessed statistical significance compared with siScr; p < 0.01, *p < 0.001. **C)** Representative immunofluorescence images of PARylation in 22Rv1 and C4-2 cells 24h post-irradiation, treated with the indicated siRNAs with and without irradiation

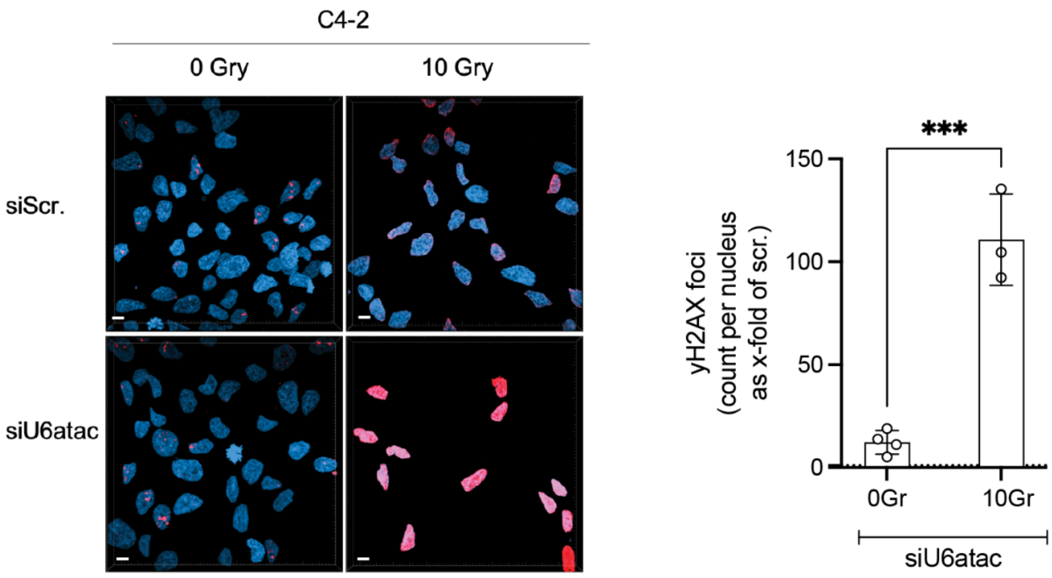


**Supplemental Figure 11. U6atac depletion increases γH2AX foci formation in C4-2 prostate cancer cells after irradiation.** Representative immunofluorescence images of γH2AX foci (magenta) in C4-2 cells treated with control siRNA (siScr) or siU6atac and collected 24 h after exposure to 0 or 10 Gy ionizing radiation. Nuclei are shown in blue. Scale bar: 10 µm. Quantification of γH2AX foci per nucleus (right) is expressed as x-fold relative to siScr 0 Gy. Data represent N = 3 independent experiments, with individual data points shown. Statistical analysis was performed using an unpaired two-tailed t-test, comparing 10 Gy vs. 0 Gy conditions: *P = 0.0003

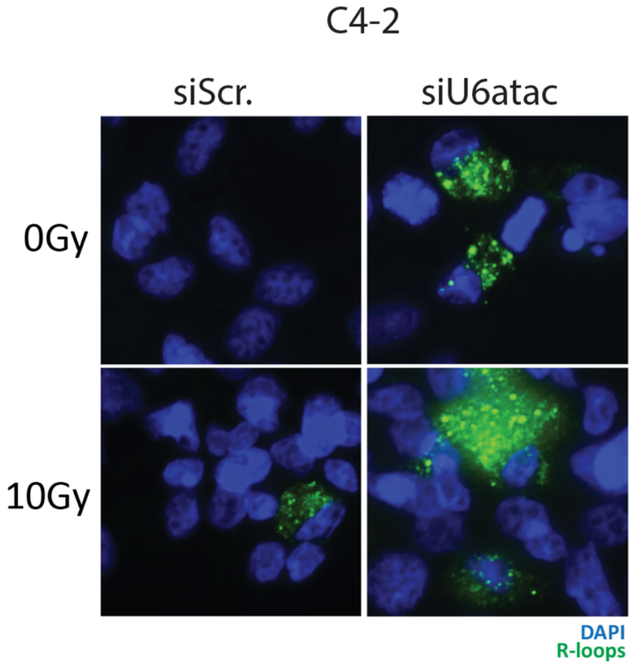


**Supplemental Figure 12.** Representative immunofluorescence images of R-loops formation 24h post-irradiation in C4-2 cells treated with the indicated siRNAs.

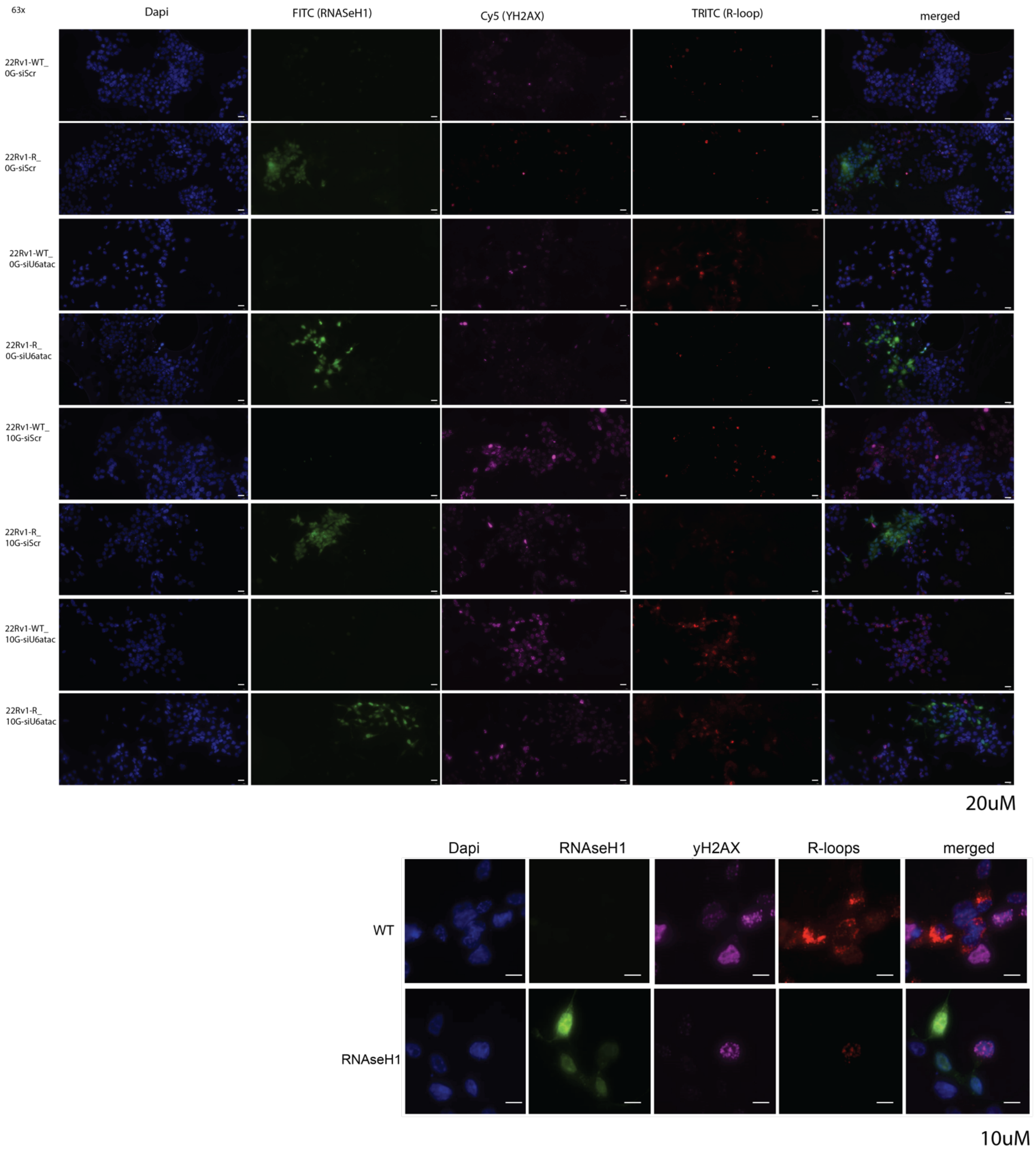


**Supplemental Figure 13 Representative immunofluorescence images of 22Rv1 cells with or without RNaseH1 overexpression.** 22Rv1 cells expressing wild-type control or RNaseH1 were stained for DAPI (nuclei), RNaseH1 (FITC), γH2AX (Cy5), and R-loops (TRITC). Images show changes in R-loop abundance and DNA damage signaling (γH2AX) upon RNaseH1 overexpression across indicated treatment conditions and doses. Scale bars as shown

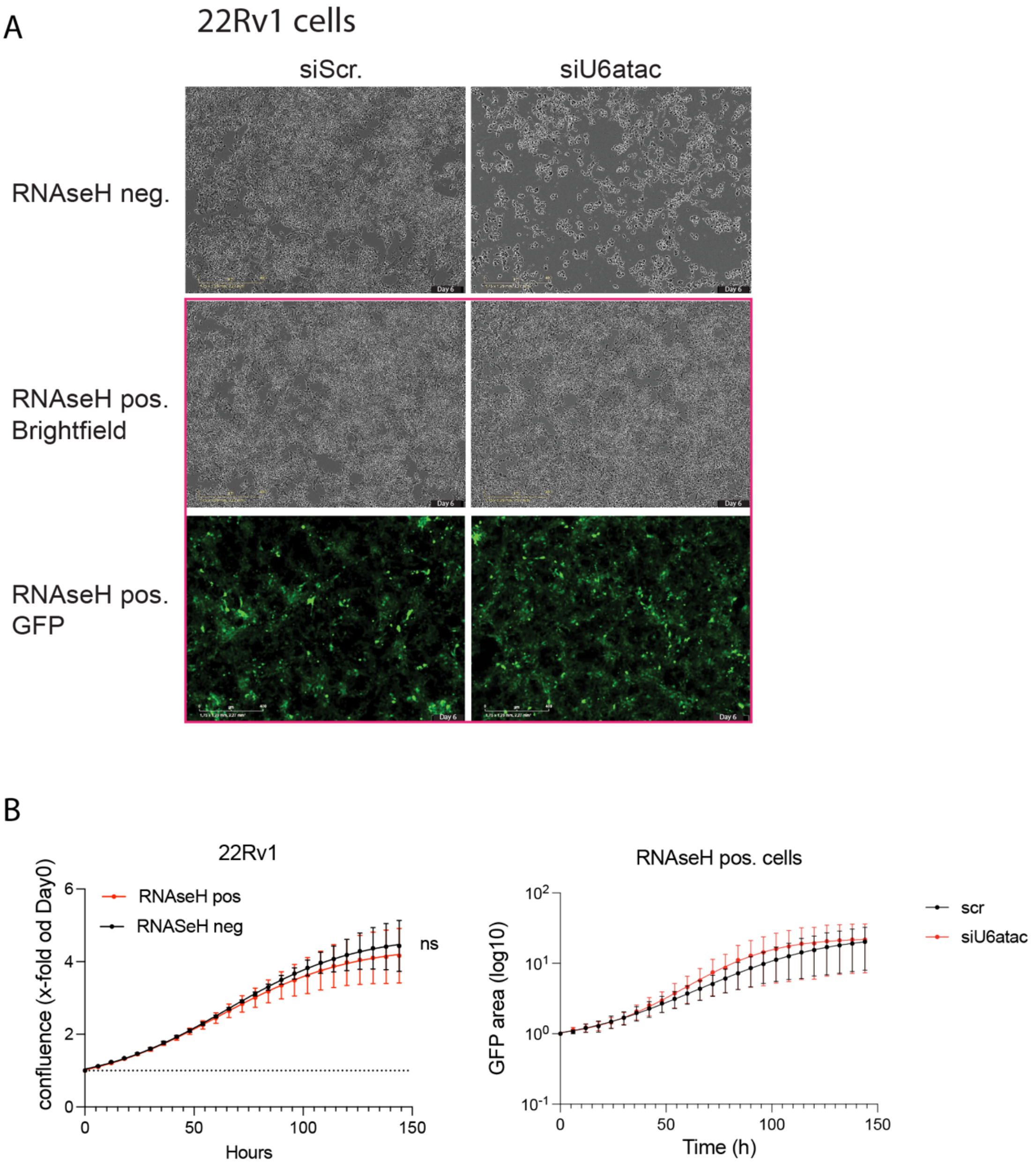


**Supplemental Figure 14. Growth of 22Rv1 WT and RNaseH1⁺ cells after U6atac depletion.** (A) Brightfield and GFP images of WT and RNaseH1-overexpressing (GFP⁺) cells treated with siScr or siU6atac. B) Growth curves of WT vs. RNaseH1⁺ cells (left) and of RNaseH1⁺ cells treated with siScr or siU6atac (right). The right panel quantifies only GFP-positive (RNaseH1⁺) cells. The right y-axis is labeled $log_{10}$, but values are plotted on a linear scale. No differences were observed (ns).

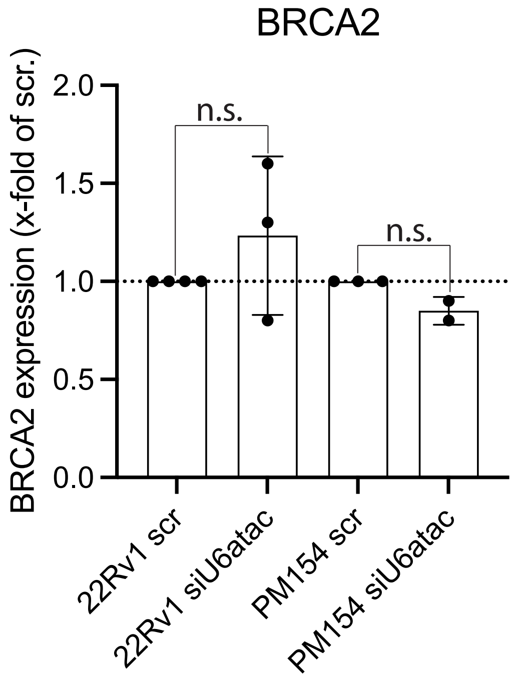


**Supplemental Figure 15 BRCA2 mRNA levels are unchanged after U6atac knockdown in 22Rv1 and PM154 cells.** Relative BRCA2 expression measured by qPCR in 22Rv1 and PM154 cells treated for 96 h with control siRNA (scr) or siU6atac. Expression values are shown as x-fold of scr. Data represent N = 3 independent experiments, with individual data points displayed. Statistical analysis was performed using a two-way ANOVA.

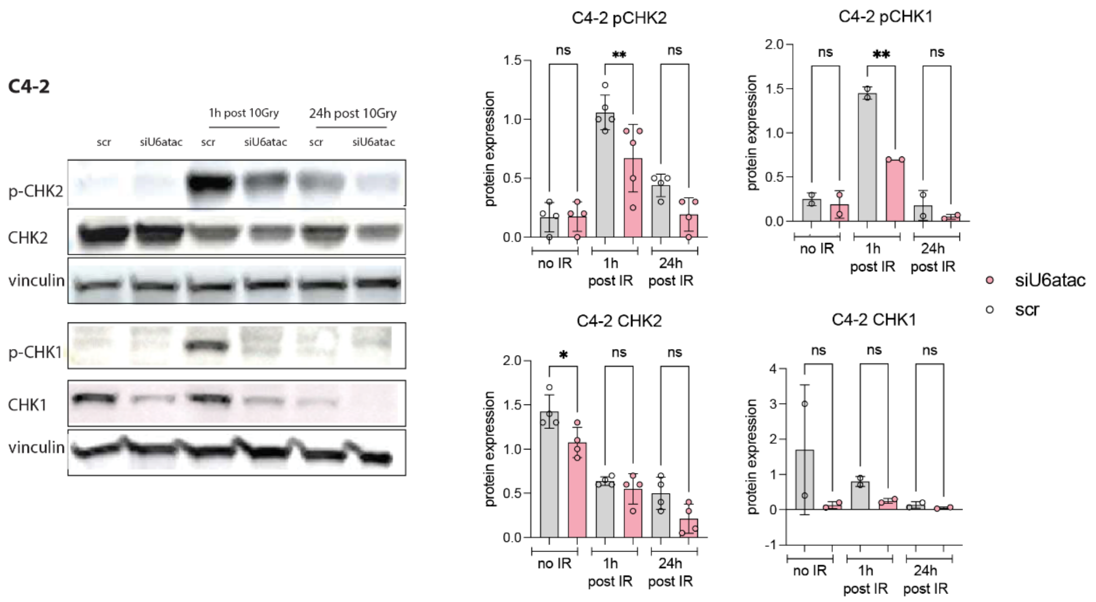


**Supplemental Figure 16 U6atac depletion alters CHK1/CHK2 signaling dynamics after irradiation in C4-2 prostate cancer cells.** Immunoblot analysis (left) and quantification (right) of phosphorylated CHK2 (p-CHK2), total CHK2, phosphorylated CHK1 (p-CHK1), and total CHK1 in C4-2 cells treated with control siRNA (scr) or siU6atac. Cells were irradiated with 10 Gy and harvested at 1 h or 24 h post-irradiation or left untreated (no IR). Vinculin was used as a loading control. Bar graphs show normalized protein expression levels relative to scr controls for each time point. Data represent N = 4 biological replicates; individual values are displayed. Statistical analysis was performed using ordinary one-way ANOVA.

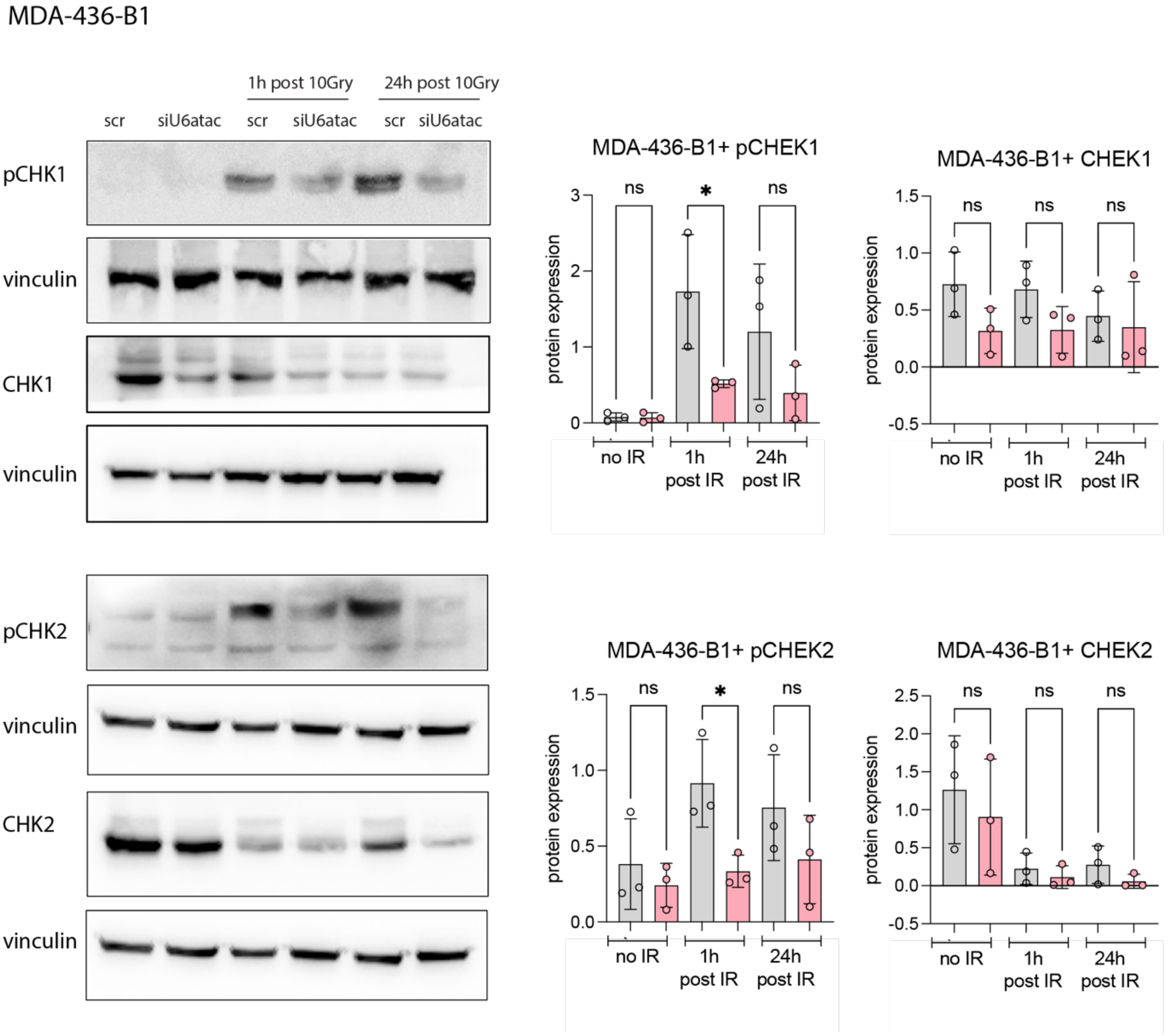


**Supplemental Figure 17 U6atac depletion alters CHK1/CHK2 signaling dynamics after irradiation in MDA-436-B1+ breast cancer cells.**
Immunoblot analysis (left) and quantification (right) of phosphorylated CHK1 (p-CHK1), total CHK1, phosphorylated CHK2 (p-CHK2), and total CHK2 in MDA-436-B1+ cells treated with control siRNA (scr) or siU6atac. Cells were irradiated with 10 Gy and harvested at 1 h or 24 h post-irradiation or left untreated (no IR). Vinculin was used as a loading control. Bar graphs show normalized protein expression levels relative to scr controls for each time point. Data represent N = 4 biological replicates; individual values are displayed. Statistical analysis was performed using ordinary one-way ANOVA.

Prostate Cancer

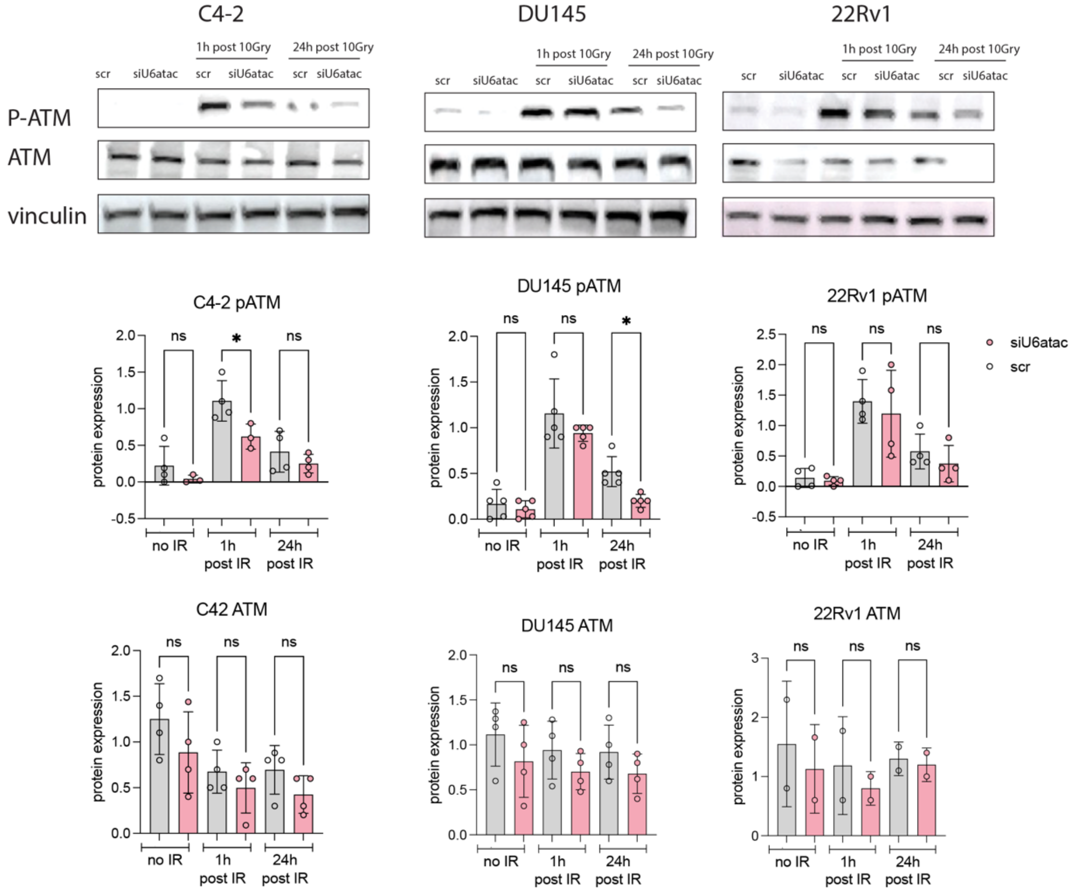


Breast Cancer

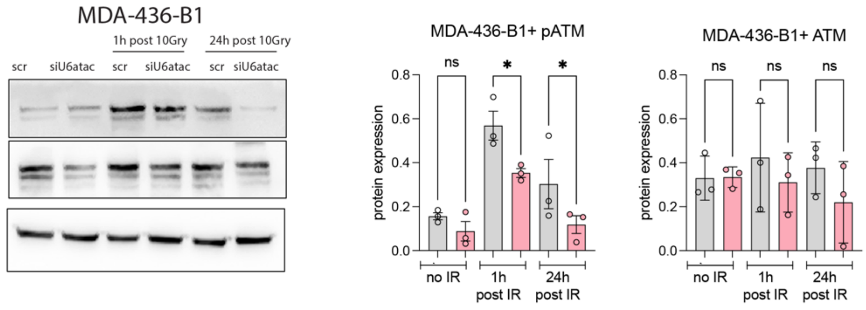


**Supplemental Figure 18. ATM activation following irradiation in prostate and breast cancer cell lines after U6atac depletion.** Immunoblot analysis of phosphorylated ATM (p-ATM) and total ATM in three PCa cell lines (C4-2, DU145, 22Rv1) and one BCa cell line (MDA-436-B1+) treated with control siRNA (scr) or siU6atac. Cells were exposed to 10 Gy ionizing radiation and harvested at 1 h or 24 h post-irradiation or left untreated (no IR). Vinculin served as a loading control. Quantification panels show normalized protein expression relative to scr controls for each cell line and time point. Data for PCa cell lines represent N = 4 biological replicates; BCa data represent N = 3 biological replicates. Statistical analysis was performed using ordinary one-way ANOVA for each condition. Significant differences are indicated as: *$P < 0.05$

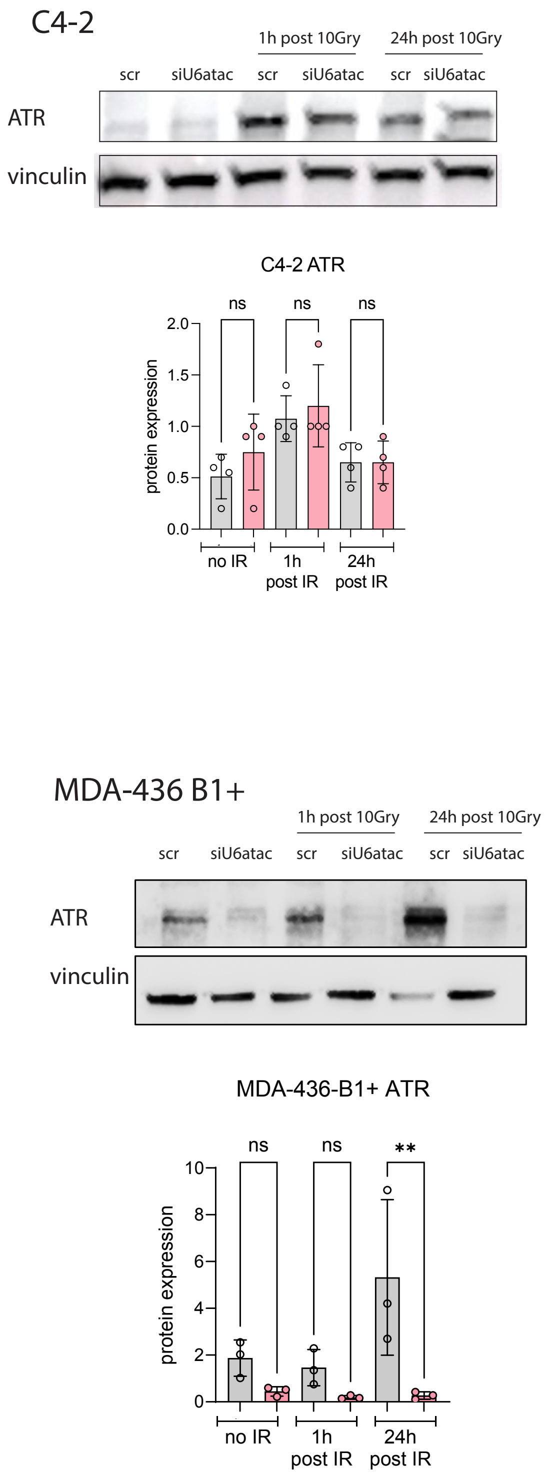


**Supplemental Figure 19. ATR levels after U6atac depletion in prostate and breast cancer cells.** Immunoblot analysis of ATR in C4-2 prostate cancer cells (N = 4) and MDA-436-B1+ breast cancer cells (N = 3) treated with scr or siU6atac and collected at 0, 1 h, and 24 h after 10 Gy irradiation. Vinculin was used as a loading control. Quantification shows no change in ATR in C4-2 cells, whereas ATR is significantly reduced in MDA-436-B1 cells at 24 h post-IR. Statistics: ordinary one-way ANOVA; ns, not significant; ** $P < 0.01$

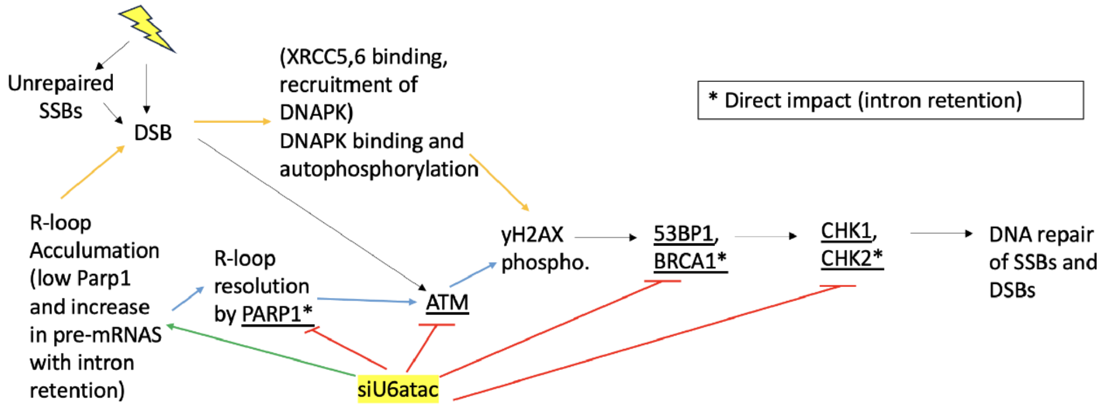


**Supplemental Figure 20. Model summarizing how U6atac depletion disrupts DNA damage responses.** Loss of U6atac impairs minor intron splicing in key DNA repair genes (indicated with *), reducing PARP1 levels, ATM/CHK1/CHK2/BRCA1 signaling, and R-loop–resolving capacity. This leads to the accumulation of unrepaired SSBs and R-loops, which convert into DSBs during replication. DSBs trigger DNA-PK binding and autophosphorylation, γH2AX formation, and 53BP1/BRCA1 recruitment, but these pathways are compromised under U6atac knockdown. Together, these defects impair repair of both SSBs and DSBs, promoting persistent DNA damage and replication stress.

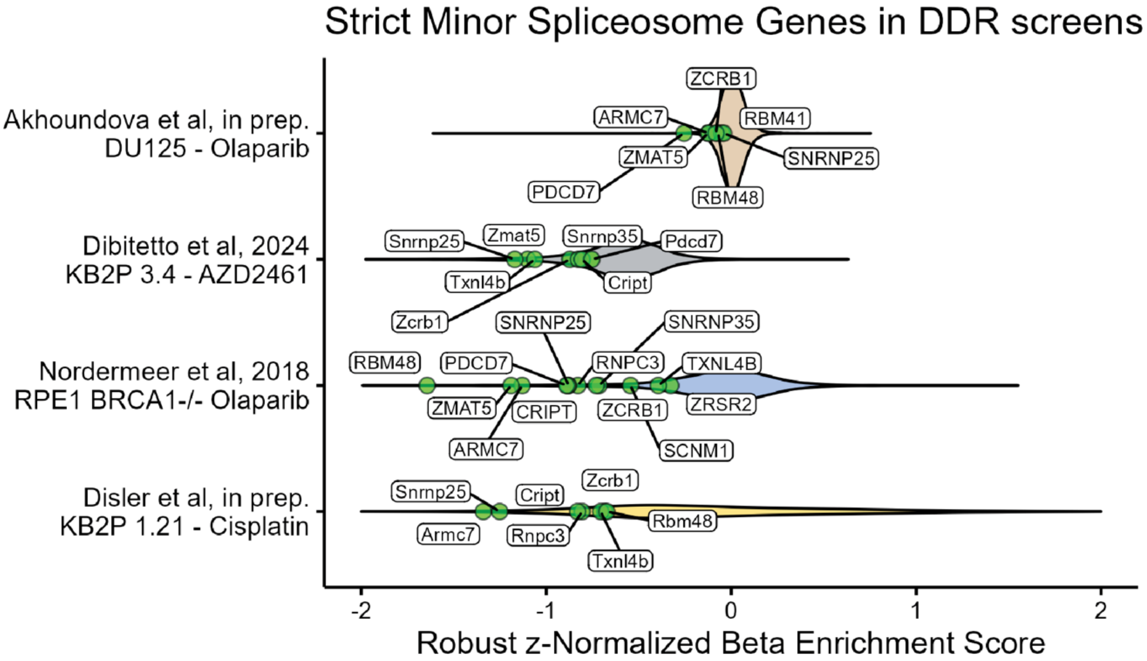


**Supplemental Figure 21 Minor spliceosome genes highlighted across DNA damage response (DDR) CRISPR/Cas9 screens.** Violin plots show robust z-normalized β-enrichment scores for strict minor spliceosome genes from 3 published BCa and our own PCa genome-wide CRISPR/Cas9 dropout screens under Olaparib or Cisplatin conditions. Genes were called depletion hits when ≥1 SD below the median in the treated but not in the untreated condition.

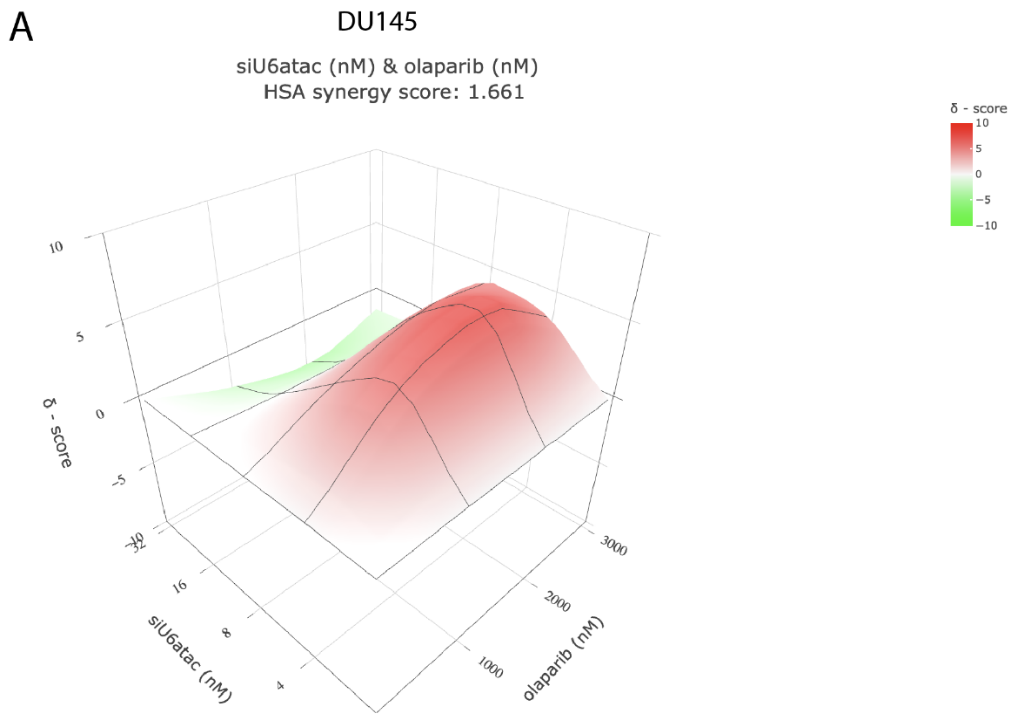


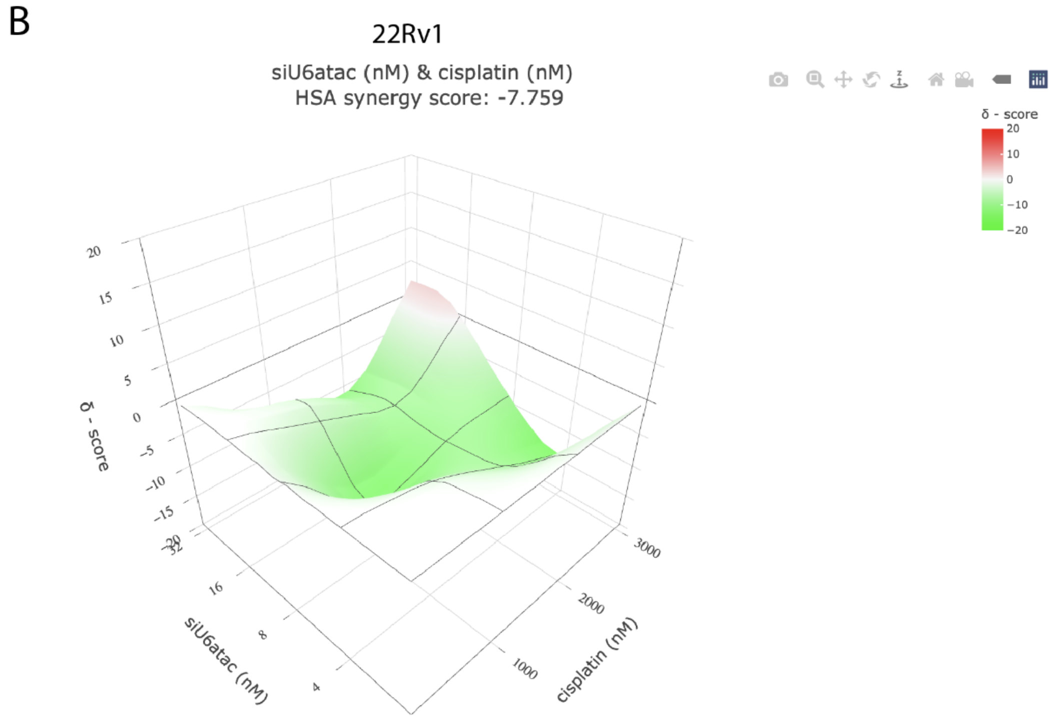


**Supplemental Figure 22. A)** 3D interaction landscapes of siU6atac (0, 4, 8, 16, 32 nM) and Olaparib (0, 0.5, 1, 2 µM) in DU145 cells, analyzed using the HSA model on SynergyFinder. Synergy scores are represented as red (>0) and green (<0). B) 3D interaction landscapes of siU6atac (0, 4, 8, 16, 32 nM) and cisplatin (0, 1, 2,3 µM) in 22Rv1 cells, analyzed using the HSA model on SynergyFinder. Synergy scores are represented as red (>0) and green (<0).

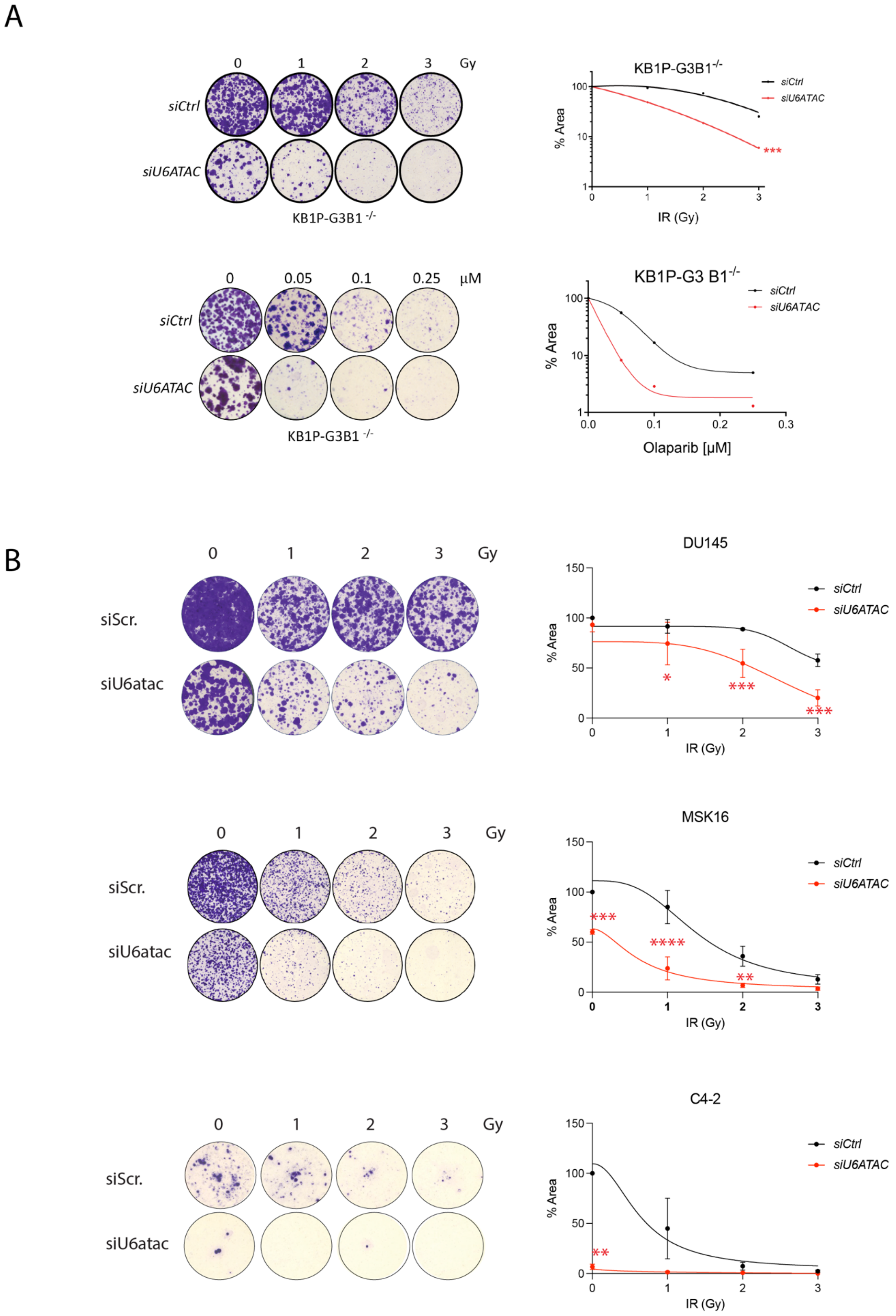


**Supplemental Figure 23 siU6atac sensitizes cancer cells to DNA-damaging agents. A)** Left panel: representative images of growth assays in KB1P-BRCA1- cells treated with the indicated siRNAs and exposed to irradiation, or olaparib at the specified doses. Right panel: statistical analysis was performed using 2-way ANOVA followed by Dunnett's test. B) Left panel: representative images of growth assays in DU145, MSK16, and C4-2 cells treated with the indicated siRNAs and exposed to irradiation at the specified doses. Right panel: statistical analysis was performed using 2-way ANOVA followed by Dunnett's test

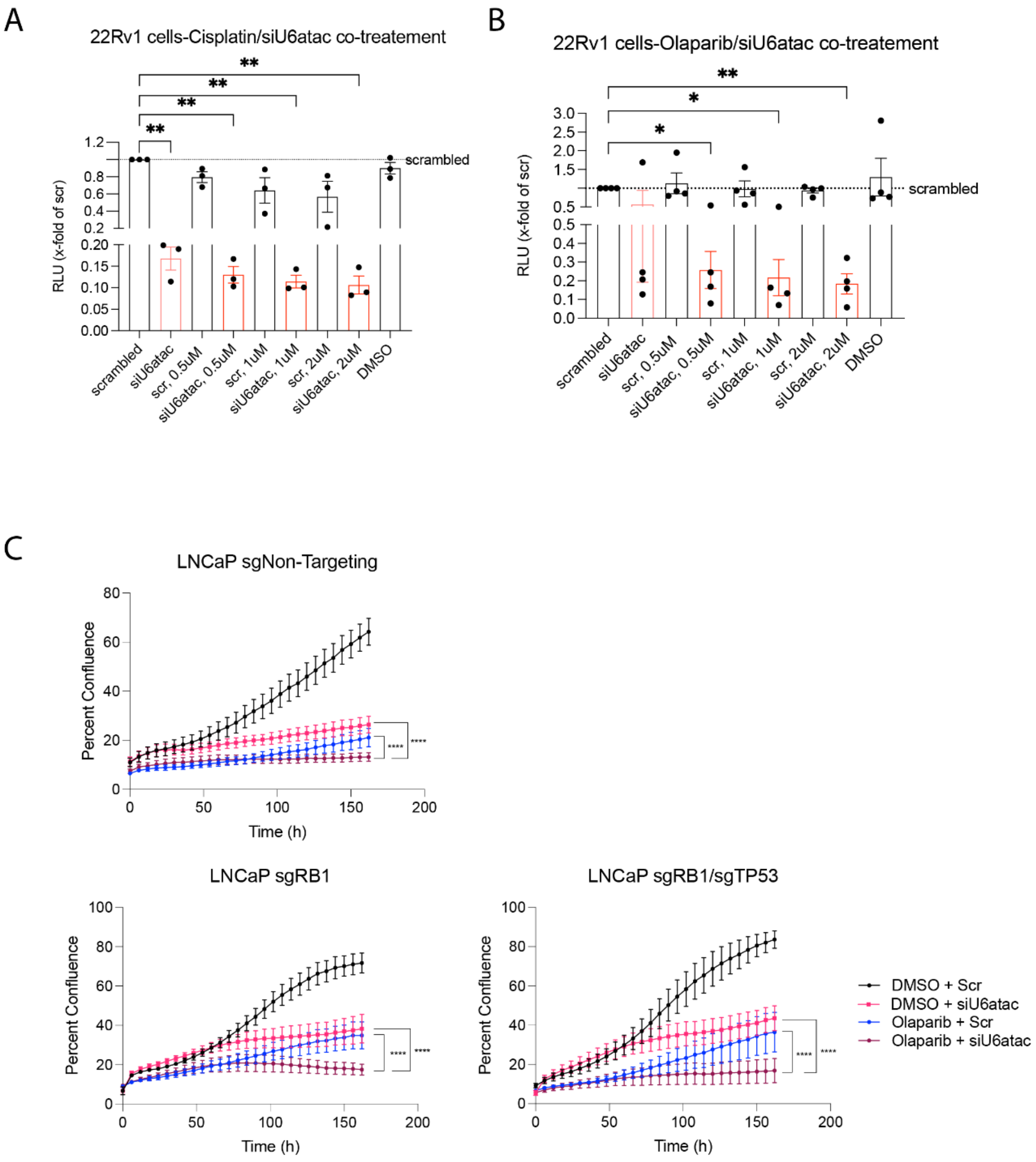


**Supplemental Figure 24 A)** Cell viability of 22Rv1 cells treated with siU6atac (2nM) in combination with Cisplatin (0.5, 1, 2 µM) compared to scrambled control (dotted line). Data are shown as x-fold of scrambled control. N=3 biological replicates, each performed in triplicate. Statistical analysis: repeated measures ANOVA with Holm-Šídák's multiple comparisons. B) Cell viability of 22Rv1 cells treated with siU6atac (2nM) in combination with Olaparib (0.5, 1, 2 µM) compared to scrambled control (dotted line). Data are shown as x-fold of the scrambled control. N=4 biological replicates, each performed in triplicate. Statistical analysis: repeated measures ANOVA with Holm-Šídák's multiple comparisons **C)** Effect of U6atac depletion and PARP inhibition on the growth of LNCaP isogenic cell lines. Growth curves of parental LNCaP, LNCaP RB1–/–, and LNCaP RB1–/–/TP53–/– cells treated with DMSO or 1 µM Olaparib and transfected with siScr or siU6atac.

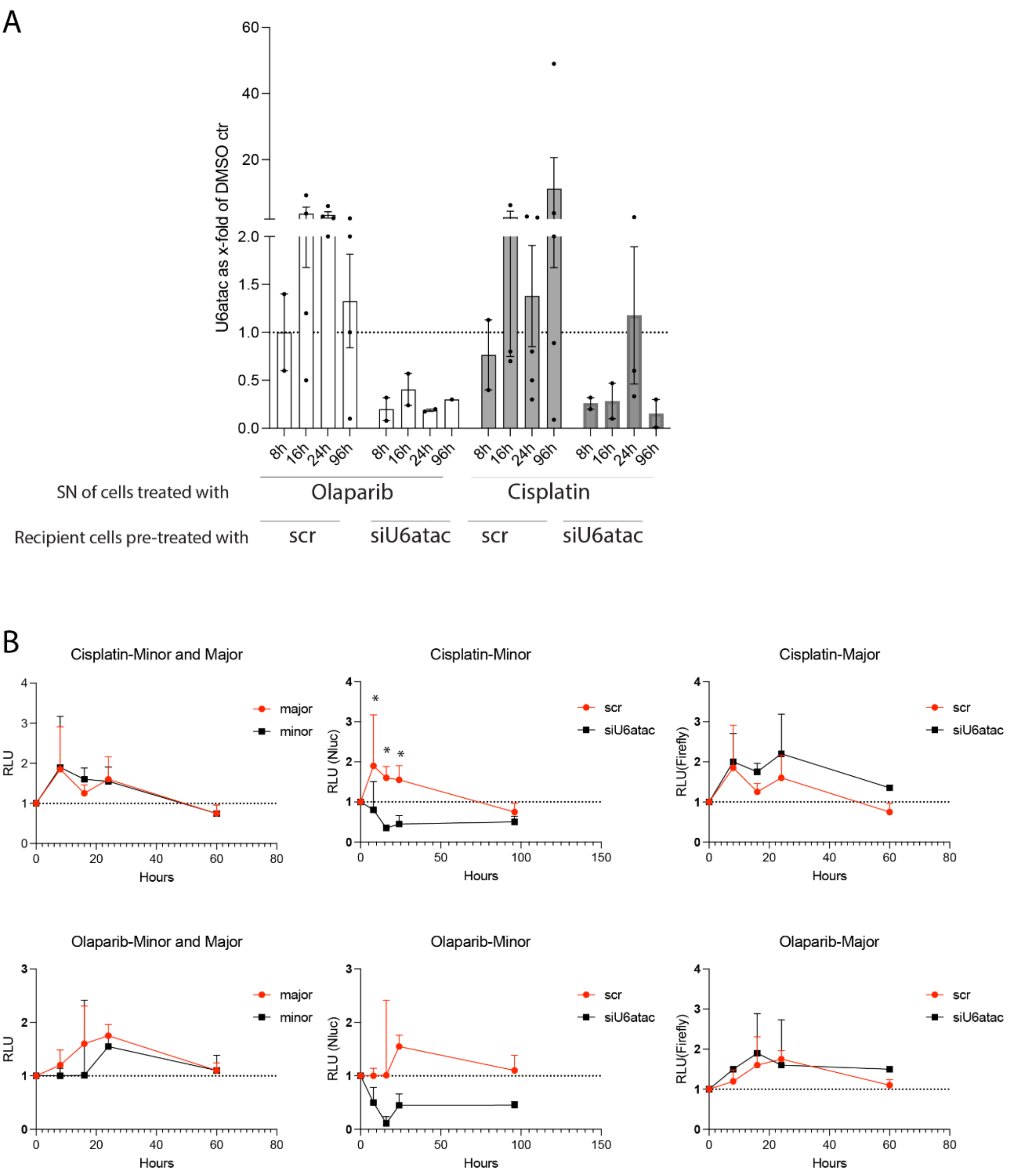


**Supplemental Figure 25. U6atac expression in recipient cells exposed to supernatants from DNA-damaged C4-2 cells.** A) qPCR analysis of U6atac levels in recipient cells pre-treated with siScr **or** siU6atac, then exposed (24 h later) to supernatants (SN) collected from C4-2 cells treated with **olaparib** or cisplatin for the indicated time points. Expression is shown as x-fold relative to DMSO controls, defined as recipient cells receiving supernatants from C4-2 cells treated with DMSO for the same time points**.** Data represent N = 2 biological replicates. B) C4-2 reporter cells pre-treated with siScr or siU6atac were incubated with supernatants from cisplatin- or olaparib-treated donor cells; timepoints indicate duration of supernatant exposure. Normalized luminescence of the minor (Nluc) and major (Firefly) reporters was measured (N = 3). siU6atac reduced minor spliceosome activity at early timepoints, while major spliceosome activity showed no significant differences.

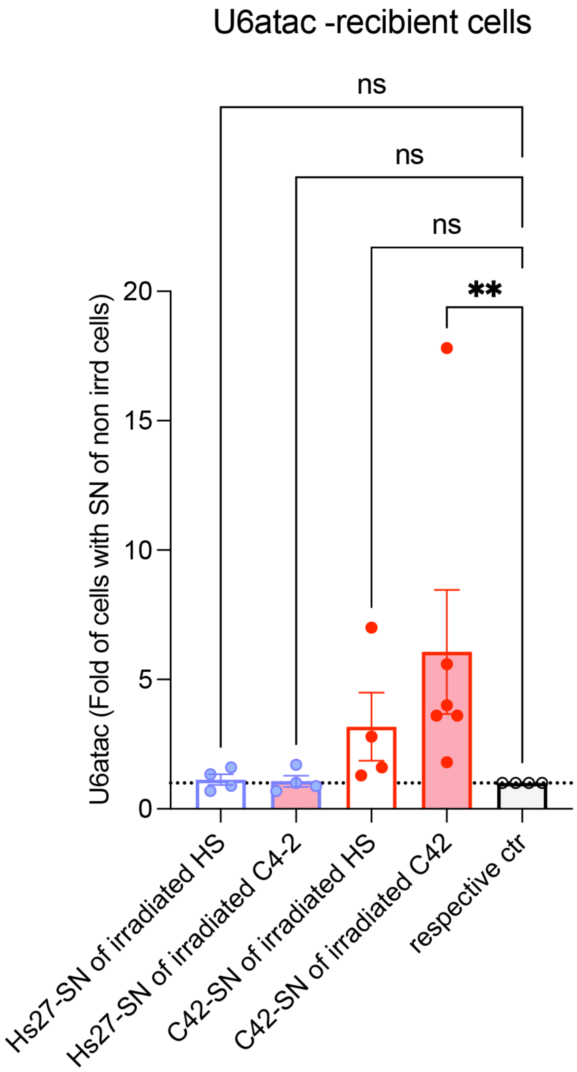


**Supplemental Figure 26. U6atac expression in Hs27 and C4-2 recipient cells after exposure to supernatants from irradiated donor cells.** qPCR of U6atac in Hs27 (N = 4) and C4-2 (N = 6) recipient cells treated with supernatants (SN) from irradiated or non-irradiated Hs27 or C4-2 donors. Expression is shown relative to each respective non-irradiated control. Dunn's post hoc: only C4-2 recipients exposed to irradiated C4-2 supernatant were significantly increased (P = 0.0058)

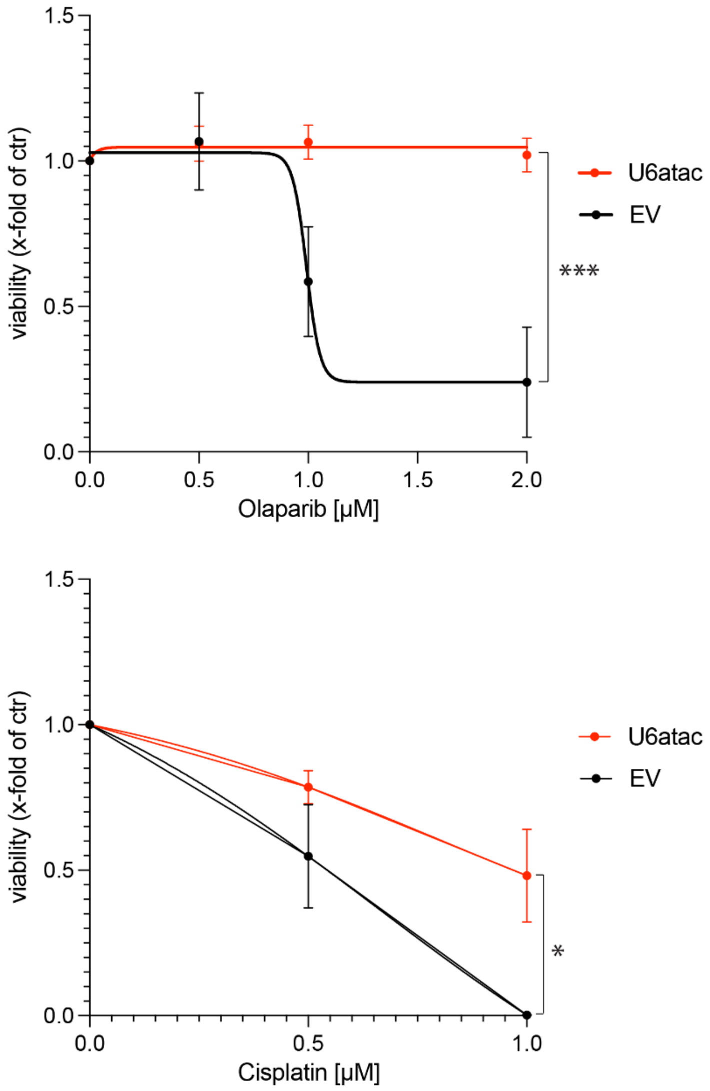


**Supplemental Figure 27. Drug sensitivity of C4-2 cells overexpressing U6atac.** Cell viability (x-fold of DMSO control) of C4-2 cells overexpressing U6atac or empty vector (EV) after treatment with the indicated concentrations of Olaparib (top, N = 4) or cisplatin (bottom, N = 3)—ordinary two-way ANOVA with Šídák's multiple comparisons test. U6atac overexpression significantly reduced sensitivity to olaparib (***P < 0.001) and mildly increased resistance to cisplatin (*P < 0.05).

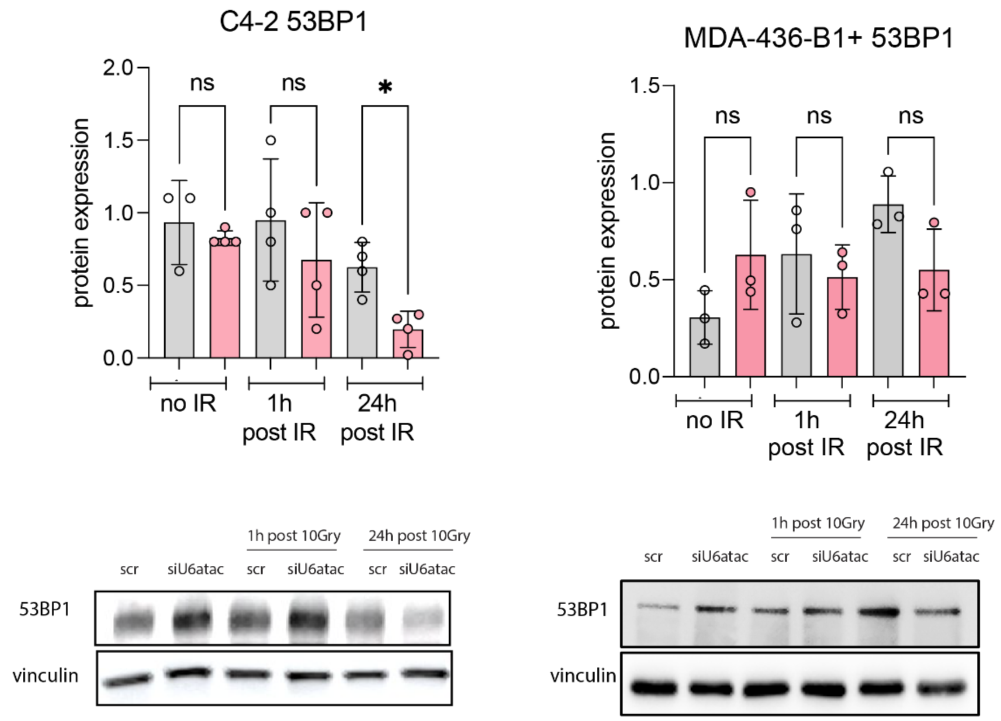


**Supplemental Figure 28. TP53BP1 protein levels after U6atac depletion and irradiation.** Immunoblot and quantification of 53BP1 in C4-2 PCa cells (N = 4) and MDA-436-B1 BCa cells (N = 3) treated with siScr or siU6atac, either untreated (no IR) or collected 1 h and 24 h after 10 Gy irradiation. Vinculin served as a loading control. Ordinary one-way ANOVA with uncorrected Fisher's LSD. (*P < 0.05; ns, not significant).

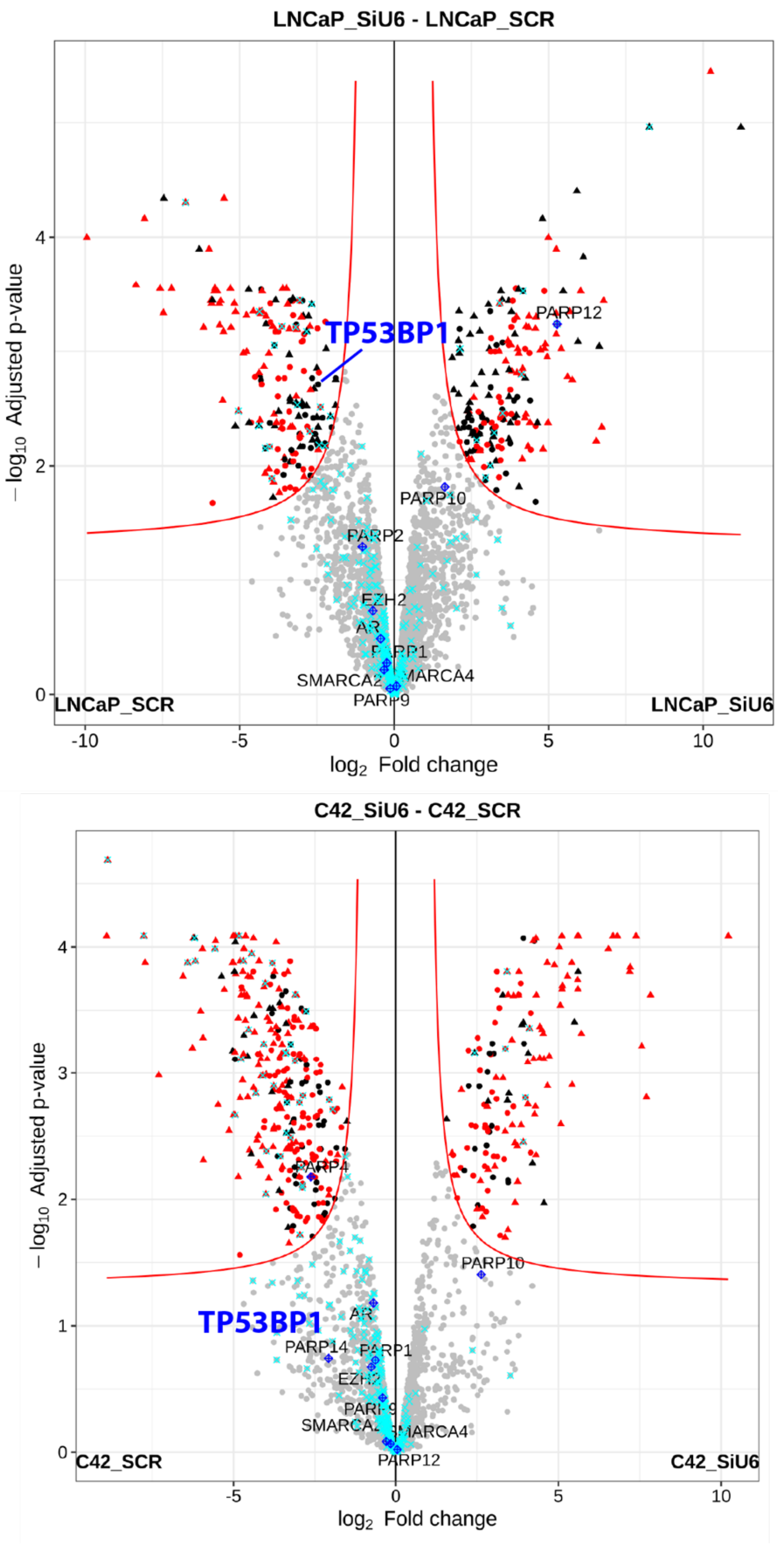


**Supplemental Figure 29. Proteomic changes after U6atac knockdown in LNCaP and C4-2 cells.** Volcano plots showing proteins up- or downregulated in LNCaP (top) and C4-2 (bottom) cells after 96 h siU6atac versus siScr. The proteomic datasets were generated in a previous publication and re-visualized here. TP53BP1 is significantly reduced in LNCaP cells, while in C4-2 cells it shows a downward trend that does not reach statistical significance.

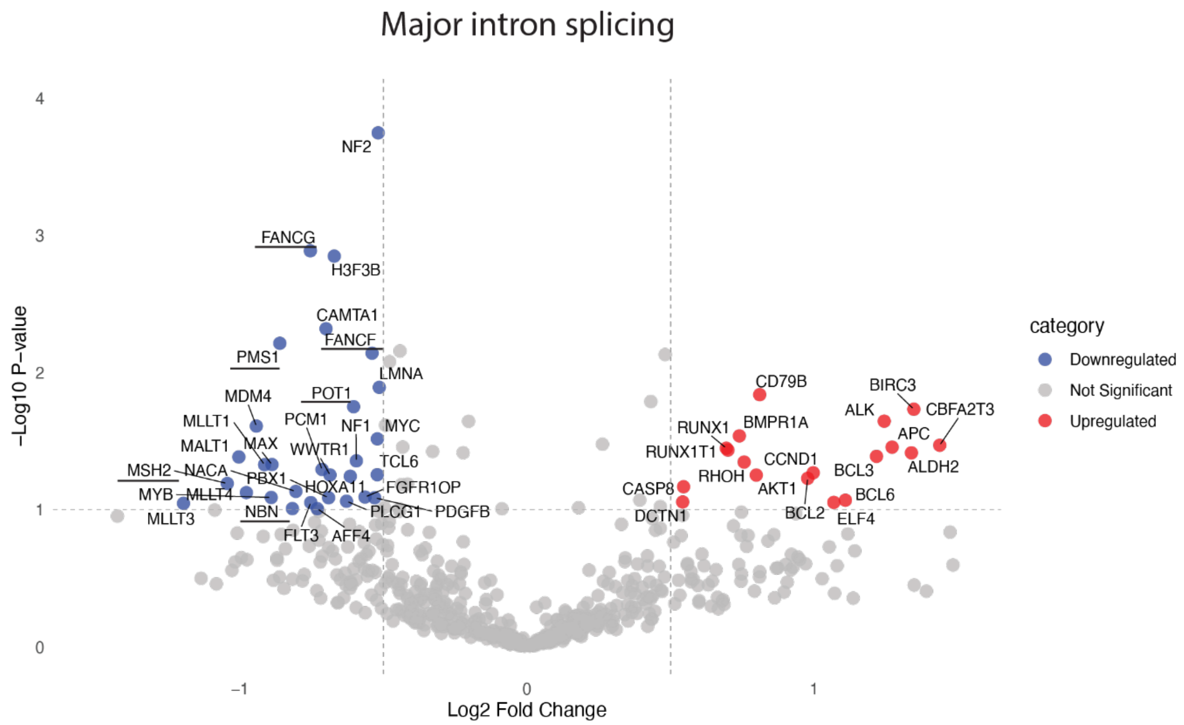


**Supplemental Figure 30.** Volcano plot comparing RNAi screen results in irradiated (10 Gy) versus non-irradiated cells. The x-axis shows the $\log_2$ fold-change in major spliceosome activity upon knockdown, and the y-axis the $-\log_{10}$ *p*-value. KDs of the genes shown on the right (in red) had a stronger effect in untreated cells, where irradiation partially rescued the major-intron splicing defect. Genes on the left (blue) represent knockdowns with a stronger effect under irradiation, where irradiation amplified the major intron splicing defect. Selected genes of interest are highlighted.

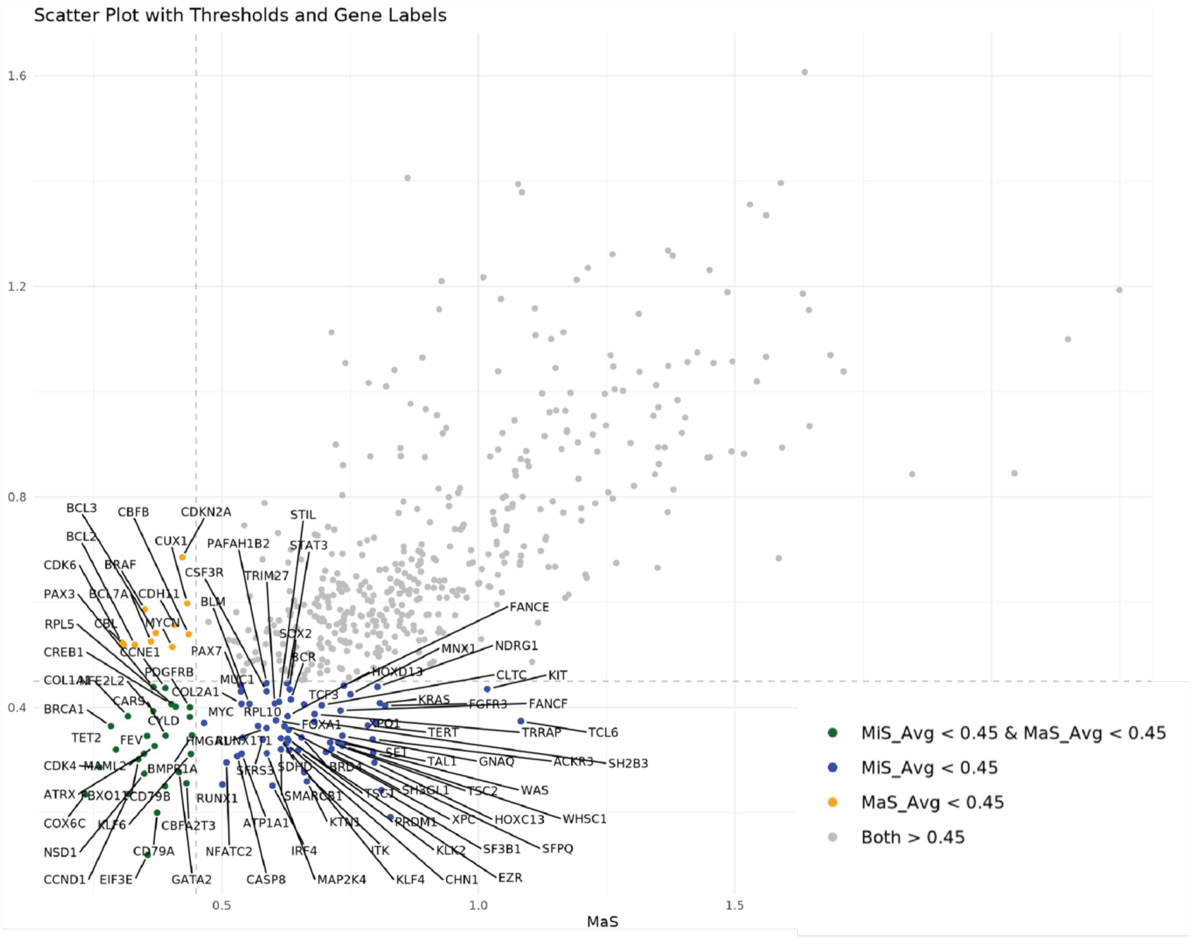


Genes of which KD decreases MiS, MaS or both

[1] "ATP1A1" "BCR" "PRDM1" "BLM" "CASP8" "RUNX1" "RUNX1T1" "CHN1" "CLTC" "COL2A1" "CSF3R" [12] "FANCE" "FANCF" "FGFR3" "GNAQ" "MNX1" "HMGA1" "FOXA1" "HOXC13" "HOXD13" "IRF4" "ITK" [23] "KIT" "KLK2" "KRAS" "KTN1" "MUC1" "MYC" "NFATC2" "PAFAH1B2" "PAX7" "TRIM27" "RPL10" [34] "SDHD" "MAP2K4" "SET" "SFPQ" "SFRS3" "SH3GL1" "STIL" "SMARCB1" "SOX2" "STAT3" "TAL1" [45] "TCF3" "TERT" "TSC1" "TSC2" "EZR" "WAS" "WHSC1" "XPC" "XPO1" "TRRAP" "KLF4" [56] "SH2B3" "NDRG1" "SF3B1" "BRD4" "TCL6" "ACKR3"

Genes of which KD decreases MiS and MaS

"ATRX" "CCND1" "BMPR1A" "BRCA1" "CARS" "CBFA2T3" "CCNE1" "CD79A" "CD79B" "CDK4" "COL1A1" "KLF6" [13] "COX6C" "CREB1" "CYLD" "GATA2" "EIF3E" "NFE2L2" "PAX3" "PDGFRB" "RPL5" "FEV" "TET2" "NSD1" [25] "FBXO11" "MAML2"

Genes of which KD decreases MaS

"BCL2" "BCL3" "BCL7A" "BRAF" "CBFB" "CBL" "CDH11" "CDK6" "CDKN2A" "CUX1" "MYCN"

Genes where KD and irradiation rescues or amplifies MiS activity

"BCL7A", "FOXL2", "BRCA1", "BTG1", "CBLB", "CDH1", "CDK4", "COX6C", "DDX10", "EXT1", "FGFR2", "HOXA11", "IL6ST", "ITK", "KIT", "KLK2", "KTN1", "MPL", "MSH2", "MYH9", "SSX2", "TAL1", "TSHR", "UTX", "TRRAP", "45541", "IL21R", "NIN", "FAM22B", "SSX4"

**Supplemental Figure 31. RNAis screen,** listed are genes that decrease MiS activity by more than 65% (blue), major spliceosome activity (MaS) by more than 65% (yellow) or both (green). Genes where KD rescued MiS activity after irradiation are listed in grey

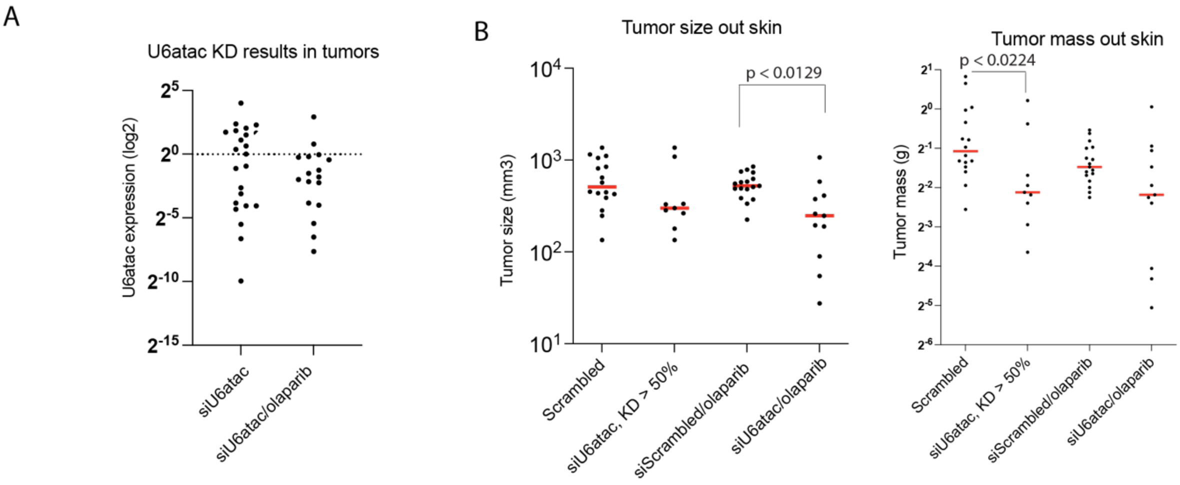


**Supplemental Figure 32. Tumor mass and size measured ex vivo across individual in vivo experiments.** A) qPCR analysis of U6atac expression in tumors harvested from mice treated with siU6atac or siU6atac + olaparib. Each dot represents one tumor. Expression values are normalized to pooled siScrambled control tumors and displayed on a $\log_2$ scale. B) Combined analysis of tumor size ($mm^3$) and tumor mass (g) from all three independent in vivo experiments. Tumors were excised and measured postmortem. Groups include scrambled control, siU6atac (KD > 50%), siScrambled + olaparib, and siU6atac + olaparib. Each dot represents an individual tumor; red bars indicate group medians. Statistical comparisons across pooled data are shown (ns or P-values as indicated).

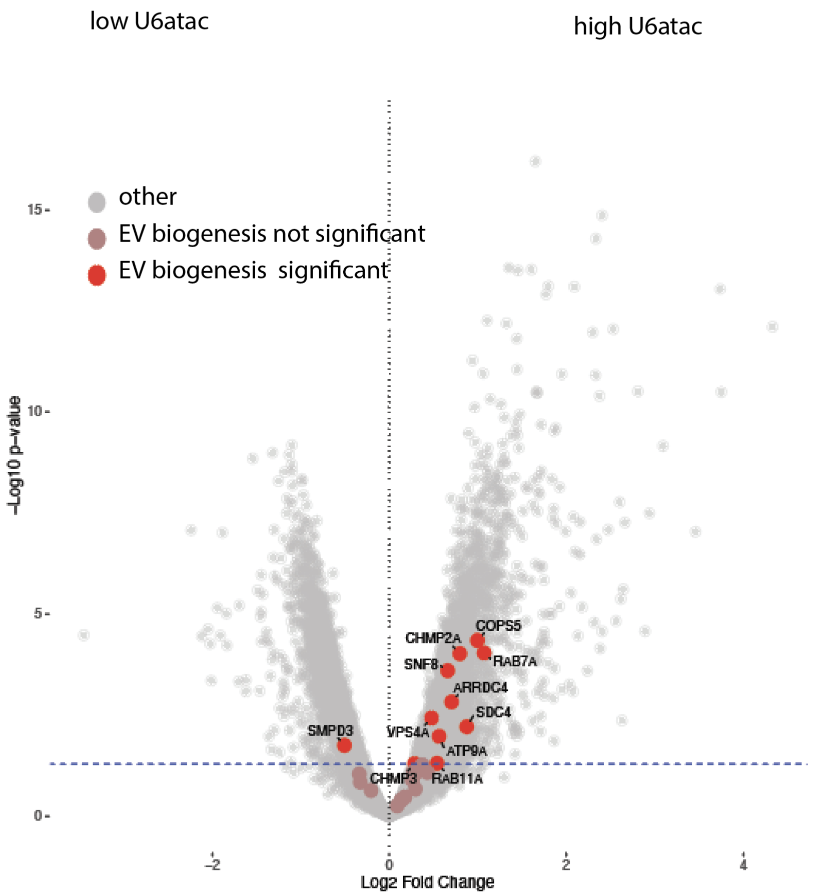


**Supplemental Figure 33 Volcano plot depicting differential gene expression between U6atac-high and U6atac-low tumors.** EV biogenesis repair–associated genes significantly enriched in U6atac-high tumors are highlighted in red, with selected key regulators of EV biogenesis labeled
